# The centromere/kinetochore is assembled through CENP-C oligomerization

**DOI:** 10.1101/2022.08.17.504347

**Authors:** Masatoshi Hara, Mariko Ariyoshi, Tomoki Sano, Ryu-suke Nozawa, Soya Shinkai, Shuichi Onami, Isabelle Jansen, Toru Hirota, Tatsuo Fukagawa

## Abstract

The kinetochore is an essential protein complex for accurate chromosome segregation. The constitutive centromere-associated network (CCAN), a subcomplex of the kinetochore, associates with centromeric chromatin providing a platform for the kinetochore assembly. A CCAN protein, CENP-C, is thought to be a central hub for the centromere/kinetochore organization. However, the crucial role of CENP-C in centromeres remains to be elucidated. Here, we demonstrated that both the CCAN-binding domain and C-terminal Cupin domain of CENP-C are necessary and sufficient for chicken CENP-C function. Our structural and biochemical analyses revealed that the Cupin domain of chicken and human CENP-C is self-oligomerization domain, which is crucial for centromeric chromatin organization. CENP-C mutants lacking the oligomerization interface cause mislocalization of CCAN and cell death. Based on these results, we conclude that the CENP-C oligomerization plays a crucial role in centromere function via providing the robust centromeric chromatin in vertebrate cells.

## Introduction

In eukaryotes, the kinetochore is a fundamental protein complex that ensures faithful chromosome segregation during mitosis and meiosis. The kinetochore is assembled on the centromere, a specialized chromatin region containing nucleosomes with the histone H3 variant CENP-A and the canonical histones (H4, H2A, and H2B) (Palmer et al., 1987; Black and Cleveland, 2011; Westhorpe and Straight, 2013). The kinetochore captures spindle microtubules and bridges the centromeric chromatin with the mitotic spindle (Fukagawa and Earnshaw, 2014; McKinley and Cheeseman, 2016; Hara and Fukagawa, 2017, 2018; Mellone and Fachinetti, 2021).

The constitutive centromere-associated network (CCAN) is a subcomplex of the kinetochore, which associates with centromeric chromatin that contains the CENP-A nucleosomes. The vertebrate CCAN, which has 16 components (CENP-C, -H, -I, -K, -L, -M, -N, -O, -P, -Q, -R, -S, -T, -U, -W, and - X) (Foltz et al., 2006; Izuta et al., 2006; Okada et al., 2006; Hori et al., 2008; Amano et al., 2009; Nishino et al., 2012), constitutively localizes to the centromere throughout the cell cycle. During late G2 and mitosis, the CCAN also provides a binding platform for another kinetochore subcomplex, the KMN (KNL1, Mis12, Ndc80 complexes) network, which directly binds to spindle microtubules (Cheeseman et al., 2006; DeLuca et al., 2006; Alushin et al., 2010; McKinley and Cheeseman, 2016; Nagpal and Fukagawa, 2016; Pesenti et al., 2016; Hara and Fukagawa, 2017). The kinetochore is an essential protein complex for chromosome segregation; therefore, it is crucial to understand how a functional kinetochore is assembled with the various proteins on the centromeric chromatin.

In recent years, extensive in vitro reconstitution and structural studies have revealed multiple protein-interaction networks in the centromere/kinetochore (Weir et al., 2016; Pesenti et al., 2018; Hinshaw and Harrison, 2019; Yan et al., 2019; Walstein et al., 2021). Most recently, the human CCAN-CENP-A nucleosome complex was reconstituted, and structures based on cryo electron microscopy (cryo-EM) results have been proposed (Pesenti et al., 2022; Yatskevich et al., 2022). These structural models provide molecular insights into the mechanisms of CCAN assembly and centromeric DNA recognition. However, these structure models exclude a large part of CENP-C, a CCAN component, due to its disordered characteristics (Pesenti et al., 2022; Yatskevich et al., 2022).

CENP-C, an essential protein for chromosome segregation and cell viability, is a conserved in most model species, from yeasts to human (Klare et al., 2015). CENP-C contains multiple functional domains and has an extended unstructured region. These functional domains bind to various proteins in the centromere/kinetochore: the KMN network through the N-terminal Mis12C-binding domain, other CCAN proteins (CENP-H-I-K-M and CENP-L-N subcomplexes: CENP-H-I-K-M/L-N) through the PEST-rich domain (or CCAN-binding domain), the CENP-A nucleosome through the central domain and CENP-C motif, and homo dimerization through the C-terminal Cupin domain (Cohen et al., 2008; Petrovic et al., 2010; Przewloka et al., 2011; Fachinetti et al., 2013; Kato et al., 2013; Basilico et al., 2014; Falk et al., 2015; Klare et al., 2015; McKinley et al., 2015; Nagpal et al., 2015; Dimitrova et al., 2016; Petrovic et al., 2016; Guo et al., 2017; Ali-Ahmad et al., 2019; Allu et al., 2019; Chik et al., 2019; Watanabe et al., 2019; Medina-Pritchard et al., 2020; Ariyoshi et al., 2021). The ability of CENP-C to bind to so many different proteins suggests that CENP-C is a hub for kinetochore assembly (Klare et al., 2015; Weir et al., 2016). The most important role of the kinetochore is bridging the centromeric chromatin with spindle microtubules. Because CENP-C binds to both the centromeric chromatin via CENP-A and the microtubule-associated KMN network, CENP-C is thought to be a key bridging protein in the kinetochore.

Unexpectedly, our previous studies revealed that neither CENP-A nor KMN interactions were necessary for CENP-C to function properly in DT40 cells (Hara et al., 2018; Watanabe et al., 2019). Instead, another CCAN protein, CENP-T, which interacts with both the KMN network and centromeric chromatin, provides the necessary and sufficient linkage between centromeric chromatin and microtubules. However, because CENP-C is an essential protein in the kinetochore assembly (Tomkiel et al., 1994; Fukagawa and Brown, 1997; Kalitsis et al., 1998; Fukagawa et al., 1999), the question remains of what crucial role(s) CENP-C plays in the centromere/kinetochore.

Here, we used a genetic complementation assay in CENP-C conditional knockout chicken DT40 cells (cKO-CENP-C) to demonstrate that the CENP-C CCAN-binding and Cupin domains are necessary and sufficient for it to function. Interestingly, our X-ray crystallographic and biochemical analyses revealed that the Cupin domain of chicken and human CENP-C forms oligomers. We identified a critical interface for the oligomerization and demonstrated that CENP-C mutants lacking this domain failed to properly recruit CCAN to the centromeres and caused growth defects. Furthermore, we found that CENP-C oligomerization is critical for maintaining centromeric chromatin organization. These comprehensive analyses indicate that the CENP-C plays a crucial role in the organization of centromeric chromatin via oligomerization to build the robust CCAN structure, which is the foundation of the kinetochore. Our results provide crucial insights into the molecular basis for centromere/kinetochore formation.

## Results

### CCAN-binding domain and Cupin domain of CENP-C are required for CENP-C function

Chicken CENP-C has multiple conserved domains that interact with other kinetochore components as well as other CENP-C to form a homodimer (Figure 1A). We previously demonstrated that the Mis12 complex (Mis12C)-binding domain (aa 1-73) and CENP-A nucleosome-binding domain (CENP-C motif, aa 653-675) are dispensable for DT40 cell viability and mitotic progression (Hara et al., 2018; Watanabe et al., 2019). Because CENP-C is known to be essential for DT40 cell viability and kinetochore function (Fukagawa and Brown, 1997; Kwon et al., 2007), a logical question is which domains of CENP-C are required for CENP-C function and kinetochore assembly.

**Figure 1.**
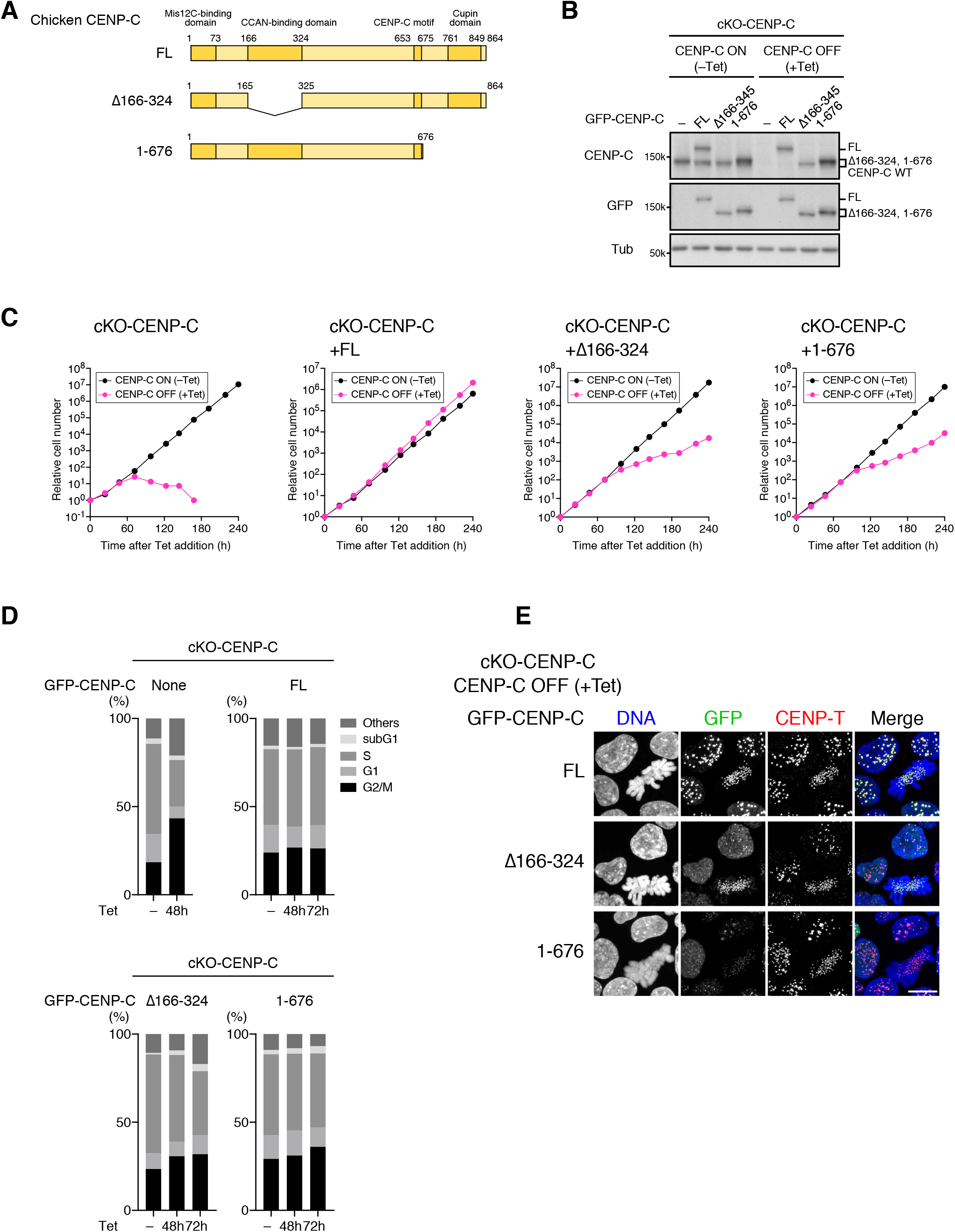
CCAN-binding and Cupin domains are required for chicken CENP-C function in DT40 cells. (A) Schematic representation of chicken CENP-C (864 amino acids: CENP-C FL, NP_001376225.2). The CCAN-binding domain (aa 166-324) was deleted in CENP-C Δ166-324. The C-terminus region, including the Cupin domain (aa 677-864), was deleted in CENP-C 1-676. (B) GFP-fused CENP-C expression in CENP-C conditional knock-out (cKO-CENP-C) chicken DT40 cells. GFP-fused CENP-C FL (FL), CENP-C Δ166-324 (Δ166-324), or CENP-C 1-676 (1-676) was expressed in cKO-CENP-C cells in which wild-type CENP-C was conditionally knocked out by tetracycline (Tet) addition. The cells were cultured with or without Tet (+Tet or –Tet) for 48 h. CENP-C proteins were examined using anti-chicken CENP-C and anti-GFP antibodies. The loading control was alpha-tubulin (Tub). cKO-CENP-C cells without GFP-CENP-C expression (–) were examined as a control. (C) Growth of cKO-CENP-C cells with or without the expression of GFP-CENP-C FL, Δ166-324, or 1-676. The cell numbers of indicated cell lines were examined at the indicated times after Tet addition (+Tet) and were normalized to those at 0 h for each line. Untreated cells were also examined (–Tet). (D) Cell cycle in cKO-CENP-C cells with or without expression of GFP-CENP-C FL, Δ166-324, or 1-676. The indicated cell lines were incubated with BrdU for 20 min at the indicated time after Tet addition and harvested (+Tet). After fixation, the cells were stained with an anti-BrdU antibody and propidium iodide. Cell cycle distribution was analyzed using flow cytometry. cKO-CENP-C cells and Tet-untreated cells in each line were also examined as controls. (E) GFP-CENP-C localization in cKO-CENP-C cells expressing GFP-CENP-C FL, Δ166-324, or 1-676. The cells were cultured as in (B), fixed, and the DNA was stained with DAPI. CENP-T was also stained with an anti-chicken CENP-T antibody and used as a centromere marker. Scale bar, 10 μm.

To address this question, we tested the remaining two domains: the CENP-H-I-K-M/L-N-binding domain (CCAN-binding domain), and the Cupin domain, which is known as a homodimerization domain in yeasts and fruit fly CENP-C homologs (Cohen et al., 2008; Chik et al., 2019; Medina-Pritchard et al., 2020) (Figure 1A). We introduced GFP-fused CENP-C, which lacks the CCAN-binding domain or the Cupin domain (GFP-CENP-C^Δ166-324^, GFP-CENP-C^1-676^) into CENP-C conditional knockout (cKO-CENP-C) DT40 cells. In these conditional knockout cells, wild-type CENP-C was expressed under the control of a tetracycline (Tet)-responsive promoter. When the cKO-CENP-C cells were treated with Tet, wild-type (WT) CENP-C expression was turned off (Figure 1B). cKO-CENP-C DT40 cells with no GFP-CENP-C expression showed growth defects and cell death by 180 h after Tet addition, mostly because of chromosome segregation errors and chromosome loss due to kinetochore malfunction (Figure 1C and 1D) (Fukagawa et al., 1999). These defects were completely suppressed by full-length CENP-C (CENP-C^FL^) expression (Figure 1C and 1D). However, the cKO-CENP-C cells expressing CENP-C^Δ166-324^ and CENP-C^1-676^ showed mild but obvious growth defects, with a slight increasement of G2/M cells, demonstrating that these domains are required for CENP-C functions (Figure 1C and 1D).

The deletion of these domains compromised centromeric localization of GFP-fused CENP-C (Figure 1E), which was consistent with the observed growth and mitotic defects. In cKO-CENP-C cells expressing GFP-CENP-C^FL^, it localized to both interphase and mitotic centromeres labelled with CENP-T. However, GFP-CENP-C^Δ166-324^ localized to centromeres only in mitotic cells, and its interphase centromeric signals were highly reduced. In addition, there was an increase in diffuse nuclear signals in the cells. Given that the aa 166-324 region binds to CENP-H-I-K-M/L-N (Klare et al., 2015; McKinley et al., 2015; Nagpal et al., 2015), the reduction in interphase centromeric localization is consistent with our previous observations that the CENP-C localization depends on CENP-H and CENP-K in interphase cells (Fukagawa et al., 2001; Kwon et al., 2007). In cKO-CENP-C cells expressing GFP-CENP-C^1-676^, GFP signals were strongly reduced in both interphase and mitotic centromeres, suggesting the importance of the Cupin domain for CENP-C centromeric localization and other functions.

### The CCAN-binding domain and Cupin domain of CENP-C are sufficient for CENP-C function

Because CENP-C has an extended unstructured domain (Suzuki et al., 2011; Suzuki et al., 2014; Klare et al., 2015), we also investigated this middle unstructured region (aa 324-653) (Figure 2A). We deleted the region together with the CENP-C motif, which was found to be dispensable for CENP-C function (Watanabe et al., 2019) (CENP-C^Δ325-676^), and expressed the resulting protein in cKO-CENP-C cells (Figure 2A and 2B). The growth of cKO-CENP-C cells expressing CENP-C^Δ325-676^ was comparable to that of cKO-CENP-C cells expressing CENP-C^FL^ in the presence of Tet, indicating that the middle unstructured region is dispensable for CENP-C function (Figure 2C).

**Figure 2.**
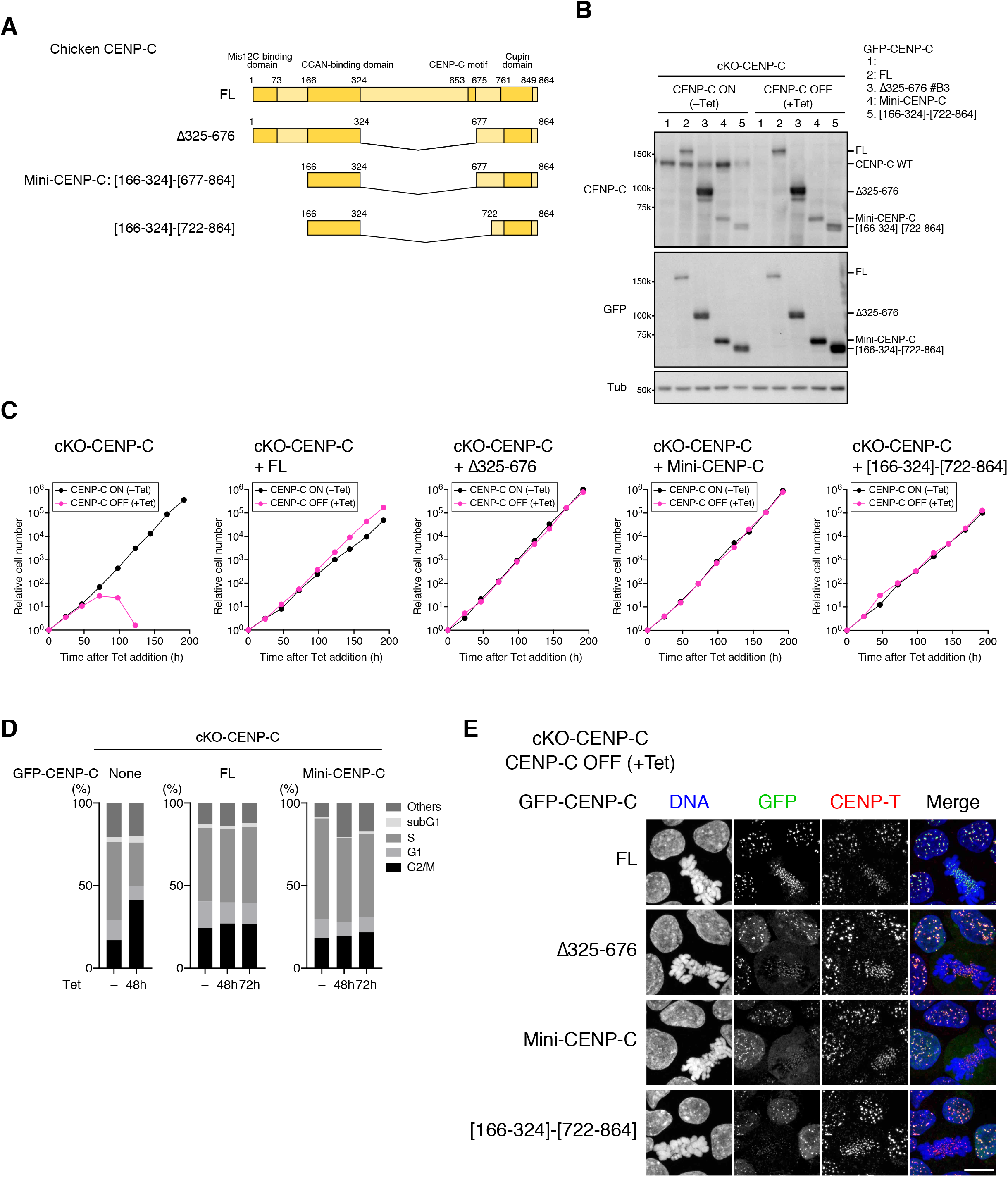
CCAN-binding and Cupin domains are sufficient for chicken CENP-C function. (A) Schematic representation of chicken CENP-C mutants. A disordered region (aa 325-656) was deleted in CENP-C Δ325-656. The CCAN-binding domain (aa 166-324) was fused with the C-terminal region, including the Cupin domain, in CENP-C [166-324]-[677-864] (Mini-CENP-C). In CENP-C [166-324]-[722-864] the shorter C-terminal region (aa 722-864) was fused with the CCAN-binding domain (aa 166-324). (B) GFP-fused CENP-C expression in cKO-CENP-C DT40 cells. GFP-fused CENP-C FL, Δ325-656, Mini-CENP-C, or [166-324]-[722-864] was expressed in cKO-CENP-C cells. The cells were cultured with or without Tet (+Tet or –Tet) for 48 h. CENP-C proteins were examined using anti-chicken CENP-C and anti-GFP antibodies. The loading control was alpha-tubulin (Tub). cKO-CENP-C cells without GFP-CENP-C expression (–) were also examined as a control. (C) Growth of cKO-CENP-C cells with or without the expression of GFP-CENP-C FL, Δ325-656, Mini-CENP-C, or [166-324]-[722-864]. The cell numbers of indicated cell lines were examined at the indicated times after Tet addition (+Tet) and were normalized to those at 0 h for each line. Untreated cells were also examined (–Tet). (D) Cell cycle in cKO-CENP-C cells with or without expression of GFP-CENP-C FL, Δ325-656, Mini-CENP-C, or [166-324]-[722-864]. The indicated cell lines were incubated with BrdU for 20 min at the indicated time after Tet addition (+Tet) and then harvested. After fixation, the cells were stained with an anti-BrdU antibody and propidium iodide. Cell cycle distribution was analyzed using flow cytometry. cKO-CENP-C cells and Tet-untreated cells in each line were also examined as controls. (E) GFP-CENP-C localization in cKO-CENP-C cells expressing GFP-CENP-C FL, Δ325-656, Mini-CENP-C, or [166-324]-[722-864]. The cells were cultured as in (B), fixed, and then their DNA was stained with DAPI. CENP-T was also stained with an anti-chicken CENP-T antibody as a centromere marker. Scale bar, 10 μm.

Because the Mis12C-binding domain is also dispensable for CENP-C function (Hara et al., 2018), we also removed the N-terminal region, including the domain from CENP-C^Δ325-676^ (CENP-C^[166-324]-[677-864]^). Strikingly, the CENP-C^[166-324]-[677-864]^, which only contained the CCAN-binding domain and a C-terminal fragment with the Cupin domain, was sufficient to complement CENP-C functions in cKO-CENP-C cells (Figure 2A-2D), and a CENP-C version with an even larger deletion in the C-terminal region (CENP-C^[166-324]-[722-864]^) also suppressed CENP-C deficiency in cKO-CENP-C cells (Figure 2A-2C). Importantly, the CCAN-binding domain and Cupin domain must be linked to hold CENP-C function, because the expression of these domains together but separately did not complement CENP-C functions in the cKO-CENP-C cells (Figure S1A-S1C), suggesting that the coupling of these two domains is important. We designated CENP-C^[166-324]-[677-864]^ as ‘Mini-CENP-C’ because of its ability to act as a minimal functional CENP-C in DT40 cells and have referred to it this way hereafter (Figure 2A).

We next examined the centromeric localization of Mini-CENP-C. In contrast to CENP-C^166-324^ or CENP-C^677-864^, neither of which localized to the centromeres in cKO-CENP-C cells (Figure S1D), Mini-CENP-C clearly localized to interphase centromeres but indistinctly to mitotic centromeres (Figure 2E). The lower centromeric localization in mitotic cells could be due to the lack of the CENP-C motif, which specifically binds to the CENP-A nucleosome during the M-phase and supports CENP-C localization to mitotic centromeres (Watanabe et al., 2019; Ariyoshi et al., 2021). This hypothesis was also supported by the localization profiles for CENP-C^Δ325-676^ and CENP-C^[166-324]-[722-864]^, which also did not localize well to mitotic centromeres (Figure 2E). These two CENP-C mutants, as well as Mini-CENP-C, but not CENP-C^FL^, showed the diffuse signals in interphase nuclei in addition to centromeric signals, suggesting that the extended disordered region could support proper centromeric localization of CENP-C. However, this disordered CENP-C region was not essential for DT40 cell growth. Based on these results, we concluded that the CCAN-binding domain and Cupin domain are necessary and sufficient for chicken CENP-C function.

### The Cupin domain of vertebrate CENP-C forms a self-oligomer

The CCAN-binding domain of CENP-C binds directly to CENP-H-I-K-M/L-N (Klare et al., 2015; McKinley et al., 2015; Nagpal et al., 2015). However, the presence of the CCAN-binding domain alone was insufficient for centromeric localization (Figure S1D), but it required the Cupin domain for the centromeric localization and function (Figure 2 and S1). Previously, the Cupin domain of yeasts and fruit fly CENP-C homologs was shown to form homodimers (Cohen et al., 2008; Chik et al., 2019; Medina-Pritchard et al., 2020). The homodimerization of CENP-C mediated by the Cupin domain is essential for the in vivo function of yeast CENP-C homologs (Cohen et al., 2008; Chik et al., 2019). The Cupin domain of vertebrate CENP-C is also thought to form homodimers, which would allow CENP-C to localize to the centromeres (Sugimoto et al., 1997; Milks et al., 2009; Walstein et al., 2021). If this were the case, the Cupin domain of vertebrate CENP-C could be interchangeable with the Cupin domain of the yeast CENP-C homolog Mif2p. However, this was not found to be the case. When we replaced the C-terminal region of Mini-CENP-C, which included the Cupin domain, with the corresponding region of Mif2p and expressed the chimeric CENP-C (CENP-C^[166-324]-[MIF2p307-549]^) in cKO-CENP-C DT40 cells (Figure S1E and S1F), the chimeric CENP-C did not complement CENP-C function despite the slight suppression (Figure S1G). This was contrast to CENP-C^FL^ or Mini-CENP-C, which did suppress the CENP-C deficiency. CENP-C^[166-324]-[Mif2p307-549]^ weakly localized to centromeres (Figure S1H), as previously shown with chimeric human CENP-C with a Mif2p Cupin domain (Walstein et al., 2021). These results indicate that the chicken CENP-C Cupin domain cannot be replaced with the Cupin domain of Mif2p. This suggests that the Cupin domain of chicken CENP-C plays an additional role in simple homodimerization.

To examine the difference between the Mif2p and chicken CENP-C Cupin domains, we determined the crystal structure of the chicken CENP-C^677-864^ at 2.8 Å resolution (Figure 3A and Table S1). The overall structure of CENP-C^677-864^ featured a classical Cupin fold consisting of ten Δ-strands (aa 761-850) with an additional structural region (aa 722-760), which was termed pre-Cupin (Figure 3A, 3B, and S2A). Both the extreme N- and C-terminal regions (aa 677-721 and aa 851-864, respectively) were disordered. The Cupin fold of chicken CENP-C indeed formed a stable homodimer interface, similar to that observed in the structures of the CENP-C Cupin domain from other species (Cohen et al., 2008; Chik et al., 2019; Medina-Pritchard et al., 2020) (Figure 3B and S2B). Furthermore, pre-Cupin region provides an additional dimer interface that is unique to chicken CENP-C. The extended pre-Cupin region of one subunit was associated with the Cupin fold of another, meaning that the dimeric structure was stabilized through inter-subunit contacts (Figure 3B).

**Figure 3.**
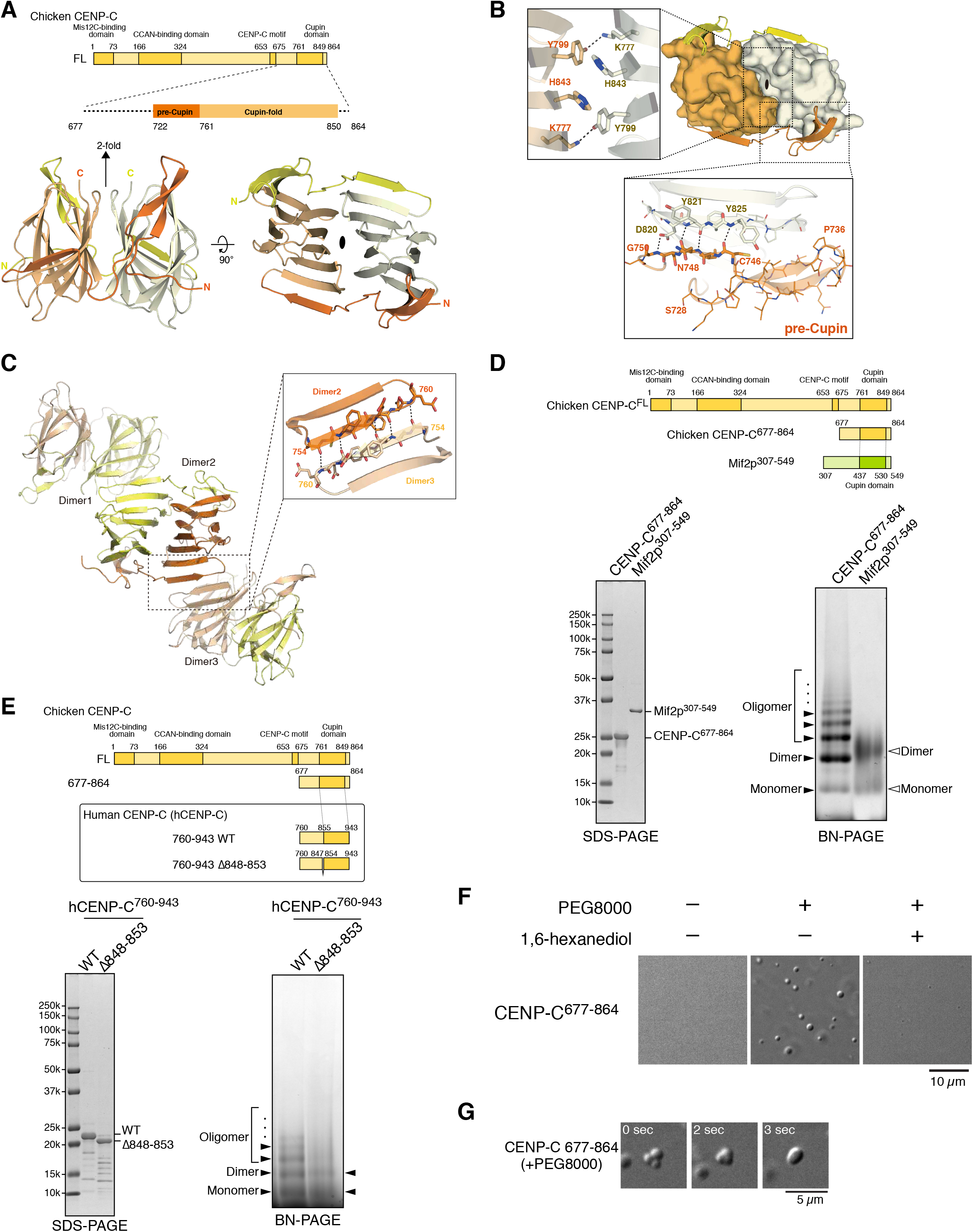
Cupin dimers of CENP-C form oligomers. (A) Crystal structure of a dimer of chicken CENP-C formed by Cupin domain. A schematic representation of the chicken CENP-C C-terminus used for structural analysis is shown on top. The Cupin monomers (colored in beige and light green) formed a dimeric structure (bottom). (B) Surface model of the Cupin fold with a 3D cartoon of the pre-Cupin region of chicken CENP-C. The insets show the major dimer interface (left) and the second interface between the pre-Cupin (orange) and Cupin fold of the next monomer (light green) (bottom). (C) A high order oligomeric structure of the Cupin dimers found in the crystal packing analyses. The Cupin dimers interacted through hydrogen bonds of the main chains (aa 754-760, inset). (D) High order oligomerization of the Cupin domain. The C-terminus of CENP-C (aa 677-864) was purified and examined using SDS-PAGE and Blue Native (BN)-PAGE. The C-terminus of Mif2p (aa 307-549) was also examined. (E) High order oligomerization of the hCENP-C Cupin domain. The C-terminus of hCENP-C (aa 760-943) wild-type (WT) and its deletion mutant, which lacks the amino acids corresponding to the oligomerization interface (Δ848-853) were purified and examined using SDS-PAGE and Blue Native (BN)-PAGE. (F) In vitro coacervate formation of the C-terminus of chicken CENP-C (aa 677-864). CENP-C 677-864 was diluted into buffer with or without 5% PEG8000 and dropped onto a coverslip. Coacervates in the drop were examined using differential interference contrast microscopy. Coacervate formation was prevented by 10% 1,6-hexanediol. Scale bar, 10 μm. (G) Fusion of CENP-C 677-864 coacervates. The coacervates were formed as in (F) and examined at the indicated time points. Time = 0 for the start of acquisition. Scale bar, 5 μm.

In addition to the homodimerization of the chicken Cupin domain, the crystal packing analysis of chicken CENP-C^677-864^ suggested the presence of further self-oligomerization state mediated by Δ-sheet formation (Figure 3C). Of note, this putative oligomer interface was formed by the main chain of the Δ3 strand in the pre-Cupin region of chicken CENP-C (aa 754-760). However, the Mif2p Cupin fold lacks the equivalent Δ-strand (Figure S2).

Because it was possible that this was a crystal packing artifact, we also examined the oligomerization of chicken CENP-C^677-864^ in solution using blue native polyacrylamide gel electrophoresis (BN-PAGE). Chicken CENP-C^677-864^ showed a ladder of bands on the BN-PAGE, whereas the Mif2p Cupin domain (aa 365-530) was observed as a monomer or dimer (Figure 3D), further supporting the conclusion that the chicken Cupin domain forms oligomers. In addition, we examined the human CENP-C (hCENP-C) fragment (aa 760-943), which is the human equivalent to chicken CENP-C^677-864^, and found that hCENP-C^760-943^ also showed a ladder profile (Figure 3E). The oligomerization of hCENP-C was previously suggested by an experiment that used crosslinked samples (Trazzi et al., 2009). To assess the oligomer interface that was identified based on the crystal structure, we deleted the Δ3 strand (aa 846-851) from hCENP-C^760-943^ and analyzed it using the BN-PAGE. The mutant Cupin domain did not show a ladder profile, indicating that hCENP-C was oligomerized through the Δ3 strand (Figure 3E).

In the presence of PEG8000, chicken CENP-C^677-864^ formed droplets that showed liquid-like properties, such as a spherical shape and fusion (Figure 3F and 3G). This also supports that CENP-C^677-864^ has a self-oligomerization activity and the hydrophobic interactions were interfered with 1-6 hexandiol that inhibits weak hydrophobic protein-protein interaction (Lin et al., 2016). These *in vitro* data suggest that the Cupin homodimers make a higher-order structure and that their structural characteristics are conserved among vertebrate CENP-C.

### Oligomerization of CENP-C through the Cupin domain is critical for CENP-C functions

To examine the importance of CENP-C oligomerization through the Cupin domain to CENP-C function, we first evaluated a dimer formation. As described above, the chicken Cupin domain has two dimer interfaces: the classical Cupin fold interface and pre-Cupin region (Figure 3B and S2). To evaluate the importance of the structure-guided dimer interfaces, we introduced mutations into these interfaces, which were expected to disrupt the dimer interactions: the classical Cupin fold (Y799A/H843A), the pre-Cupin region (Δ725-753), or both (Δ725-753/Y799A/H843A) (Figure 3B and S3A). Each CENP-C mutant was expressed in cKO-CENP-C cells to determine whether they could complement CENP-C function (Figure S3B). The growth of cKO-CENP-C cells expressing either CENP-C^FL^, CENP-C^F799A/H843^, or CENP-C^Δ725-753^ was comparable, whereas cKO-CENP-C cells expressing CENP-C^Δ725-753/F799A/H843^ exhibited growth retardation (Figure S3C). This suggests that Cupin domain dimerization is required for CENP-C function and that either the classical Cupin fold interface or the pre-Cupin region is sufficient to support Cupin domain dimerization. Consistent with this, CENP-C^Δ725-753^ and CENP-C^F799A/H843^, but not CENP-C^Δ725-753/F799A/H843^, were localized to centromeres (Figure S3D).

To examine the biological significance of CENP-C oligomerization, we introduced GFP-fused CENP-C that lacked the oligomerization interface (CENP-C^Δ754-760^) into the cKO-CENP-C cells (Figure 4A and 4B). Expression of CENP-C^FL^, but not CENP-C^Δ754-760^, suppressed the cell growth defects and cell death, with G2/M-phase enrichment in cKO-CENP-C cells (Figure 4C and 4D), indicating that the oligomerization interface is crucial for CENP-C function.

**Figure 4.**
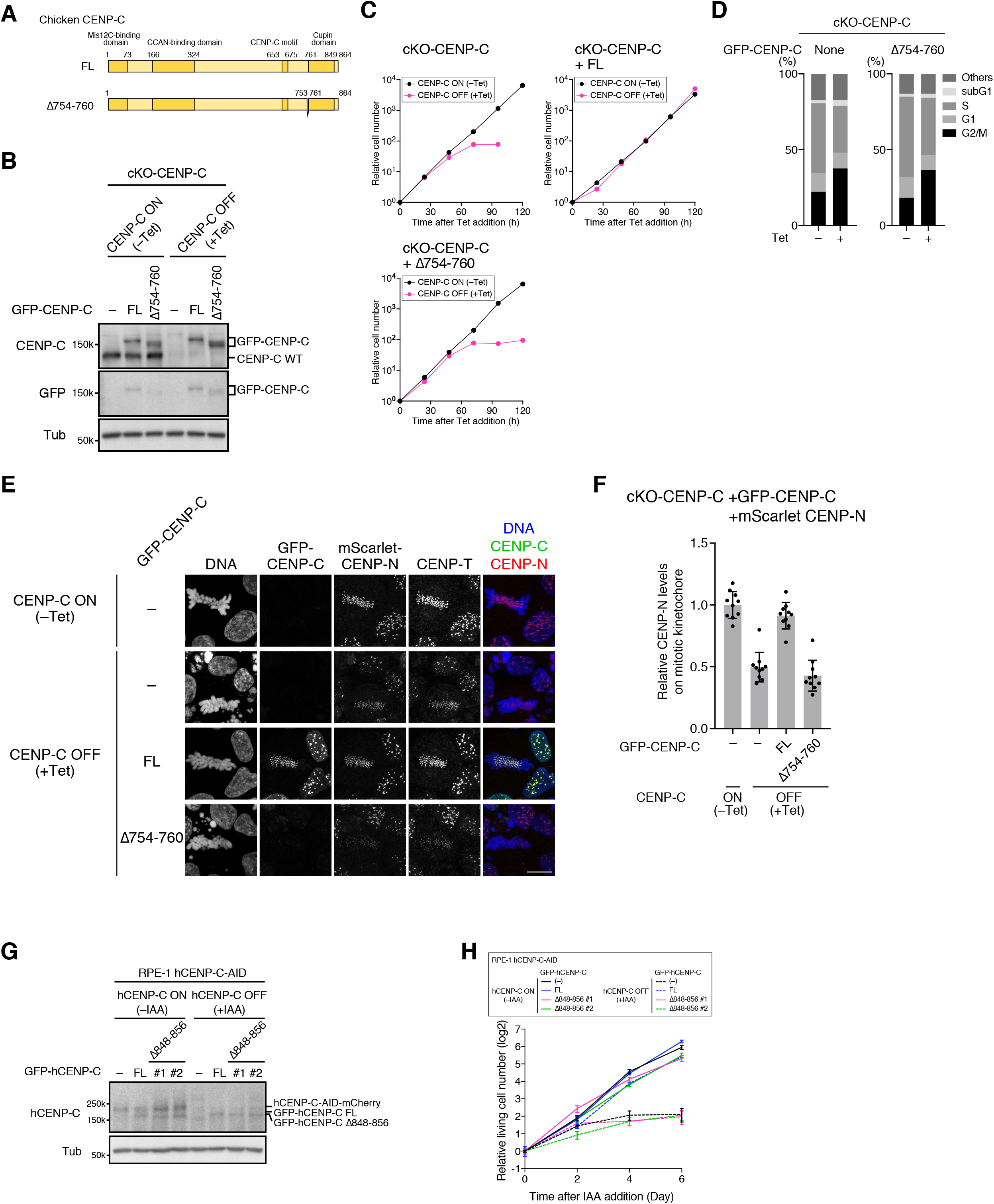
The oligomer interface is required for CENP-C function in chicken DT40 cells and human RPE-1 cells. (A) Schematic representation of the chicken CENP-C mutant. The oligomer interface (aa 754-760) was deleted in CENP-C Δ754-760. (B) GFP-fused CENP-C expression in cKO-CENP-C DT40 cells. GFP-fused CENP-C FL or Δ754-760 was expressed in cKO-CENP-C cells. The cells were cultured with or without Tet (+Tet or –Tet) for 48 h. CENP-C proteins were examined using anti-chicken CENP-C and anti-GFP antibodies. The loading control was alpha-tubulin (Tub). cKO-CENP-C cells without GFP-CENP-C expression (–) were also examined as a control. (C) Growth of cKO-CENP-C cells with or without expression of GFP-CENP-C FL or Δ754-760. The cell numbers of the indicated cell lines were examined at the indicated times after Tet addition (+Tet) and were normalized to those at 0 h for each line. Untreated cells were also examined (–Tet). (D) Cell cycle distribution in cKO-CENP-C cells with or without the expression of GFP-CENP-C FL or Δ754-760. The indicated cell lines were incubated with BrdU for 20 min at 48 h after Tet addition and harvested (+Tet). Following fixation, the cells were stained with an anti-BrdU antibody and propidium iodide. Cell cycle distribution was analyzed by flow cytometry. cKO-CENP-C cells and Tet-untreated cells in each line were also examined as controls. (E) Localization of GFP-CENP-C and mScarlet-CENP-N in cKO-CENP-C cells expressing GFP-CENP-C FL or Δ754-760. GFP-CENP-C FL or Δ754-760 was expressed in cKO-CENP-C in which endogenous CENP-N was replaced with mScarlet-CENP-N. The cells were cultured as in (B), fixed, and then their DNA was stained using DAPI. CENP-T was also stained with an anti-chicken CENP-T antibody as a centromere marker. Scale bar, 10 μm. (F) Quantification of mScarlet-CENP-N signals on mitotic centromeres. mScarlet-CENP-N signals on centromeres in mitotic cells in (E) were quantified. (Unpaired t-test, two-tailed, n = 10; – (–Tet) vs. – (+Tet): p<0.0001, – (+Tet) vs. FL (+Tet): p<0.0001, – (+Tet) vs. Δ754-760 (+Tet): ns (p = 0.2222)). (G) GFP-fused human CENP-C (hCENP-C) expression in RPE-1 hCENP-C-AID cells. GFP-fused hCENP-C FL or Δ848-856 (equivalent to Δ754-760 of chicken CENP-C) was expressed in RPE-1 hCENP-C-AID-mCherry (hCENP-C-AID) cells, in which endogenous hCENP-C was replaced with hCENP-C-AID that was depleted by IAA treatment. The cells were cultured with or without IAA (+IAA or –IAA) for 2 h. GFP-hCENP-C proteins were examined using anti-hCENP-C and anti-GFP antibodies. Two independent lines of RPE-1 hCENP-C-AID expressing GFP-hCENP-C Δ848-856 (#1, #2) were examined. The loading control was alpha-tubulin (Tub). RPE-1 hCENP-C-AID cells without GFP-hCENP-C expression (–) were examined as a control. (H) Growth of RPE-1 hCENP-C-AID cells with or without the expression of GFP-CENP-C FL or Δ848-856. The cell numbers of the indicated cell lines were examined at the indicated times after IAA addition (+IAA) and were normalized to those at 0 h for each line. Untreated cells were also examined (–IAA). Two independent lines of RPE-1 hCENP-C-AID expressing GFP-hCENP-C Δ848-856 (#1, #2) were examined.

We then examined whether CENP-C oligomerization was involved in centromere localization. Although protein levels of GFP-CENP-C^Δ754-760^ were comparable to those of GFP-CENP-C^FL^ in cKO-CENP-C cells (+Tet, Figure 4B), the signals of GFP-CENP-C^Δ754-760^ on centromeres were not detectable in both interphase and mitotic cells, whereas GFP-CENP-C^FL^ did localize to centromeres (Figure 4E). To test CCAN assembly, we introduced mScarlet-fused CENP-N, a CCAN component, into endogenous CENP-N alleles in cKO-CENP-C and quantified mScarlet signals on the mitotic centromeres (Figure 4E, 4F, and S3E). The mScarlet-CENP-N signals on mitotic centromeres were significantly reduced in cKO-CENP-C without GFP-CENP-C expression after Tet addition (GFP-CENP-C [–]; Figure4E, 4F, and S3E). The mScarlet-CENP-N signals were restored by GFP-CENP-C^FL^, but not by GFP-CENP-C^ϕλ754-760^ expression (Figure4E, 4F, and S3E). We also observed that neither GFP-CENP-C^166-324^ nor GFP-CENP-C^1-676^ supported the recovery of mScarlet-CENP-N levels at mitotic centromeres in cKO-CENP-C cells (Figure S4A-D). In contrast, Mini-CENP-C supported a significant recovery of mScarlet-CENP-N levels at mitotic centromeres (Figure S4A-D). Taken together, CENP-C oligomerization via the Cupin domain was required for the proper centromeric localization of CENP-N and for CENP-C to build kinetochores on centromeres in chicken DT40 cells.

Next, we tested whether oligomerization was also essential for hCENP-C function using an auxin-inducible degron (AID)-based system in RPE1 cells (RPE-1 CENP-C-AID cells) (Watanabe et al., 2019). We expressed GFP-fused hCENP-C that lacked the oligomerization interface (hCENP-C^ϕλ846-851^) in RPE-1 CENP-C-AID cells and then examined cell proliferation in two independent cell lines (Figure 4G and 4H). After the addition of indole-3-acetic acid (IAA), RPE-1 CENP-C-AID cells without GFP-hCENP-C stopped growing (Figure 4H). This cell proliferation defect was restored by GFP-hCENP-C^FL^, but not GFP-hCENP-C^ϕλ846-851^, expression, indicating that oligomerization is required for hCENP-C function (Figure 4G and 4H).

### Artificial oligomerization of CENP-C facilitates CCAN centromeric localization

To further investigate the importance of CENP-C oligomerization, we used an artificially inducible oligomerization system to oligomerize a CENP-C fragment without the Cupin domain. In this system, an engineered FKBP (DmrB), which forms a self-dimer upon addition of a small chemical dimerizer (B/B Homodimerizer), is tandemly repeated to allow the formation of oligomers following the addition of the dimerizer (Figure S4E and S4F) (Clackson et al., 1998; Takamatsu et al., 2013). Indeed, when GFP-fused single DmrB (1XDmrB) or three tandem repeats of DmrB (3XDmrB) were expressed in DT40 cells, GFP-3XDmrB, but not GFP-1XDmrB, formed molecular condensates in cells treated with the dimerizer. This indicated that the artificial oligomerization of 3XDmrB had occurred in the DT40 cells (Figure S4G).

Next, we swapped the Cupin domain of chicken Mini-CENP-C with either 1XDmrB or 3XDmrB (CENP-C^[166-324]-[677-724]-1XDmrB^ or CENP-C^[166-324]-[677-724]-3XDmrB)^ (Figure 5A). We expressed either of them or CENP-C^166-324^ in cKO-CENP-C cells and examined their proliferation (Figure 5A-5C). As described above (Figures S1C), the cKO-CENP-C cells and the cells expressing GFP-CENP-C^166-324^ showed cell proliferation defects after Tet addition (Figure 5C). The expression of CENP-C^[166-324]-[677-724]-1XDmrB^ restored neither the growth defects nor G2/M enrichment in cKO-CENP-C cells after Tet addition even with the dimerizer (Figure 5C and 5D). In contrast, the cell proliferation defects were partially but substantially suppressed by CENP-C^[166-324]-[677-724]-3XDmrB^ expression in the presence of the dimerizer (Figure 5C and 5D).

**Figure 5.**
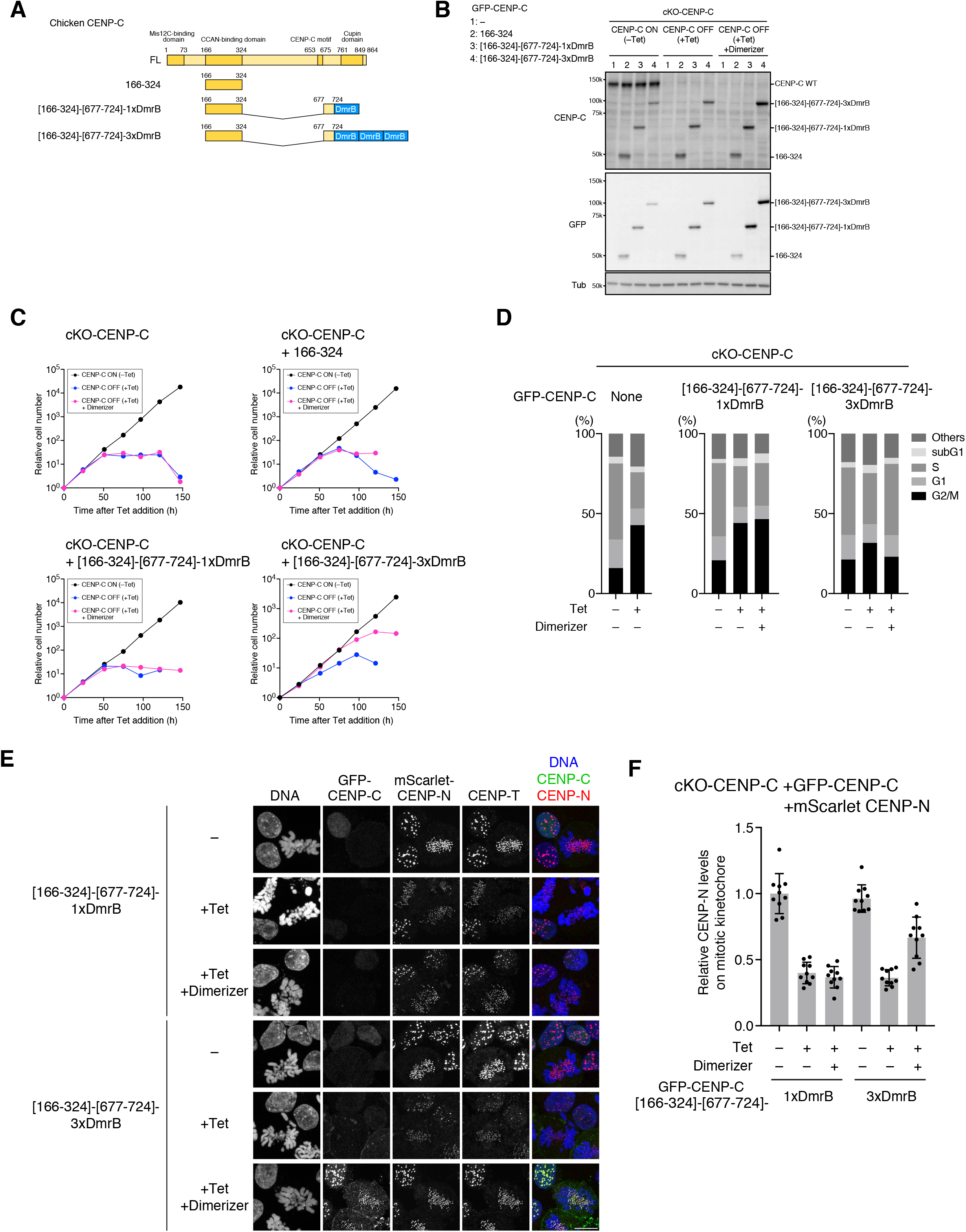
Oligomerization of CENP-C facilitates centromere localization of CENP-C and CCAN assembly. (A) Schematic representation of the chicken CENP-C mutant. The CCAN-binding domain (166-324) was fused with the CENP-C C-terminus in which the oligomerization domain was replaced with a single DmrB domain, an artificial inducible homodimerization domain ([166-324]-[677-724]-1xDmrB). For inducible oligomerization, three tandem-repeated DmrB was fused with [166-324]-[677-724] ([166-324]-[677-724]-3xDmrB). (B) GFP-fused CENP-C expression in cKO-CENP-C DT40 cells. GFP-fused CENP-C 166-324, [166-324]-[677-724]-1xDmrB, or [166-324]-[677-724]-3xDmrB was expressed in cKO-CENP-C cells. The cells were treated with no chemical (–Tet), Tet (+Tet), or both Tet and B/B Homodimerizer (+Tet, +Dimerizer) for 48 h. CENP-C proteins were examined with anti-chicken CENP-C and anti-GFP antibodies. The loading control was alpha-tubulin (Tub). cKO-CENP-C cells without GFP-CENP-C expression (–) were also examined as a control. (C) Growth of cKO-CENP-C cells with or without expression GFP-fused CENP-C 166-324, [166-324]-[677-724]-1xDmrB, or [166-324]-[677-724]-3xDmrB. The cell numbers of indicated cell lines were examined at the indicated times after Tet addition (+Tet) with or without dimerizer and were normalized to those at 0 h for each line. Untreated cells were also examined (–Tet). (D) Cell cycle distribution in cKO-CENP-C cells with or without expression of GFP-fused CENP-C 166-324, [166-324]-[677-724]-1xDmrB, or [166-324]-[677-724]-3xDmrB. The indicated cell lines were incubated with BrdU for 20 min at 48 h after Tet addition (+Tet) with or without dimerizer and then harvested. After fixation, the cells were stained with an anti-BrdU antibody and propidium iodide. Cell cycle distribution was analyzed by flow cytometry. cKO-CENP-C cells and Tet-untreated cells in each line were also examined as controls. (E) GFP-CENP-C and mScarlet-CENP-N localization in cKO-CENP-C cells expressing GFP-CENP-C 166-324, [166-324]-[677-724]-1xDmrB, or [166-324]-[677-724]-3xDmrB. The cells were cultured as in (B), fixed, and then their DNA was stained using DAPI. CENP-T was also stained with an anti-chicken CENP-T antibody as a centromere marker. Scale bar, 10 μm. (F) Quantification of mScarlet-CENP-N signals on mitotic centromeres. mScarlet-CENP-N signals on centromeres in mitotic cells in (E) were quantified. (Unpaired t-test, two-tailed, n = 10; 1xDmrB +Tet/–Dimerizer vs. +Tet/+Dimerizer: ns (p = 0.381), 3xDmrB +Tet/–Dimerizer vs. +Tet/+Dimerizer: p<0.0001).

These cell growth results were consistent with the localization of CENP-C fragments fused with 1x or 3xDmrB. CENP-C^[166-324]-[677-724]-1XDmrB^ did not localize to centromeres, even after the addition of the dimerizer (Figure 5E). However, GFP-CENP-C^[166-324]-[677-724]-3XDmrB^ did clearly localize to centromeres after the addition of the dimerizer, whereas its centromeric localization was not detected in the absence of the dimerizer (Figure 5E). These data suggest that oligomerization of the CCAN-binding domain is necessary and sufficient for centromeric localization. In addition to CENP-C fragment localization, artificial oligomerization of the CCAN-binding domain through 3XDmrB also restored CENP-N levels on mitotic centromeres, which were reduced in cKO-CENP-C cells after Tet addition (Figure 5E, 5F, and S4H). These data strongly suggest that oligomerization of CENP-C facilitates its own kinetochore localization, leading to CCAN assembly.

### CENP-C depletion changed CENP-A cluster organization in centromeres

Based on in vitro reconstitution and structural studies, it has been speculated that CENP-C folds centromeric chromatin that contains CENP-A nucleosomes into spatially organized structures (Ali-Ahmad et al., 2019; Allu et al., 2019). This idea is also supported by the centromere chromatin fiber assay, which suggests that CENP-C stabilizes centromeric chromatin (Ribeiro et al., 2010).

To quantitatively examine the organization of centromeric chromatin in cells, we used a single molecular localization microscope (SMLM), STochastic Optical Reconstruction Microscope (STORM), to analyze the distribution of CENP-A molecules in the centromeres. We stained CENP-A in RPE-1 CENP-C-AID cells with an anti-CENP-A antibody and inspected CENP-A signals in mitotic cells with high Cyclin B1 signals. Under a conventional fluorescence microscopy, CENP-A molecules labeled with the antibodies at the centromere were observed as a single dot due to diffraction limitations (Figure 6A). In the 2D-STORM images, multiple fluorophores labeling CENP-A molecules were distributed in each centromere and revealed the elongated round shape of the cluster of CENP-A nucleosomes in a mitotic centromere (Figure 6A and 6B), which is consistent with a previous report (Andronov et al., 2019). We then measured CENP-A clusters of each centromere using the pair correlation function (Figure S5A; see Methods). The average of calculated radial distance of CENP-A clusters in the control cells was 229±5.11 nm, which was comparable to the CENP-A chromatin length observed using immuno-electron microscopy (Marshall et al., 2008) (hCENP-C ON, Figure 6C and S5B). CENP-A cluster size was significantly reduced in CENP-C-depleted RPE-1 cells (Figure 6A-C and S5B).

**Figure 6.**
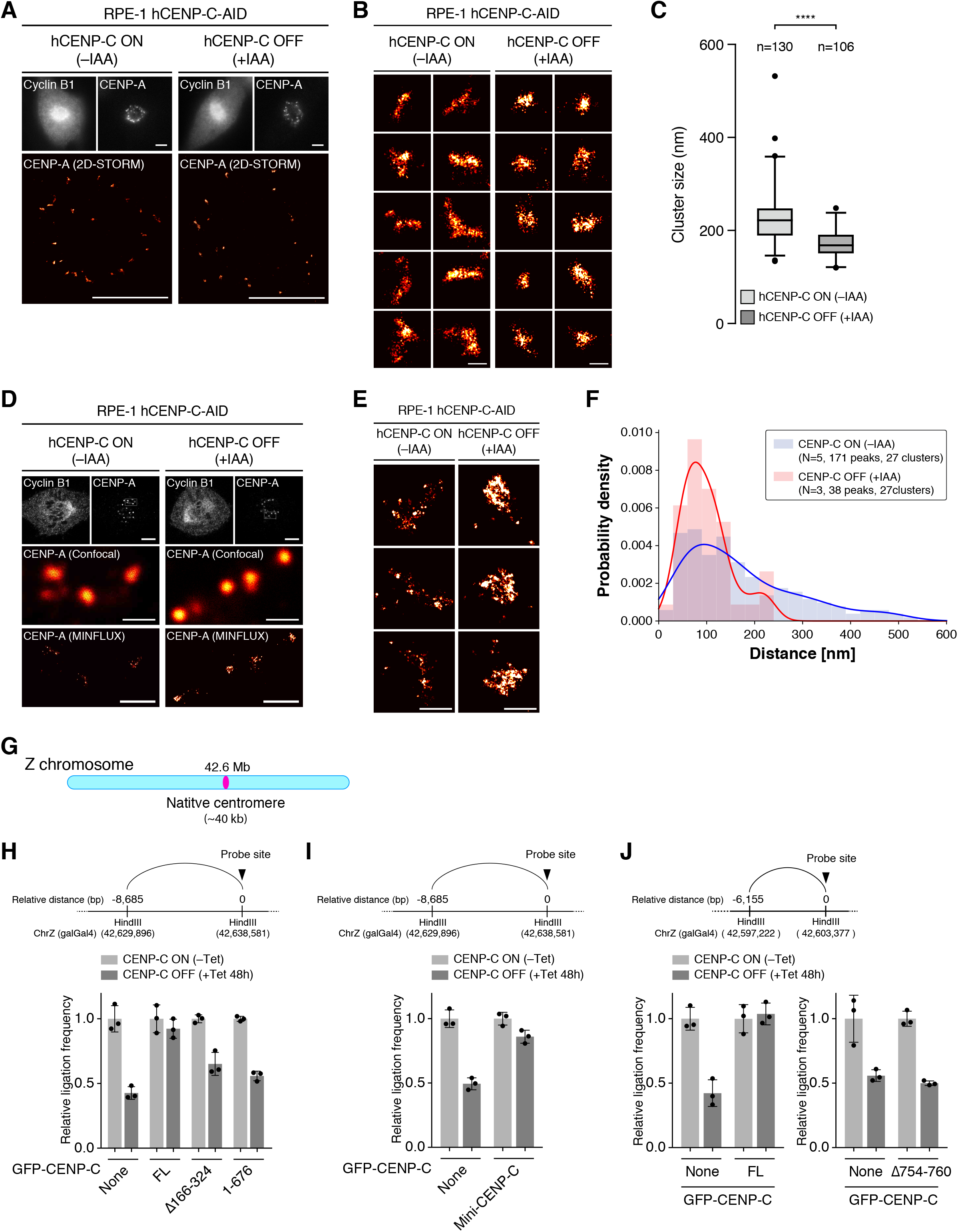
The CENP-C oligomer interface is required for centromeric chromatin formation. (A) 2D-STORM images of CENP-A in RPE-1 cells. RPE hCENP-C-AID cells were fixed at 2 h after treatment with or without IAA. CENP-A was stained with an anti-human CENP-A antibody. Cyclin B1 was also stained with an anti-human Cyclin B1 antibody as a mitotic cell marker. Upper panels are widefield microscopy images. Bottom panels are overviews of the STORM results. Scale bar, 5 μm. (B and C) CENP-A clusters rendered using 2D-STORM analysis. (B) The panels show CENP-A clusters in STORM images in (A). (C) The characteristic length of each cluster was analyzed using the normalized pair correlation function (Figure S5A and S5B; see also Methods). The CENP-A cluster size is given by the value of *r* at the position intersecting the line of random distribution (*g*(*r*) = 1). The graph shows the average of clusters. Scale bar, 200 nm. (D) MINFLUX images of CENP-A in RPA-1 cells. RPE hCENP-C-AID cells were stained as in (A). Upper panels are confocal microscopy images of mitotic cells. Scale bar, 5 μm. Middle and bottom panels are CENP-A clusters of confocal and MINFLUX images, respectively. Scale bar, 1 μm. (E and F) CENP-A clusters as determined by MINFLUX analysis. (E) The panels show CENP-A clusters in MINFLUX images in (D). Scale bar, 200 nm. (F) The clusters were analyzed using the pair correlation function. The graph shows that the normalized numbers of peaks on *g(r)* appeared above *g(r)* = 1 at each distance with a 30-nm binning width (Figure S5C). The bold blue and red curves show the Gaussian kernel density estimates for CENP-C on and off states, respectively. (G) Schematic representation of the chicken Z chromosome. The chicken Z chromosome has a native centromere spanning 40 kb in the 42.6 Mb region (Hori et al., 2017; Nishimura et al., 2019). (H) 3C-qPCR analysis of centromere chromatin in cKO-CENP-C cells expressing CENP-C Δ166-324 or 1-676. cKO-CENP-C cells with or without GFP-CENP-C FL, Δ166-324, or 1-676 were fixed 48 h after treatment with or without Tet. The 3C-libraries were prepared, and the ligation frequency at the indicated sites within the centromere region was examined using TaqMan real-time PCR. The marked probe site indicates the location of the TaqMan probe site. (I) 3C-qPCR analysis of centromere chromatin in cKO-CENP-C cells expressing Mini-CENP-C. Ligation frequency at the indicated sites in cKO-CENP-C cells expressing GFP-Mini-CENP-C was analyzed by 3C-qPCR, as in (H). (J) 3C-qPCR analysis of centromere chromatin in cKO-CENP-C cells expressing CENP-C Δ754-760. cKO-CENP-C with or without expression of GFP-CENP-C FL or Δ754-760 was fixed 48 h after treatment with or without Tet. The 3C-libraries were prepared, and the ligation frequency at the indicated sites within the centromere region was examined using TaqMan real-time PCR. The marked probe site indicates the TaqMan probe site.

We also used another SMLM, minimal photon flux (MINFLUX) nanoscopy, which enabled us to detect fluorophores with a resolution of several nanometers (Balzarotti et al., 2017; Schmidt et al., 2021). MINFLUX rendered elongated CENP-A clusters, as observed by STORM, with higher resolution and robustness (Figure 6D and 6E). Within the clusters, several small groups of labeled CENP-A molecules were observed in control cells (hCENP-C ON, Figure 6E). Pair correlation analysis depicted the distances between CENP-A groups within a cluster as peaks on the curve of the graph (Figure S5C). The peaks appeared at around 100-400 nm, indicating that the CENP-A groups were distributed with 100-400 nm distances in a cluster (Figure 6F and S5C). In the CENP-C-depleted RPE-1 cells (hCENP-C OFF), CENP-A clusters shrank into roundish shapes with packed CENP-A groups, resulting in peaks that were less clear with a distance of 100 nm in pair correlation analysis (Figure 6D and S5C). Together with the STORM analysis, these observations suggest that CENP-C is required for the maintenance of CENP-A cluster organization in centromeric chromatin.

### The centromere forms self-associated chromatin through CENP-C oligomerization

Centromeric chromatin, which includes the CENP-A nucleosomes and H3 nucleosomes with specific histone modifications, has been shown to be held in unique three-dimensional structures (Sullivan et al., 2001; Blower et al., 2002; Sullivan and Karpen, 2004; Ribeiro et al., 2010; Fukagawa and Earnshaw, 2014). We previously demonstrated that centromeric chromatin is self-associated, using 4C-seq analysis of neocentromeres in DT40 cells (Nishimura et al., 2019). It has also been suggested that CCAN proteins are involved in centromeric chromatin formation. Indeed, our STORM and MINFLUX results found that CENP-C is required for the maintenance of CENP-A cluster organization, suggesting that CENP-C plays a key role in maintaining the centromeric chromatin structure. To test this hypothesis, we quantitatively assessed the chromatin structure within centromeres at the sequence level by applying a TaqMan probe-based 3C-qPCR method (Hagege et al., 2007) to a DT40 cell line containing a neocentromere at the 35 Mb region on the Z chromosome (Neo-Centromere cell) (Shang et al., 2013; Nishimura et al., 2019). (Figure S6A).

For 3C-qPCR in the neocentromere region, we prepared 3C libraries with HindIII from Neo-centromere cells, and also wild-type cells, which have a centromere at the original locus (the 42 Mb region) on the Z chromosome (Chr.Z) as a control (Control cells). We designed a TaqMan probe at the HindIII site in the 35 Mb region, which is shown as the probe site in Figure S6A (relative distance: 0 bp), and examined the interaction between the probe site and each HindIII site several thousand base pairs away from the probe sites within the neocentromere. We used a ligation frequency to determine the interaction (Figure S6A). The values of ligation frequency in the 35 Mb region were in the Neo-centromere cells much higher than the Control cells, suggesting that long-range interactions (2-15 kb in length) frequently occurred in the neocentromere (Figure S6A and S6B). These results demonstrate that our 3C-qPCR method worked well in detecting unique self-associated chromatin characteristics in the neocentromere.

We then examined whether CENP-C was involved in chromatin organization in the neocentromere, using Neo-centromere cells in which endogenous CENP-C was replaced with AID-tagged CENP-C (Nishimura et al., 2019). The AID-tagged CENP-C was rapidly depleted within two hours of IAA treatment, without changes in cell cycle distribution (Figure S6C and S6D). In CENP-C-depleted Neo-centromere cells (+IAA), the ligation frequencies of all tested sites in the neocentromere were reduced compared with those in the cells without IAA treatment (–IAA). Curiously, the SMLM data suggested that CENP-A clusters on centromeric chromatin shrank after CENP-C depletion (Figure S5). This appears to contradict the 3C-qPCR results. However, considering that the ligation frequency is dependent not only on the three-dimensional distance between two DNA fragments but also on the chromatin flexibility or dynamics, the shrunken CENP-A clusters in CENP-C-depleted cells could be stiff and less dynamic, which could reduce the ligation frequency in 3C-qPCR (see also Discussion). We emphasize that CENP-C is required for the maintenance of the proper chromatin organization in centromeres.

We also observed a reduction of ligation frequencies in the neocentromeres of CENP-H-depleted cells (Nishimura et al., 2019), although it was less than that observed after CENP-C depletion (Figure S6E). Even though CENP-H was rapidly degraded, cells in the G2/M phase were slightly increased two hours after IAA addition (Figure S6D). However, the G2/M increase is unlikely to be a reason for the lower ligation frequencies after CENP-H depletion, because nocodazole treatment, which induced the same extent of G2/M increase caused by CENP-H depletion (e.g., 2 h) did not reduce the ligation frequencies compared to CENP-H removal, although prolonged nocodazole treatment decreased the ligation frequency (Figure S6F and S6G). These data suggest that CENP-H is involved in the formation of a specific chromatin structure in the neocentromere but to a lesser extent than CENP-C.

Chicken Chr.Z also contains a non-repetitive centromere at the 42 Mb region (Figure 6G) (Shang et al., 2010; Shang et al., 2013; Hori et al., 2017; Nishimura et al., 2019). Taking advantage of this unique feature, we investigated whether CENP-C was involved in the chromatin organization of the native centromere (Figure S7A). 3C-qPCR revealed that the ligation frequencies in the 42 Mb region were reduced in cKO-CENP-C cells treated with Tet and that the reduced values were restored by GFP-CENP-C^FL^ expression (Figure 6H, S7B, and S7C). Importantly, although Tet treatment increased the G2/M phase population in cKO-CENP-C cells, this was not the reason for the reduction of ligation frequencies. Because the 4 h nocodazole-treatment increased the G2/M population but did not largely change the ligation frequencies (Figure S7C and S7D), we concluded that the reduction of ligation efficiency in cKO-CENP-C after Tet-treatment was due to CENP-C deficiency but not an effect of G2/M-cell enrichment.

Because the CCAN-binding domain and Cupin domain are essential for CENP-C function (Figure 1), we examined whether these domains were required for centromeric chromatin formation. We analyzed centromeric chromatin on Chr.Z in cKO-CENP-C cells expressing GFP-CENP-C^Δ166-324^ or - CENP-C^1-676^ using 3C-qPCR. Neither one restored the ligation frequencies reduced by CENP-C knockout (Figure 6H, S7B, and S7C). In contrast, Mini-CENP-C expression largely suppressed the reduction of ligation frequencies caused by CENP-C knockout. These results suggest that CCAN-binding and Cupin domains are necessary and sufficient for the CENP-C-dependent centromeric chromatin organization in the native centromere on Chr.Z (Figure 6I, S7B, and S7E).

Finally, we analyzed the role of CENP-C oligomerization in centromeric chromatin organization. We expressed GFP-CENP-C^Δ754-760^ in cKO-CENP-C cells and assessed centromeric chromatin using the 3C-qPCR. Expression of CENP-C^Δ754-760^, unlike CENP-C^FL^, did not restore the ligation frequency reduced in cKO-CENP-C cells after Tet addition (Figure 6J, S7B, and S7F-S7H), suggesting that CENP-C oligomerization is required for the CENP-C-dependent centromeric chromatin organization.

## Discussion

In this study, we elucidated that both the CCAN-binding domain and Cupin domain are essential for CENP-C function in the centromere/kinetochore. These two domains are sufficient for CENP-C function, as we successfully established a functionally Mini-CENP-C, which contained only the CCAN-binding and Cupin domains. Interestingly, the Cupin homodimers of vertebrate CENP-C self-associate into high-order oligomers (Figure 7A) and that oligomerization is essential for CENP-C function in maintaining the high-order structure of centromeric chromatin (Figure 7B). These findings led to a model in which CENP-C associates with other CCAN components and bundles up CCAN complexes through Cupin domain oligomerization to create self-associated unique centromeric chromatin (Figure 7B). We propose that this process plays an essential role in centromere/kinetochore function.

**Figure 7.**
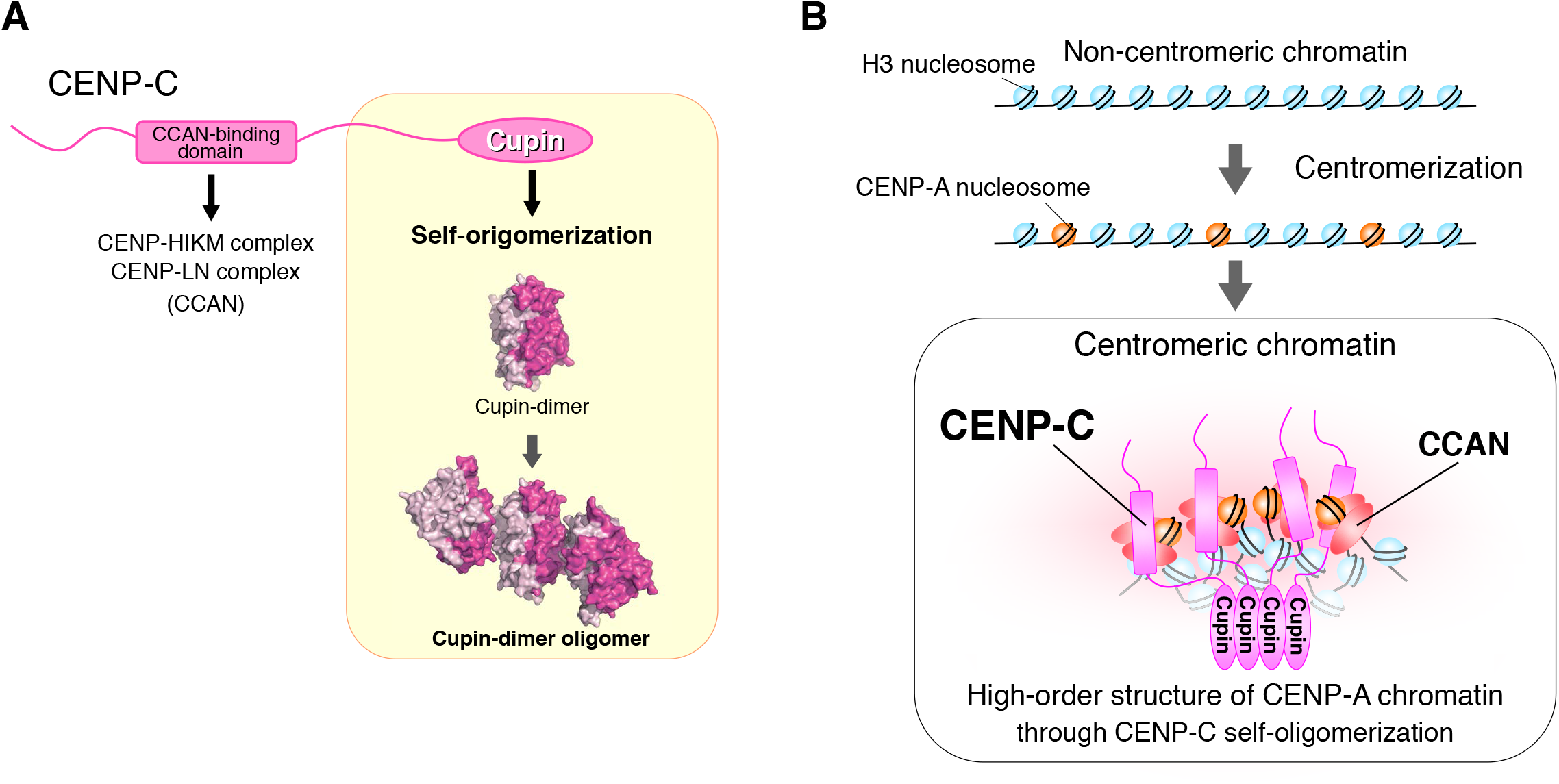
CENP-C self-oligomerization via the Cupin domain is essential for its function. (A) The essential domains for chicken CENP-C function. Two conserved domains are crucial for chicken CENP-C function: The CCAN-binding domain for binding of other CCAN subcomplexes, CENP-H-I-K-M and -L-N (Klare et al., 2015; McKinley et al., 2015; Nagpal et al., 2015), and the Cupin domain for self-oligomerization. (B) A model for establishment and maintenance of centromeric chromatin structure through oligomerization of CENP-C. A non-centromeric chromatin region, which has little or no self-interaction, would be folded into a high-order structure, when the region is centromerized through loading of CENP-A and binding of centromere/kinetochore proteins, including CENP-C. CCAN-binding and self-oligomerization of CENP-C are crucial to maintain unique centromeric chromatin structure with clusters of CENP-A nucleosomes on the centromere. The centromeric chromatin is collapsed with shrunken CENP-A clusters after CENP-C depletion.

In recent years, extensive studies using in vitro reconstitution and structural analyses of the kinetochore complex have revealed interaction networks of kinetochore proteins (Weir et al., 2016; Pesenti et al., 2018; Hinshaw and Harrison, 2019; Yan et al., 2019; Walstein et al., 2021; Pesenti et al., 2022; Yatskevich et al., 2022). These studies have greatly improved our understanding of kinetochore assembly. However, the CENP-C proteins used in the reconstituted CCAN studies lacked the Cupin domain, and a large part of the CENP-C information is missing in the human CCAN structural models (Pesenti et al., 2022; Yatskevich et al., 2022). In the present study, we demonstrated that the oligomerization of CENP-C via its Cupin domain is crucial for assembling the CCAN complex on centromeres. This finding fills in one of the key missing pieces of the current CCAN structure model and will improve our understanding of how vertebrate CCAN is built on centromeric chromatin.

Homodimerization of the Cupin domain is a conserved structural feature of CENP-C. The Cupin domain of Mif2p, a CENP-C homolog of *Saccharomyces cerevisiae* with point centromeres, forms a homodimer via the classical Cupin fold (Cohen et al., 2008). Homodimerization of the Cupin domain has also been observed in other species that possess regional centromeres, such as *Schizosaccharomyces pombe* and *Drosophila melanogaster* (Chik et al., 2019 Medina-Pritchard, 2020 #5777). The CENP-C Cupin domains of these species have an additional structural feature that the Mif2p Cupin domain does not, namely pre-Cupin region to support dimer formation, which suggests that the additional dimerization interface is a specific feature of CENP-C with the regional centromeres (Chik et al., 2019 Medina-Pritchard, 2020 #5777). The results of our structural and in vitro studies support this hypothesis. The additional dimerization interface of the CENP-C Cupin domain, pre-Cupin region, is conserved in chickens and humans, both of which have regional centromeres.

In addition, we revealed that chicken and human Cupin homodimers assemble into high-order oligomers. Because the CENP-C Cupin domains of fission yeast and fruit fly have no obvious Δ-strand, which would correspond to the oligomerization interface found in chicken and human CENP-C, the oligomerization of the Cupin domain might be a unique feature of vertebrate CENP-C. However, it would be interesting to test whether the CENP-C Cupin domain in invertebrates is oligomerized.

CENP-C oligomerization through the Cupin domain is required for CCAN recruitment and the maintenance of centromeric chromatin structures. We propose a model in which Cupin oligomerization is physically involved in the centromeric chromatin architecture (Figure 7B).

The significance of CENP-C oligomerization was further supported by an inducible artificial oligomerization system that used 3xDmrB. Oligomerization of the CCAN-binding domain through 3xDmrB largely restored the growth defects caused by CENP-C deficiency. However, this suppression was imperfect. In contrast, Mini-CENP-C, in which the CCAN-binding domain is fused with the CENP-C oligomerization domain (Cupin domain), fully suppressed CENP-C deficiency. This indicates that the Cupin oligomer has additional properties that the 3xDmrB oligomers lack. One possibility is that CENP-C Cupin oligomers could be assembled into a unique architecture, which could not be reproduced by the 3xDmrB oligomers. Another possibility is that the Cupin oligomers might be dynamic, rather than static, structures. Our previous studies have demonstrated that CENP-C changes its interaction partners on centromeres during cell cycle progression (Fukagawa et al., 2001; Kwon et al., 2007; Hara et al., 2018; Hara and Fukagawa, 2018; Watanabe et al., 2019). Cupin oligomers may undergo dynamic structural changes or assembly/disassembly during cell cycle progression, which could not be reproduced by 3xDmrB-mediated oligomerization. Alternatively, it is also possible that the Cupin domain might play a role that does not involve oligomerization. In fact, during meiosis, the Cupin domain interacts with the meiotic-specific proteins Meikin and Moa1 in mice and fission yeast, respectively (Kim et al., 2015; Chik et al., 2019; Maier et al., 2021). The CENP-C Cupin domain may also interact with other proteins during mitosis.

Several alternative 3D-structure models for centromeric chromatin have been proposed (Fukagawa and Earnshaw, 2014). We optimized the 3C-qPCR method to assess the self-interaction within the neocentromere and native centromere on chicken Chr.Z, both of which are composed of non-repetitive unique DNA sequences. Our 3C-qPCR method can quantitatively evaluate centromeric chromatin features as a ligation frequency between two distant loci (several kb away) within the centromere. After applying 3C-qPCR analysis to the neocentromere, we found clear evidence that centromerization induces the formation of a unique chromatin structure with self-interactions in a chromatin region that originally had fewer or no self-interactions (Figure 7B). Furthermore, we demonstrated that CENP-C largely contributes to the maintenance of centromere chromatin structure through self-interactions. However, although ligation probabilities within the centromeric region were reduced by CENP-C depletion, they were still largely higher than those in non-centromeric chromatin. This observation suggests that once the centromere/kinetochore structure is established, multiple factors cooperate to maintain the centromeric chromatin architecture. Consistent with this idea, a recent study has suggested that CENP-N contributes to the compaction of centromeric chromatin (Zhou et al., 2022).

We also visualized the CENP-A cluster on centromeres in human RPE-1 cells using SMLM and found that CENP-C maintained the proper shape of CENP-A clusters. CENP-C depletion shrank the clusters (Figure 6A-F). This seems to be contrary to what was expected from the 3C-qPCR analyses in DT40 cells, in which CENP-C depletion reduced ligation frequencies (Figure 6H-J). One possible explanation is that the centromeric chromatin with the shrunken CENP-A cluster in CENP-C-depleted cells is less dynamic, resulting in a reduction of crosslinking and ligation probabilities among chromatin loci during 3C-library preparation. Another explanation is that oligomerized CENP-C might counteract forces compressing centromeric chromatin, which could be mediated by CENP-N and/or heterochromatin surrounding centromeres (Larson et al., 2017; Zhou et al., 2022). CENP-C depletion might cause over-compaction via collapse of the proper centromeric chromatin. Alternatively, the difference between repetitive (alpha-satellite DNA repeats in human centromeres) and non-repetitive DNA sequences (neocentromere and centromere on chicken Chr.Z) might explain these contradictory observations. These repetitive alpha-satellite DNA sequences are known to fold into secondary structures in solution (Gallego et al., 1997; Garavis et al., 2015). CENP-C may oppose random chromatin folding through alpha-satellite DNA repeats and thus maintain proper chromatin organization in centromeres. CENP-C depletion might induce secondary structures in alpha-satellite DNA repeats, resulting in compaction of the centromere with repetitive DNA sequences. Recently, Chardon et al. (Chardon et al., 2022) reported that CENP-B compacts the centromeric chromatin. CENP-B is an alpha-satellite binding protein that localizes to human centromeres, but its homolog has not been detected in chickens. CENP-B may cause the hypercompaction of centromeres after CENP-C depletion in human cells. However, because there is no CENP-B homolog in chickens, this may not occur in chicken non-repetitive centromeres. Nevertheless, we would like to emphasize that CENP-C is involved in centromere-specific chromatin organization via oligomerization.

We found that Mini-CENP-C was sufficient for CENP-C function. This further supports our previous observations that two CENP-C functional domains, the Mis12- and CENP-A-binding domains, are dispensable for CENP-C function in chicken DT40 cells (Hara et al., 2018; Watanabe et al., 2019; Watanabe et al., 2022). Given that the Mis12- and CENP-A-binding domains are related to mitotic kinetochore functions, our results lead to an intriguing idea that CENP-C plays no direct essential roles in kinetochore function during mitosis in tissue culture cells. Indeed, acute CENP-C depletion did not rapidly accumulate mitotic cells, but CENP-H depletion did (in this study) (Fukagawa et al., 2001; Nishimura et al., 2019). Furthermore, the centromeric signals from Mini-CENP-C during mitosis were very weak, but they increased in interphase cells, which also supports the idea that CENP-C is not directly involved in essential mitotic kinetochore function. However, CENP-C is required for the maintenance of centromere chromatin integrity during S-phase (Giunta and Funabiki, 2017; Zasadzinska et al., 2018; Nechemia-Arbely et al., 2019). These roles, as well as unidentified functions in interphase, would make CENP-C essential. However, given the evolutionary conservation of the Mis12- and CENP-A-binding domains, it is unlikely that these domains are completely dispensable. CENP-C may play an important role in mitosis that could not be detected under the tissue culture conditions

CENP-C is one of the proteins that were first identified as human centromere components over 35 years ago (Earnshaw and Rothfield, 1985). Since then, many studies on CENP-C have been conducted to determine its essential functions in the centromere/kinetochore. Despite the recent sophisticated structural model of CCAN (Pesenti et al., 2022; Yatskevich et al., 2022), the fundamental role(s) of CENP-C for the centromere/kinetochore in the vertebrate cells remained unclear. In this study, we addressed this long-standing question, and our findings will greatly contribute to understanding how the centromere/kinetochore is formed and maintained in vertebrate cells.

## Limitations of the Study

Whereas we propose a model in which Cupin oligomerization is physically involved in the centromeric chromatin architecture (Figure 7B), we cannot rule out the possibility that CENP-C oligomerization is only required for centromeric localization of CCAN proteins and is not directly associated with the architecture of centromeric chromatin, because CENP-C mutants that lacked the ability to oligomerize failed to localize to centromeres and reduced the centromeric localization of CENP-N. Nevertheless, our results highlight the significance of CENP-C oligomerization for its functions.

## STAR METHODS

### Key Resources Table

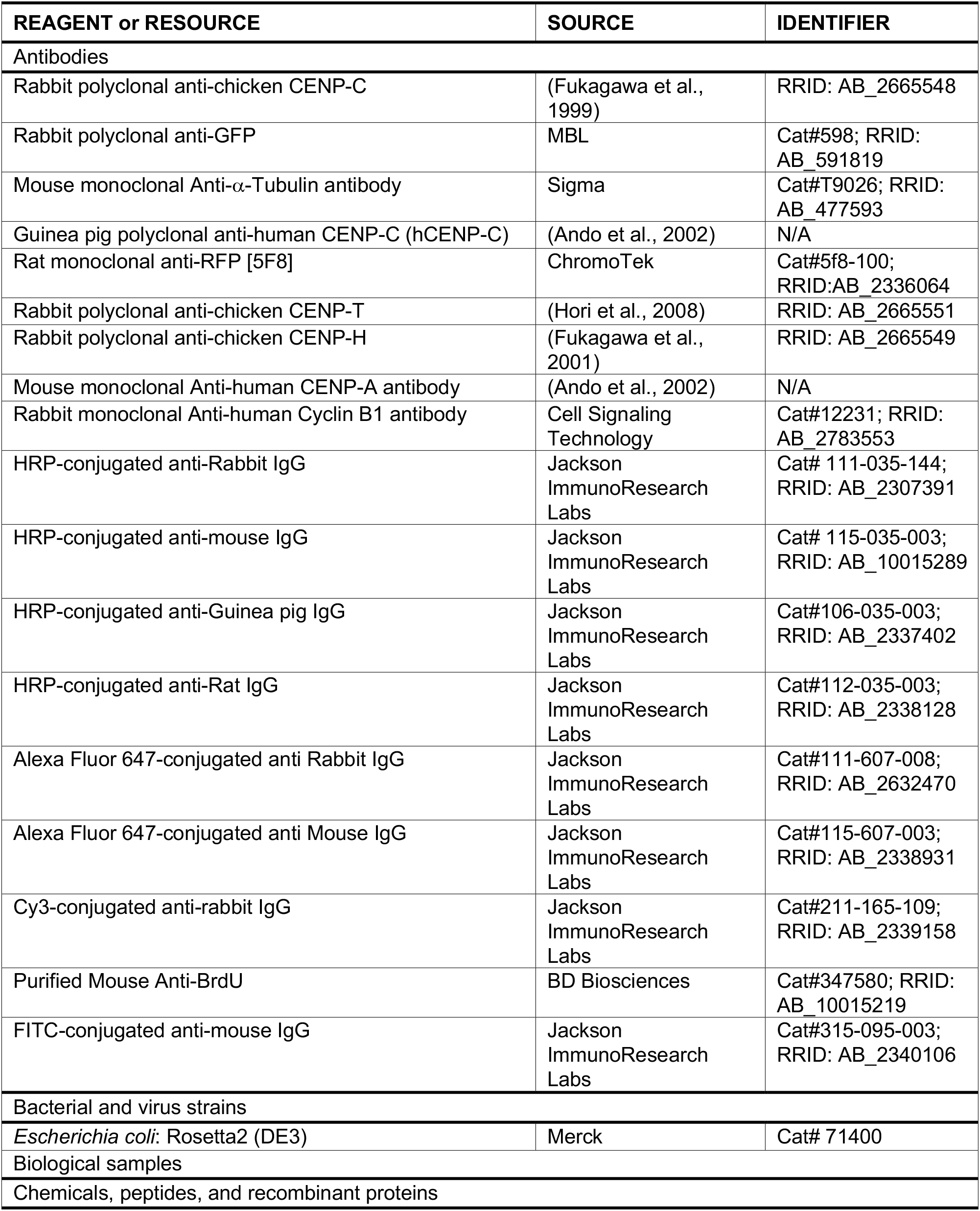
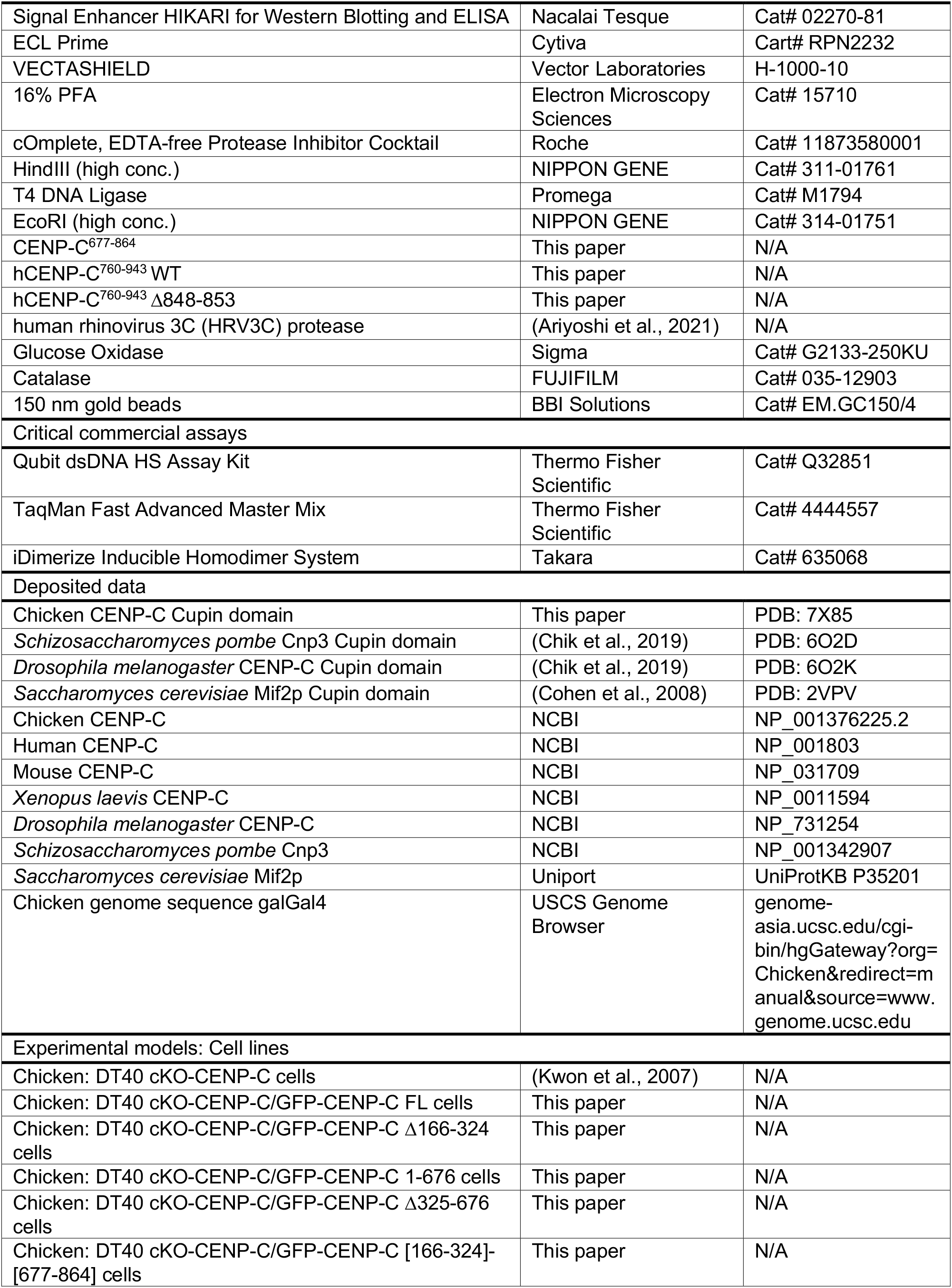
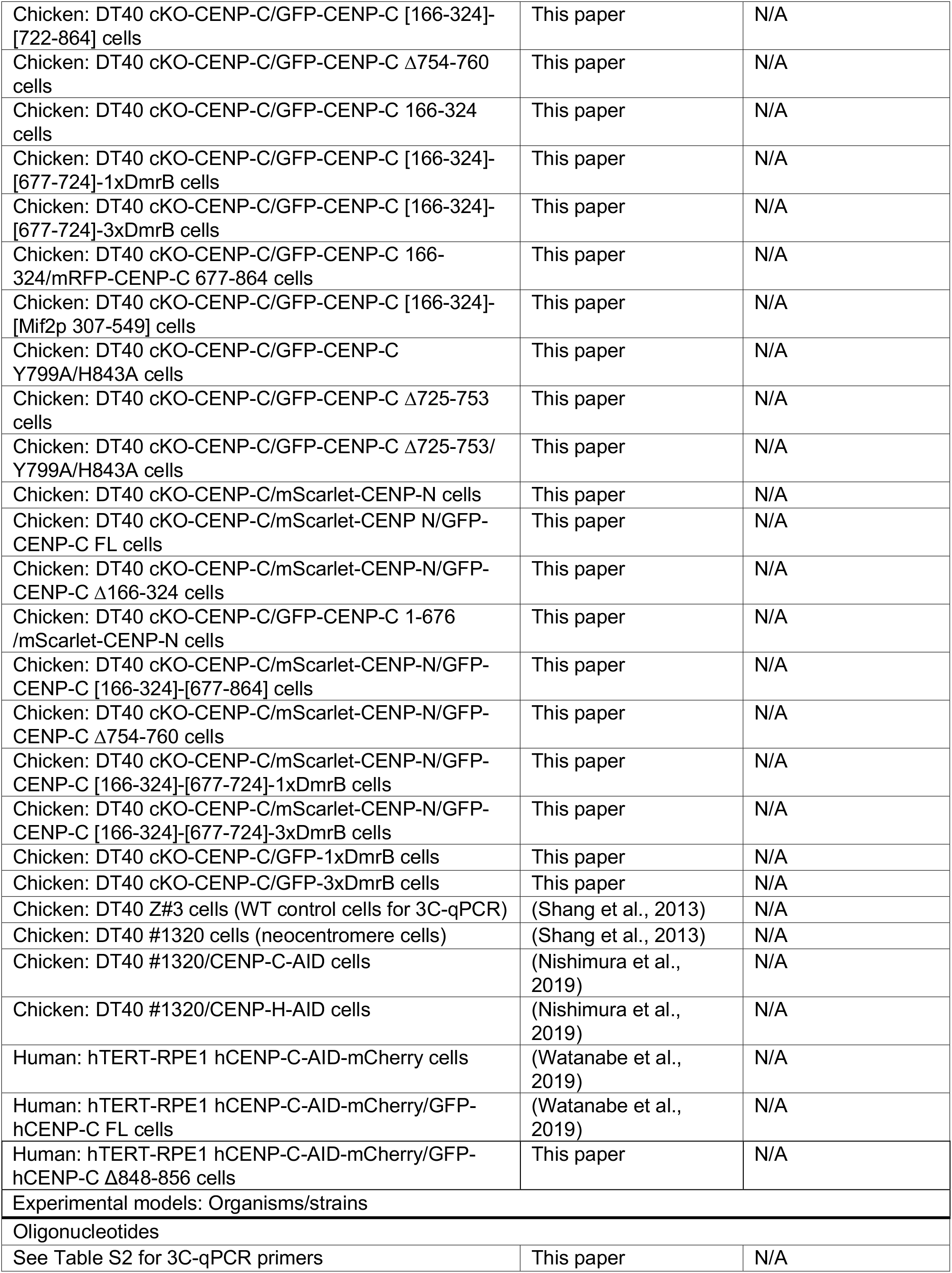
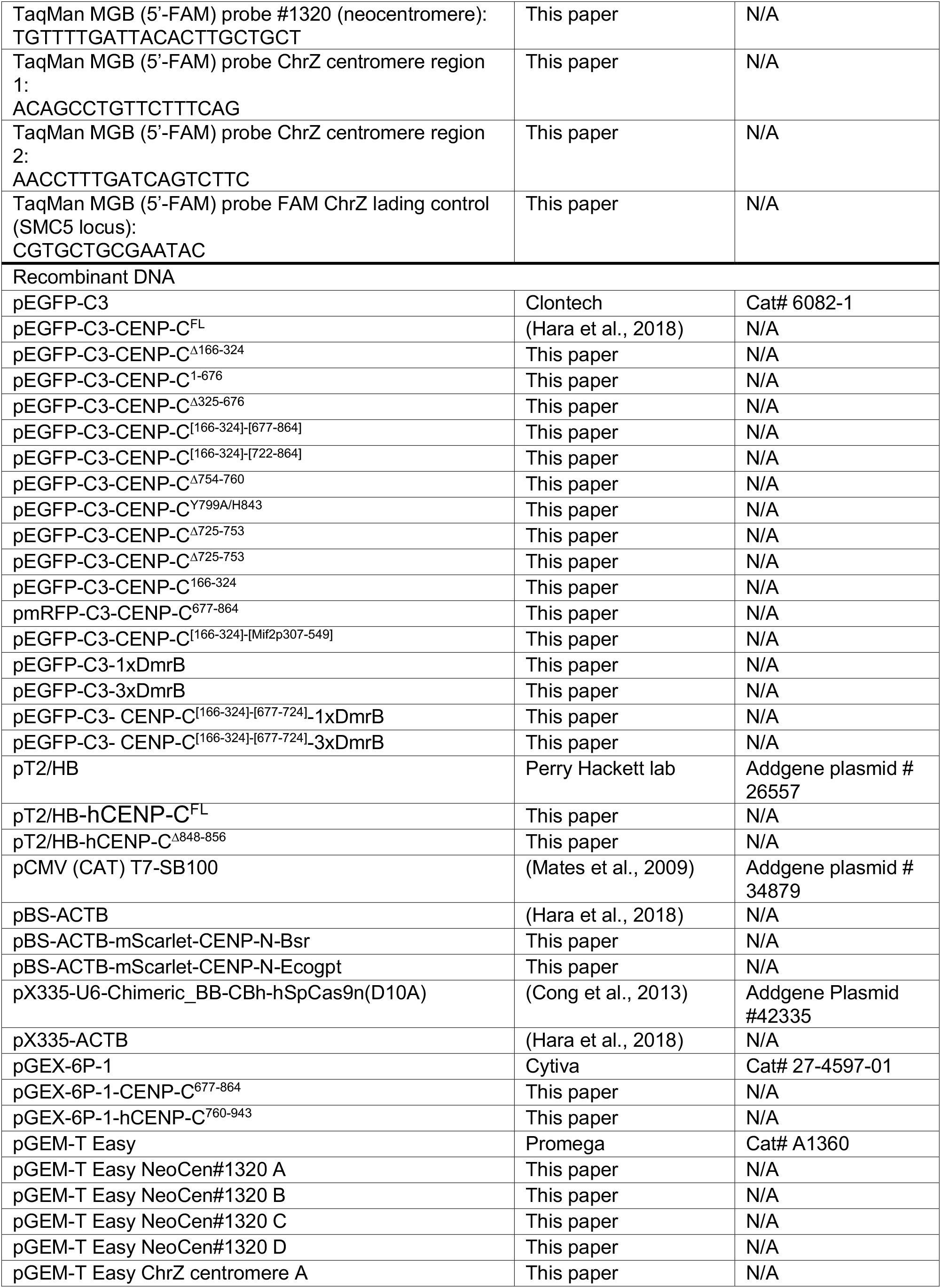
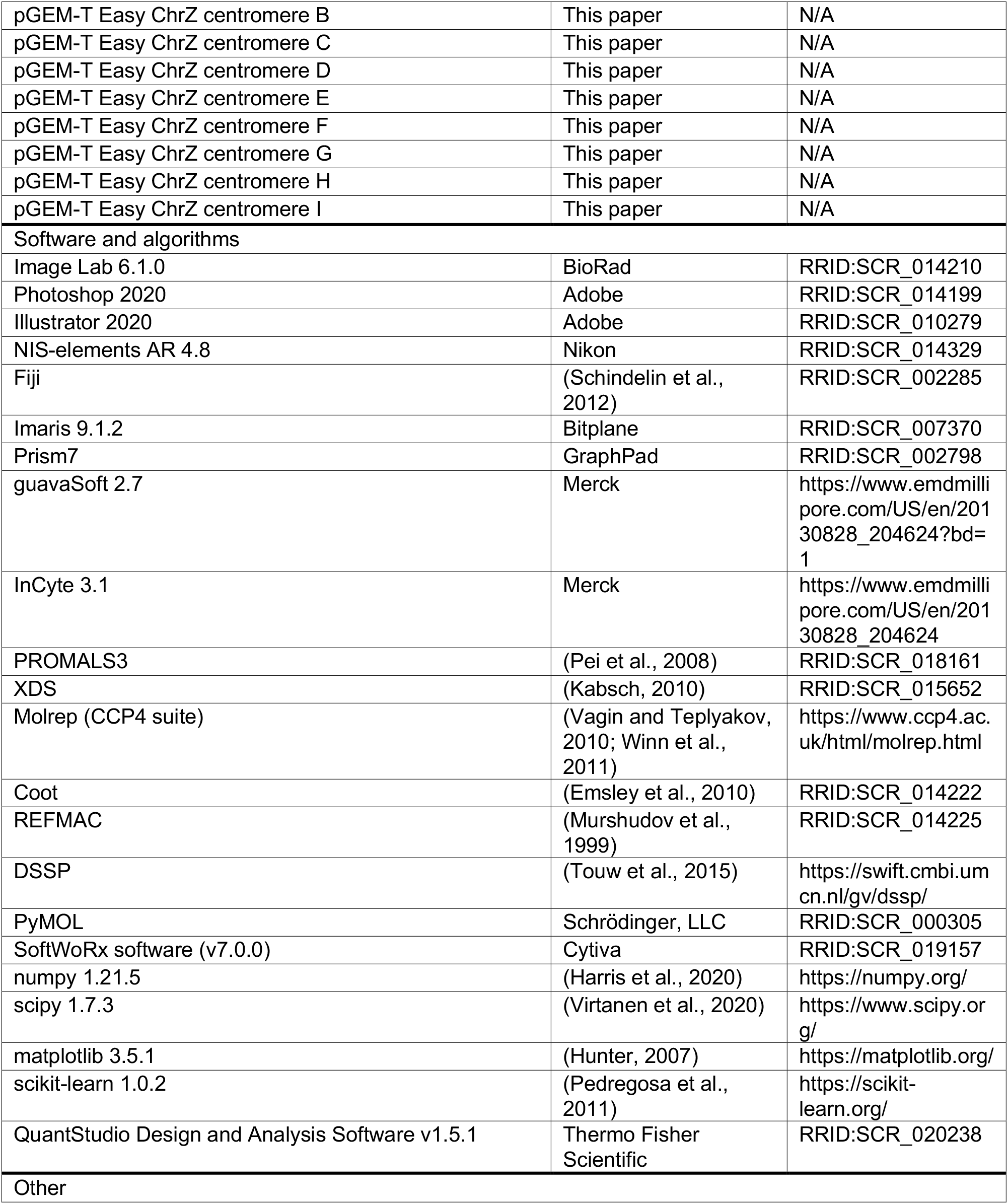

### Resource availability

#### Lead contact

Further information and requests for resources and reagents should be directed to and will be fulfilled by the Lead Contact, Tatsuo Fukagawa.

#### Materials availability

All unique reagents generated in this study are available from the lead contact with a completed Materials Transfer Agreement.

#### Data and code availability

The coordinates and structure factors are available from the PDB with accession code 7X85, 6O2D, 6O2K, 2VPV. Any additional information required to reanalyze the data reported in this paper is available from the lead contact upon request.

### Experimental model and subject details

#### Chicken DT40 cells

A chicken DT40 cell line CL18 was used as the wild-type (WT) cell (Buerstedde et al., 1990). DT40 cells were cultured at 38.5°C in DMEM medium (Nacalai Tesque) supplemented with 10% fetal bovine serum (FBS; Sigma), 1% chicken serum (Thermo Fisher Scientific), and Penicillin-Streptomycin (Thermo Fisher Scientific) (DT40 culture medium).

The chicken CENP-C conditional knockout (cKO-CENP-C) DT40 cell line was described before (Kwon et al., 2007). The cKO-CENP-C cell line expressing GFP-CENP-C full-length (FL: cKO-CENP-C/GFP-CENP-C^FL^) (Hara et al., 2018). To conditionally knockout CENP-C, the cells were cultured in DT40 culture medium containing 2 μg/ml tetracycline (Tet, Sigma).

The cKO-CENP-C expressing GFP-CENP-C Δ166-324, 1-676 or Δ325-676 (cKO-CENP-C/GFP-CENP-C^Δ166-324^, cKO-CENP-C/GFP-CENP-C^1-676^, cKO-CENP-C/GFP-CENP-C^Δ325-676^) (Nagpal et al., 2015) was re-established in this study by transfection of a plasmid encoding GFP-fused CENP-C Δ166-324, 1-676 or Δ325-676 and a neomycin resistance gene into cKO-CENP-C cells using electroporation. Briefly, plasmid constructs were linearized by appropriate restriction enzymes and transfected into DT40 cells using Gene Pulser II electroporator (Bio-Rad). The transfected cells were selected in DT40 culture medium containing 2 mg/ml G418 (Santa Cruz Biotechnology).

The cKO-CENP-C cell line expressing GFP-CENP-C [166-324]-[677-864], [166-324]-[722-864], Δ754-760, 166-324, [166-324]-[677-724]-1xDmrB, [166-324]-[677-724]-3xDmrB, [166-324] fused with Mif2p^307-549^, Y799A/H843, Δ725-753, Δ725-753/Y799A/H843 was generated by transfection of a plasmid encoding GFP-fused CENP-C with indicated mutations as described above.

To generate cKO-CENP-C cell line expressing both GFP-CENP-C^166-324^ and mRFP-fused CENP-C^677-864^, a plasmid encoding mRFP-fused CENP-C C^677-864^ was transfected with a plasmid encoding *Egogpt* gene into the cKO-CENP-C cell line expressing GFP-CENP-C^166-324^ as described above. The transfected cells were selected in DT40 culture medium containing 25 μg/ml Mycophenolic acid (Tokyo Chemical Industry) and 125 μg/ml Xanthine (Sigma). (cKO-CENP-C/GFP-CENP-C 166-324/mRFP-CENP-C 677-864).

To express mScarlet-fused chicken CENP-N under control of endogenous beta-actin (ACTB) promoter, mScarlet-CENP-N was integrated into endogenous ACTB locus using CRISPR/Cas9 genome editing as previously described (Hara et al., 2018). cKO-CENP-C cell lines expressing indicated GFP-CENP-C protein were transfected with a plasmid encoding SpCas9 nickase (D10A) (pX335-U6-Chimeric_BB-CBh-hSpCas9n(D10A): a gift from Feng Zhang, Addgene plasmid # 42335) (Cong et al., 2013) and gRNA sequence for ACTB (Hara et al., 2018) together with donor plasmids, which contained mScarlet-CENP-N cDNA and a drug resistance gene expression cassette (BSR or Ecogpt) between 5’ and 3’ homology arm regions (approximately 1 kb each) franking ACTB start codon using Neon Transfection System (Thermo Fisher Scientific). The targeted cells were selected in DT40 culture medium containing 0.5 μg/ml Blasticidin S (FUJIFILUM), 25 μg/ml Mycophenolic acid and 125 μg/ml Xanthine. (cKO-CENP-C/mScarlet-CENP-N/GFP-CENP-C FL, Δ754-760, [166-324]-[677-724]-1xDmrB, [166-324]-[677-724]-3xDmrB, Δ166-324, 1-676, [166-324]-[677-864]).

To induce dimerization or oligomerization of GFP-CENP-C [166-324]-[677-724]-1xDmrB or [166-324]-[677-724]-3xDmrB, respectively, cKO-CENP-C cells expressing these proteins were treated with 100 nM B/B Homodimerizer (Takara).

#### Human RPE-1 cells

RPE-1 cells were cultured at 37°C in DMEM medium (Nacalai Tesque) supplemented with 10% FBS (Sigma) and Penicillin-Streptomycin (Thermo Fisher Scientific) (RPE-1 culture medium).

The RPE-1 cell line in which Human CENP-C (hCENP-C) protein was conditional depleted by auxin-inducible degron was previously described system (RPE-1 hCENP-C-AID cell) (Watanabe et al., 2019). The hCENP-C-AID cell line expressing GFP-fused hCENP-C FL was also described in Watanabe et al. (Watanabe et al., 2019). To deplete the AID-tagged hCENP-C (hCENP-C-AID-mCherry), the cells were cultured in RPE-1 culture medium containing 500 μM IAA (indole-3-acetic acid, Sigma).

To establish RPE-1 hCENP-C-AID cell line expressing GFP-fused hCENP-C^Δ848-856^, the GFP-fused hCENP-C^Δ848-856^ cDNA was integrated in the genome using Sleeping Beauty transposon system (Mates et al., 2009) (RPE-1 hCENP-C-AID /GFP-hCENP-C^Δ848-856^). Plasmid constructs were transfected into RPE-1 hCENP-C-AID cells with Neon Transfection System. Since the transgene cassette has a neomycin resistance gene, the cell lines were selected in RPE-1 culture medium containing 500 μg/ml G418.

### Method Details

#### Cell counting

To count DT40 cell numbers, the culture was mixed with the same volume of 0.4% (w/v) Trypan Blue Solution (FUJIFILM) and the cell numbers were counted by Countess II (Thermo Fisher Scientific).

To count RPE-1 cell numbers, the cells were trypsinized with 2.5 g/l-Trypsin-1 mmol/l EDTA solution (Nacalai tesque) and suspended into RPE-1 culture medium. The cell suspension was mixed with the same volume of 0.4% (w/v) Trypan Blue Solution (FUJIFILM) and the cell numbers were counted by Countess II (Thermo Fisher Scientific).

#### Plasmid constructions

Chicken CENP-C full-length (FL) cDNA sequence (NP_001376225.2) (Fukagawa and Brown, 1997) was cloned into pEGFP-C3 (Clontech) (pEGFP-C3-CENP-C^FL^) (Hara et al., 2018). To prepare GFP-fused CENP-C mutants, deletion or mutation was introduced into pEGFP-C3-CENP-C^FL^ (pEGFP-C3-CENP-C Δ166-324, 1-676, Δ325-676, [166-324]-[677-864], [166-324]-[722-864], Δ754-760, 166-324, Y799A/H843, Δ725-753, Δ725-753/Y799A/H843). CENP-C^166-324^ was cloned into pEGFP-C3 (pEGFP-C3-CENP-C^166-324^). CENP-C^677-864^ was cloned into pEGFP-C3 whose EGFP was replaced with mRFP (pmRFP-CENP-C^677-864^). *MIF2^307-549^* cDNA sequence was amplified by PCR and cloned into pEGFP-C3-CENP-C using InFusion HD (Takara) (pEGFP-C3-CENP-C^[166-324]-[Mif2p307-549]^). Single or three tandem-repeat of DmrB, an inducible homodimerization domain (Takara), was amplified by PCR and cloned into pEGFP-C3 plasmid (Clontech) (pEGFP-C3-1xDmrB, pEGFP-C3-3xDmrB). CENP-C^[166-324]-[677-724]^ was amplified by PCR and cloned into pEGFP-C3-1xDmrB and pEGFP-C3-3xDmrB (pEGFP-C3-CENP-C^[166-324]-[677-724]^-1xDmrB, pEGFP-C3-CENP-C^[166-324]-[677-724]^-3xDmrB).

Chicken CENP-N cDNA-fused with mScarlet and a drug resistance gene expression cassette (BSR or Ecogpt) was cloned in a plasmid which contains ACTB homology arms for knock-in (Hara et al., 2018) (pBS-ACTB, pBS-ACTB-mScarlet-CENP-N-Bsr, pBS-ACTB-mScarlet-CENP-N-Ecogpt).

hCENP-C FL cDNA sequence was cloned into pEGFP-C3 (pEGFP-C3-hCENP-C^FL^) (Watanabe et al., 2019). aa 848-856 was deleted from pEGFP-C3-hCENP-C^FL^ by PCR (pEGFP-C3-hCENP-C^Δ848-856^). The GFP-CENP-C expression cassette together with neomycin resistance gene cassette in the pEGFP-C3 plasmid was amplified by PCR and cloned into pT2/HB (a gift from Perry Hackett, Addgene plasmid # 26557), a plasmid used for Sleeping Beauty transposon system (Mates et al., 2009), using InFusion HD (Takara) (pT2/HB-hCENP-C^FL^, pT2/HB-hCENP-C^Δ848-856^). The plasmids were used with pCMV (CAT) T7-SB100 (a gift from Zsuzsanna Izsvak, Addgene plasmid # 34879) (Mates et al., 2009).

Chicken CENP-C^677-864^ cDNA sequence was cloned into inserted pGEX-6P-1 (Cytiva) for bacterial protein expression (pGEX-6P-1-CENP-C^677-864^). hCENP-C^760-943^ cDNA sequence was cloned into pGEX-6P-1 (Cytiva) for bacterial protein expression (pGEX-6P-1-hCENP-C^760-943^). Amino acid residues 848-856 were deleted from pGEX-hCENP-C^760-943^ by PCR (pGEX-6P-1-hCENP-C^760-943Δ848-856)^.

For the 3C-qPCR standards, the ligated chimeric DNA fragments of target regions in 3C library were amplified and cloned into pGEM-T Easy (Promega)

#### Immunoblotting

DT40 cells were harvested, washed with PBS and suspended in 1xLSB (Laemmli sample buffer) (final 1×10^4^ cells/μl), followed by sonication and heating for 5 min at 96°C. In Figure S6C, insoluble fractions of DT40 cells were examined; DT40 cells were harvested, washed with PBS twice, followed by another wash with TMS (10 mM Tris-HCl pH 7.5, 5 mM MgCl_2_, 0.25 M sucrose). The cells were suspended in TMS supplemented with 0.5% Triton X-100 (TMS-Triton) and incubated on ice for 5 min. After centrifugation, proteins of the precipitates were extracted by adding 1xLSB, sonication and heating for 5 min at 96°C. Proteins were separated on SuperSep Ace, 5-20% (FUJIFILM) and transferred to Immobilon-P (Merck) using HorizeBLOT (ATTO).

RPE-1 cells were trypsinized and harvested, washed with PBS and suspended in 1xLSB (Laemmli sample buffer) (final 1×10^4^ cells/μl) followed by sonication and heating for 5 min at 96°C. Proteins were separated on SuperSep Ace, 5-20% (FUJIFILM) and transferred to Immobilon-P (Merck) using HorizeBLOT (ATTO).

Primary antibodies used in this study were rabbit anti-chicken CENP-C (Fukagawa et al., 1999), rabbit anti-GFP (MBL), mouse anti-alpha tubulin (Sigma), guinea pig anti-hCENP-C (Ando et al., 2002), rat anti-RFP (Chromotek), rabbit anti-chicken CENP-H (Fukagawa et al., 2001), and rabbit anti-chicken CENP-T (Hori et al., 2008). Secondary antibodies were HRP-conjugated anti-Rabbit IgG (Jackson Immuno Research), HRP-conjugated anti-Mouse IgG (Jackson ImmunoResearch), HRP-conjugated anti-Rat IgG (Jackson ImmunoResearch), and HRP-conjugated anti-guinea pig (Sigma). To increase sensitivity and specificity, Signal Enhancer Hikari (Nacalai Tesque) was used to dilute all antibodies except for anti-alpha tubulin (5% skim milk in TBS 0.05% tween) and HRP-conjugated anti-mouse IgG (TBS 0.05% tween). The antibodies were incubated with the blotted membranes for 1 h at room-temperature or for overnight at 4°C. Proteins reacting with antibodies were detected with ECL Prime (Cytiva) and visualized with ChemiDoc Touch (BioRad). Acquired images were processed using Image Lab 6.1.0 (BioRad) and Photoshop 2020 (Adobe).

#### Immunofluorescence

For cKO-CENP-C DT40 cells expressing GFP-CENP-C, the cells were cytospun onto glass slides (Matsunami). The cells were fixed with 4% paraformaldehyde (PFA) in PHEM buffer (60 mM PIPES, 25 mM HEPES, 10 mM EGTA, 2 mM MgCl_2_, pH 7.0 (NaOH)) for 10 min, permeabilized with 0.5% Triton X-100 in PHEM for 10 min, blocked with 0.5% BSA in PBS for 5 min and incubated with Cy3-labeled anti-chicken CENP-T (Hori et al., 2008) diluted with 0.5% BSA in PBS for 1 h at 37°C and then washed with 0.5% BSA in PBS 3 times. The cells were post-fixed with 4%PFA in PHEM for 10 min, washed with PBS and stained DNA with 100 ng/ml DAPI in PBS for 15 min. The stained samples were washed with PBS and mounted with VECTASHIELD Mounting Medium (Vector Laboratories).

For cKO-CENP-C DT40 cells expressing GFP-CENP-C and mScarlet-CENP-N, the cells were cytospun onto precision cover glasses (#1.5H Thickness, Tholabs). The cells were fixed and permeabilized as above. After blocking with 0.5% BSA in PBS for 5 min, the cells were incubated with anti-chicken CENP-T (Hori et al., 2008) diluted with 0.5% BSA in PBS for overnight at 4°C. After 3 times wash with 0.5% BSA in PBS, the cells were incubated with Alexa Fluor 647-conjugated anti-rabbit IgG (Jackson ImmunoResearch) diluted with 0.5% BSA in PBS for 1 h at 37°C and then washed with 0.5% BSA in PBS 3 times. The cells were post-fixed with 4%PFA in PHEM for 10 min, washed with PBS and stained DNA with 100 ng/ml DAPI in PBS for 15 min. The stained samples were washed with PBS and mounted with VECTASHIELD Mounting Medium (Vector Laboratories).

RPE-1 cells were cultured on precision cover glasses (#1.5H Thickness, Tholabs). The cells were fixed with 4% paraformaldehyde (PFA) in PHEM buffer for 10 min, permeabilized with 0.5% Triton X-100 in PHEM for 10 min, rinsed with 0.1% Triton X-100 in PBS, blocked with blocking solution (3% BSA, 0.1% Triton X-100, 0.1% NaN_3_ in TBS) for 10 min, and then incubated with primary antibodies: mouse anti-human CENP-A (Ando et al., 2002) and rabbit anti-human Cyclin B1 (Cell Signaling Technology) diluted with blocking solution for overnight at 4°C. After 3 times wash with 0.1% Triton X-100 in PBS, the cells were incubated with Alexa Fluor 647-conjugated anti-mouse IgG (Jackson ImmunoResearch) and Cy3-conjugated anti-rabbit IgG (Jackson ImmunoResearch) diluted with blocking solution for 1 h at RT and then washed with 0.1% Triton X-100 in PBS 3 times. The cells were post-fixed with 4%PFA in PHEM for 10 min, washed with PBS and stained DNA with 20 μM NucleoSeeing (Funakoshi) in PBS for 15 min. The samples were stored in PBS at 4°C for STORM and MINFLUX nanoscopy analysis.

#### Spinning-disk confocal microscopy

Immunofluorescence images were acquired at 0.2 μm intervals in the z-axis using a Zyla 4.2 sCMOS camera (Andor) mounted on a Nikon Ti inverted microscope with an objective lens (Nikon; Plan Apo lambda 100x/1.45 NA) with a spinning-disk confocal unit (CSU-W1, Yokogawa) controlled with NIS-elements AR 4.8 (Nikon) at RT. The images in figures are the maximum intensity projection of the Z-stack generated with Fiji (Schindelin et al., 2012). Acquired images were processed using Fiji (Schindelin et al., 2012) and Photoshop 2020 (Adobe).

#### Quantification of mScarlet-CENP-N on mitotic centromeres

The fluorescence signal intensities of mScarlet-CENP-N co-localized with CENP-T on mitotic centromere with in DT40 cells were quantified using Imaris (Bitplane). The fluorescence signals on about 50-100 centromeres in each of 10 mitotic cells were quantified and subtracted with mean of background signals in non-centromeric regions in each cell and the mean value of the mScarlet-CENP-N signals in each cell was calculated. Data are shown as Mean ± standard deviation using GraphPad Prims7 (GraphPad software). The unpaired t-test (two-tailed) was used. A p value less than 0.05 is considered statistically significant.

#### Flow cytometry analysis

Ten ml of DT40 cell culture (5×10^6^ cells/ml) was labeled with 20 μM BrdU (5-bromo-2’-deoxyuridine) for 20 min and harvested. The cells were washed with 10 ml ice-cold PBS, gently suspended with 13 ml 70% ethanol and stored at – 20°C. Following wash with 1 ml 1%BSA in PBS, the cells were incubated in 1 ml 4 N HCl with 0.5% Triton X-100 for 30 min at RT and washed with 1 ml 1% BSA in PBS 3 times. The cells were incubated with 30 μl anti-BrdU (BD) for 1 h at RT, washed with 1 ml 1% BSA in PBS twice, incubated with 30 μl FITC-labeled anti-mouse IgG (Jackson ImmunoResearch, 1/20 in 1%BSA in PBS) for 30 min at RT, washed with 1 ml 1% BSA in PBS and stained DNA with 1 ml propidium iodide (10 μg/ml) in 1%BSA in PBS for overnight at 4°C. The stained cells were applied to Guava easyCyte (Merck). Obtained data were analyzed with InCyte software (Merck).

#### Protein expression and purification

*E. coli* cells (Rosetta2(DE3), Novagen) were transformed with pGEX-6P-1-CENP-C^677-864^ to express chicken CENP-C^677-864^ as a glutathione-S-transferase (GST)-fused recombinant protein. Cells were grown at 37°C until OD600 reached 0.6; protein expression was induced by addition of IPTG to a final concentration of 0.2 mM, and culture was continued for overnight at 17°C. Cells were harvested by centrifugation. The cell pellet was resuspended in buffer A (20 mM HEPES-NaOH pH 7.5, 500 mM NaCl, 5% Glycerol and 2 mM tris(2-carboxyethyl)phosphine (TCEP) and lysed by sonication on ice. The lysate was clarified by centrifugation and applied to a glutathione sepharose 4FF column (Cytiva) pre-equilibrated with buffer A. After extensive column washing with buffer A containing 1 M NaCl, the tag-free protein was eluted from the column by the GST-tag cleavage with human rhinovirus 3C (HRV3C) protease. The CENP-C^677-864^ fragment was applied to a HiTrap SP HP cation exchange column (Cytiva), eluted with a linear NaCl gradient from 100 mM to 750 mM, and further purified by size-exclusion chromatography using HiLoad Superdex 16/60 200 pg column (Cytiva) in a buffer containing 20 mM HEPES-NaOH pH 7.5, 500 mM NaCl and 2 mM TCEP. Peak fractions containing the Cupin domain were combined, concentrated (typically to 8-10 mg/ml), and stored at - 80°C until further use for crystallization and biochemical assay.

GST-fused hCENP-C^760-943^ or hCENP-C^760-943Δ848-856^ was expressed in *E. coli* cells (Rosetta2(DE3)), lysed and purified by glutathione affinity column chromatography in the same manner to GST-fused chicken CENP-C^677-864^. After the GST tag cleavage with HRV3C protease, the hCENP-C^760-943^ or hCENP-C^760-943Δ848-856^ fragment was purified using a HiTrap SP HP cation exchange column (Cytiva) with a linear NaCl gradient from 75 mM to 850 mM for elution. The peak fractions containing the hCENP-C fragment was further purified by size-exclusion chromatography using HiLoad Superdex 16/60 200 pg column (Cytiva) in a buffer containing 20 mM HEPES-NaOH pH 7.5, 300 mM NaCl, 5% Glycerol and 2 mM dithiothreitol (DTT). Peak fractions containing the hCENP-C^760-943^ or hCENP-C^760-943Δ848-856^ were combined and concentrated (2 mg/ml of hCENP-C^760-943^, 4 mg/ml of hCENP-C^760-943Δ848-856^).

#### Crystallization and structural determination

Crystallization of the chicken CENP-C^677-864^ was performed using the sitting-drop vapor diffusion method at 20°C. Octahedral crystals were grown from drops consisting of 200 nl of protein solution (10.4 mg/ml) and 100 nl of reservoir solution containing 0.2 M lithium citrate, 0.1 M MES-NaOH pH 6.5 and 20% PEG4000. For X-ray diffraction measurements, the crystals were cryoprotected in reservoir solution supplemented with 20% Glycerol. X-ray diffraction datasets were collected at 100 K on the synchrotron beamline BL-26B1 at SPring8 (Himeji, Japan). Diffraction data were processed using XDS (Kabsch, 2010). Data collection statistics are summarized in Table S1. The structure of the chicken CENP-C Cupin domain was determined by the molecular replacement method using Molrep (Vagin and Teplyakov, 2010) in the CCP4 suite (Winn et al., 2011). The coordinates of the *Saccharomyces cerevisiae* Mif2p Cupin domain (PDB ID: 2VPV) were used as a search model. The initial model was built using Coot (Emsley et al., 2010). Structural refinement was conducted using REFMAC (Murshudov et al., 1999). The final model containing residues 722-849 was built by iterative reciprocal space refinement and manual rebuilding. The statistics of structural refinement and the quality of the final model are summarized in Table S1. Secondary structure assignment was performed using DSSP (Touw et al., 2015). All figures depicting the crystal structure were produced using the PyMOL molecular graphics system (Schrödinger, LLC).

#### Blue native-polyacrylamide gel electrophoresis (PAGE)

Eight to ten micrograms of purified chicken CENP-C^677-864^, hCENP-C^760-943^ or hCENP-C^760-943Δ848-856^ were dissolved in 5 μl Native PAGE sample buffer (50 mM Tris-HCl pH 7.5, 10% Glycerol, 50 mM NaCl, 0.001% Ponceau S, 0.5% Coomassie G-250) and applied to a 10%–20% gradient gel (NativePAGE Bis-Tris gel; Thermo Fisher Scientific) and migrated at 150 V for 180 min at 4°C. Native PAGE running buffer (50 mM Bis-Tris, 50 mM Tricine, pH 6.8) was used as a positive electrode and Native PAGE running buffer supplemented with 0.02% Coomassie G-250 was used as a negative electrode. The bands were visualized by destaining the gel with water.

#### In vitro condensation assay

The CENP-C^677-864^ fragment (final 54 μM) was diluted into condensation buffer (25 mM Tris-HCl pH 7.5, 5 mM MgCl_2_, 1 mM DTT, 5% Glycerol) with or without 5% PEG8000. Ten percent of 1,6-hexamediol was added into condensation buffer with PEG8000 to prevent the coacervate formation. Immediately after dilution, five microliters of the mixtures were dropped onto a cover glass (Matsunami). Differential interference contrast images of the CENP-C condensates were observed using the DeltaVision Elite system (GE Healthcare Inc., Chicago, USA) equipped with pco.edge 4.2 sCMOS camera (PCO, Kelheim, Germany) and 60× PlanApo N OSC oil-immersion objective lens (numerical aperture [NA] = 1.4, Olympus, Tokyo, Japan). The microscope was operated with the built-in SoftWoRx software (v7.0.0). The brightness of the images was changed using Fiji (Schindelin et al., 2012) for better visualization without changing the gamma settings. Acquired images were processed using Fiji (Schindelin et al., 2012) and Photoshop CC (Adobe).

#### 2D-STORM

The immunostained RPE-1 cells on the cover glass were mounted with freshly prepared oxygen scavenger imaging buffer (50 mM Tris pH8.0, 10 mM NaCl, 10% Glucose, 0.7 µg/µL Glucose Oxidase (Sigma, #G2133-250KU), 0.04 µg/µL Catalase (FUJIFILM, #035-12903) and 70 mM 2-mercaptoethanol) based on the glucose oxidase enzymatic system (GLOX) for STORM using a frame-sealed incubation chamber (Bio-Rad). 2D-STORM acquisitions were performed using a Nikon N-STORM with Eclipse Ti-E inverted microscope and laser TIRF illuminator (Nikon Instruments Inc.). Alexa 647 fluorophores were stochastically excited using the 640 nm laser beam with an additional 405 nm weak pulse. Images were acquired with an Andor iXon 897 EMCCD camera (Andor Technologies) and a CFI Apo TIRF 100× objective lens (N.A. 1.49). For each cell, a stack of 10,000 frames was acquired at an exposure time of 16 ms and information including the XY coordinates of the detected blinking spots was extracted using Nikon NIS-Element imaging software (Nikon Instruments Inc.).

#### MINFLUX nanoscopy

For the stabilization process during MINFLUX measurements, the immunostained RPE-1 cells as above were incubated for 5-10 min with an undiluted dispersion of 150 nm gold beads (EM.GC150/4, BBI Solutions). After washing with PBS, the samples were mounted with GLOX+MEA imaging buffer (50 mM TRIS/HCl, 10 mM NaCl, 10% (w/v) Glucose, 64 µg/ml catalase, 0.4 mg/ml glucose oxidase, 25-30 mM MEA, pH 8.0) (Balzarotti et al., 2017). The MINFLUX measurements and the corresponding confocal images were acquired with an abberior MINFLUX (Abberior Instruments GmbH) equipped with a 642 nm (CW) excitation laser for confocal and MINFLUX imaging, a 488 (CW) excitation laser for confocal imaging and a 405-nm (CW) laser for activation. The 642 nm beam was shaped using a spatial light modulator. The EOD-based MINFLUX scanning was performed as described before (Schmidt et al., 2021). A 60x magnification 1.42 NA oil objective lens (UPLXAPO60XO, Olympus) was used for all measurements. The emission upon 640 nm excitation was detected on two avalanche photodiodes with fluorescence filters from 650 to 780 nm. The fluorescence emitted at 488 nm excitation was counted on an avalanche photodiode with a fluorescence filter from 500 to 620 nm. To stabilize the sample during MINFLUX measurements, a reflection-based stabilization system with a 980 nm laser was applied (Schmidt et al., 2021).

#### Rendering 2D-STORM and MINFLUX images

Two-dimensional single-molecule localization microscopy data mainly consists of the localization coordinates (*x_n_*, *y_n_*), the intensity *I_n_*, and the lateral localization accuracy *σ*_*n*_ for the *n*th blinking dot: 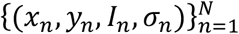. Then, we calculated the total intensity by

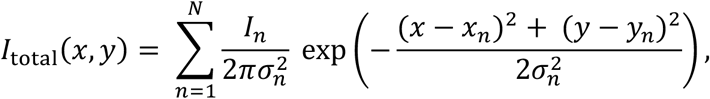

and mapped it on a pixel region with the Gaussian filter for smoothing. Here, we developed the rendering algorithm as Python code. In MINFLUX, since the localization precision needs to be determined empirically via the spread of the localizations on each trace (Schmidt et al., 2021), we set *σ_n_* = 2 nm.

#### Pair correlation analysis

First, to crop appropriate clusters of dots of fluorophore emitters labeling CENP-A molecules, we applied the density-based clustering algorithm DBSCAN (Density-Based Spatial Clustering of Applications with Noise). We set the main two parameters *eps* = 100 nm, which is the maximum distance between two dots for one to be considered to be connected to the other, and *mnsPts* = 5, the minimum point number in a neighborhood for a dot to be considered as a core dot.

Then we calculated the normalized pair correlation function for each cropped localization coordinate data 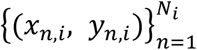 of the *i*th cluster in the cropped rectangular region. In principle, the pair correlation function for randomly distributed dots reveals the unit value. However, the pair correlation function for dots in a cropped region inevitably includes the boundary effect. In order to eliminate the boundary effect, we numerically prepared uniformly distributed *N_i_* dots in the same rectangle region for the *i*th cluster. Based on the pair-wise Euclidean distance between dot *n* and *m*, *r_nm_*, we calculated the histogram *h*_cluster_(*r*) and *h*_random_(*r*) of the distance for both clustered and random dots with the 2-nm binning width. For smoothing *h*_random_(*r*), we prepared 100 sets of random dots and averaged these histograms. Finally, the normalized pair correlation function was calculated by: *g*(*r*) = *h*_cluster_(*r*)/ *h*_random_(*r*) (Figure S5A). From this definition, when the normalized pair correlation function curve monotonically decays from *r* = 0 with *g*(*r*) > 1 and crosses *g*(*r*) = 1, we can state that the dots are clustering within the distance for *g*(*r*) > 1. Therefore, the distance satisfying *g*(*r*) = 1 can be a characteristic length of the clustering dots.

#### 3C-qPCR

The ligation frequencies within the centromeric chromatin were measured by chromosome conformation capture (3C)-qPCR (Hagege et al., 2007) with several modifications to optimize for the centromeric chromatin in DT40 cells.

##### Crosslinking and lysis

DT40 cells (1×10^7^ cells) were suspended in 10 ml fresh DT40 culture medium and crosslinked by adding final 1% PFA (16%, Electron Microscopy Sciences). After 10 min at RT, the cells were moved onto ice and added final 0.2 mM Glycine to quench the crosslinking reaction. The reaction mixture was centrifuged for 5 min at 220 x *g* at 4°C and the supernatant was aspirated. The cell pellets were washed with 10 ml ice-cold PBS twice. After centrifugation for 5 min at 220 x *g* at 4°C, the supernatant was aspirated, and the cell pellets were resuspended in 5 ml lysis buffer (10 mM Tris-HCl pH 7.5, 10 mM NaCl, 0.2% NP-40, 1x complete EDTA-free proteinase inhibitor (Roche)) and incubated for 15 min on ice. The mixture was centrifuged for 5 min at 400 x *g* at 4°C, and the supernatant was aspirated. The cell pellets were resuspended with 5 ml lysis buffer, centrifuged for 5 min at 400 x *g* at 4°C, and the supernatant was aspirated.

##### Restriction enzyme digestion and ligation

The precipitates were resuspended and transferred into a new 1.5 ml tube with 0.5 ml 1.2 x restriction enzyme B-buffer (Nippon Gene). Seven point five microliters of 20% (w/v) SDS (final 0.3%) were added into the suspension. The reaction mixture was incubated for 1 h at 37°C with shaking at 900 rpm. Following 50 μl 20% (v/v) Triton X-100 addition (final 2%), the mixture was incubated for 1 h at 37°C with shaking at 900 rpm. Four hundred and eighty units of HindIII (4 μl of 120 U/μl, Nipone Gene) was added into the reaction and incubated for overnight at 37°C with shaking at 900 rpm. After addition of 40 μl 20% (w/v) SDS (final 1.6%), the mixture was incubated for 20 min at 65°C. The reaction mixture was transferred to 50 ml tube, diluted with 6.125 ml 1.15 x ligation buffer (1 x ligation buffer: 60 mM Tris-HCl pH 7.5, 5 mM DTT, 5 mM MgCl_2_, 1 mM ATP) and supplemented with 375 μl 20% Triton X-100 (final 1%). The mixture was incubated for 1 h at 37°C with shaking at 180 rpm. Five microliters of 20 U/μl T4 ligase (Promega) were added into the mixture and incubated for 4 h at 16°C, followed by further incubation for 30 min at RT. The reaction mixture was digested by adding 15 μl 20 mg/ml Proteinase K and incubation for overnight at 65°C. Digestion efficiency of samples was assessed following Hagège et al (Hagege et al., 2007).

##### DNA purification

Thirty microliters of 10 mg/ml RNase A were added to the reaction mixture and incubate for 30 min at 37°C. The digested sample was mixed with 7 ml Phenol/Chloroform/Isoamyl alcohol (25:24:1, Nippon Gene) and centrifuged for 15 min at RT at 2,200 x *g*. The aqueous phase was transferred to a new tube, mixed with 7 ml Chloroform and centrifuged for 15 min at RT at 2,200 x *g*. The aqueous phase was transferred to a new tube, mixed with 700 μl sodium acetate and 17. 5 ml ethanol and placed at −80°C for overnight. The mixture was centrifuged for 15 min at 4°C at 2,200 x *g*. After removal of supernatant, ten milliliters of 70% ethanol were added into the tube and centrifuged for 15 min at 4°C at 2,200 x *g*. The supernatant was discarded. The pellets were briefly dried at RT, then dissolved in 150 μl 10 mM Tris-HCl pH 7.5 and transferred into a new 1.5 ml tube.

##### Secondary restriction enzyme digestion

The purified DNA was mixed with 25 μl H_2_O, 20 μl 10x restriction enzyme H-buffer (Nippon Gene) and 5 μl 20U/μl EcoRI (final 100 U, Nippon Gene), and incubated for 2 h at 37°C. The digested sample was mixed with 200 μl Phenol/Chloroform/Isoamyl alcohol (25:24:1, Nippon Gene) and centrifuged for 5 min at RT at 16,000 x *g*. The aqueous phase was transferred to a new tube, mixed with 200 μl Chloroform and centrifuged for 5 min at RT at 16,000 x *g*. The aqueous phase was transferred to a new tube, mixed with 20 μl sodium acetate and 0.5 ml ethanol and placed at −80°C for overnight. The mixture was centrifuged for 5 min at 4°C at 16,000 x *g*. After removal of supernatant, half a milliliter of 70% ethanol was added into the tube and centrifuged for 5 min at 4°C at 16,000 x *g*. The supernatant was discarded. The pellets were briefly dried at RT and then dissolved in 50 μl 10 mM Tris-HCl pH 7.5. After measuring DNA concentration using Qubit dsDNA HS Assay Kit (Thermo Fisher Scientific), the 3C library was stored at −80°C.

##### TaqMan qPCR

TaqMan MGB (5’-FAM) probes for the native centromere of the Z chromosome and the neo-centromere on the Z chromosome were designed and synthesized at Thermo Fisher Scientific. For a loading control, a TaqMan MGB (5’-FAM) probe for SMC5 locus on the Z chromosome was also designed and synthesized at Thermo Fisher Scientific. Fifty nanograms of DNA template (3C library) were mixed with TaqMan qPCR reaction mixture: 900 nM constant primer, 900 nM test primer, 250 nM TaqMan probe in 10 μl 1x TaqMan Fast Advanced Master Mix (Thermo Fisher Scientific). qPCR was performed on QuantStadio 3 (Tehrmo Fisher Scientific) with the following cycling condition: 2 min at 95°C, 40 cycles of 1 s at 95°C and 20 s at 60°C. To make a standard curve, a plasmid which contained the ligated chimeric DNA fragment of corresponding to each target site was serially diluted and used for qPCR analysis. All samples were run in triplicate and the mean value of the triplicates was calculated. The slope (*a*) and *y*-intercept (*b*) values were determined from the standard curve for each experiment. The ligation frequency values were calculated as follows: *v*=10^(Ct-*b*)/*a*^. The values were normalized with the loading control value (*v_smc5_*=10^(meanCt-*b*)/*a*^) of each 3C library to obtain the relative ligation frequencies.

## Supplemental Information

The article contains 7 Supplemental Figures and 2 Supplemental Tables.

## Acknowledgments

The authors are very grateful to members of the Fukagawa Lab for fruitful discussion. We also thank R. Fukuoka and K. Oshimo for technical assistance, Y. Hirano for the acquisition of a differential interference contrast microscopy, E. Kakizono and J. Miao for preliminary analyses on this study, to D. Fachinetti for his reagents, Nikon Solutions Co. Ltd., especially K. Tokunaga for STORM imaging. This work was supported by CREST of JST (JPMJCR21E6), JSPS KAKENHI Grant Numbers 17H06167, 20H05389, 21H05752, 22H00408, and 22H04692 to TF, JSPS KAKENHI Grant Numbers 16K18491, and 21H02461 to MH.

## Author contributions

M.H. designed and performed entire experiments in this study. M.A. and S.T. performed structure studies on CENP-C oligomer. R.-S.N., S.S., S.O., I.J.,T.H. performed imaging analyses with SMLM. TF contributed to generation of various DT40 and RPE-1 cell lines. T.F. and M.H. wrote the manuscript, discussing with all authors.

## Supplementary Figure legends

**Figure S1.**
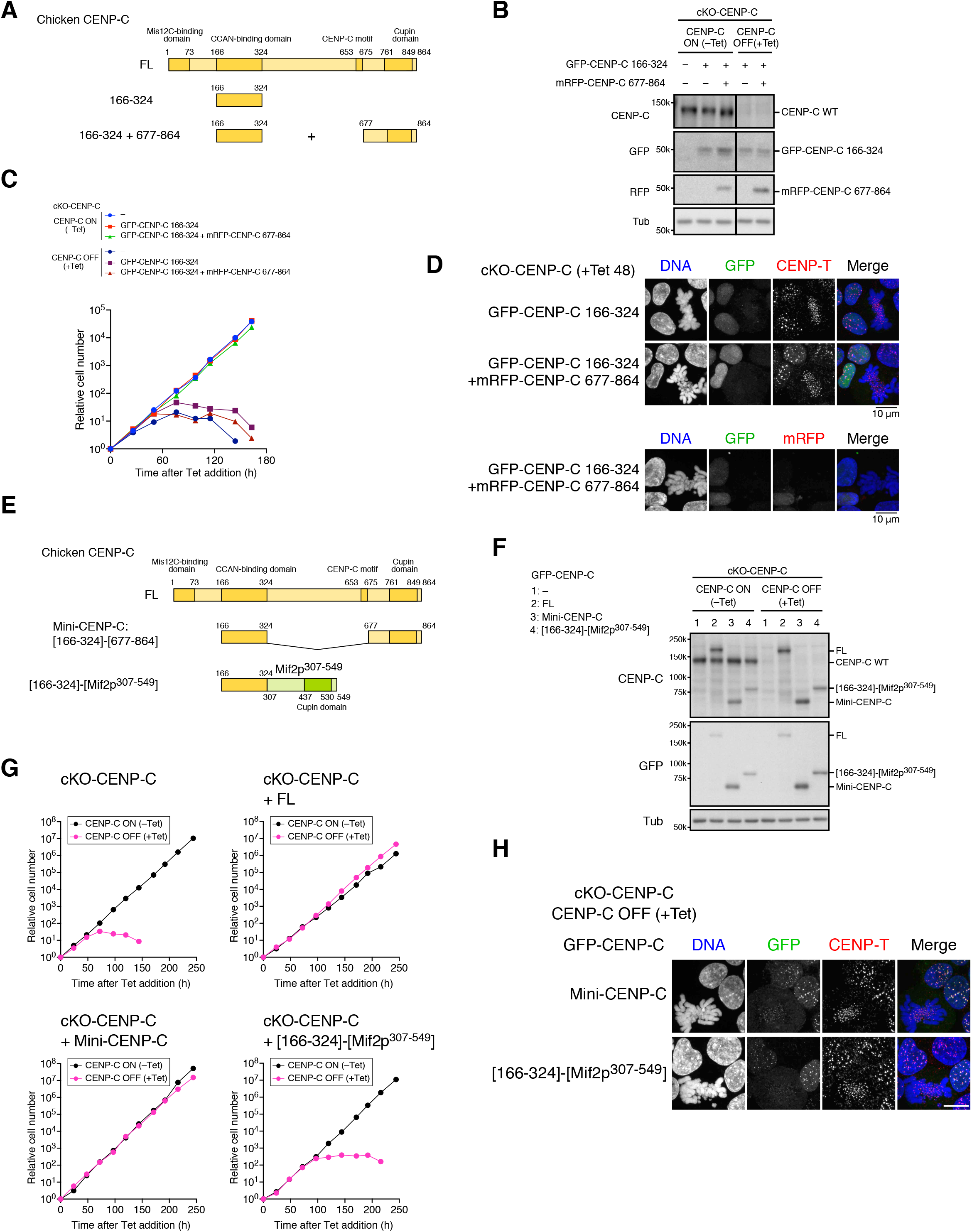
Expression of CCAN-binding and Cupin domains of CENP-C in cKO-CENP-C DT40 cells. Related to Figure 2 and Figure 3. (A) Schematic representation of chicken CENP-C mutants expressed in cKO-CENP-C cells. GFP-fused CCAN-binding domain (aa 166-324) was expressed with or without mRFP-fused CENP-C C-terminus including Cupin domain (677-864) in CENP-C conditional knock-out (cKO-CENP-C) chicken DT40 cells. (B) Protein expression of GFP-CENP-C 166-324 and mRFP-CENP-C 677-864 in cKO-CENP-C cells. GFP-fused CENP-C 166-324 (Δ166-324) was expressed with or without mRFP-CENP-C 677-864 in cKO-CENP-C cells in which wild-type CENP-C (CENP-C WT) was conditionally knocked out by tetracycline (Tet) addition. The cells were cultured with or without Tet (+Tet or –Tet) for 48 h. CENP-C WT protein was examined using an anti-chicken CENP-C antibody. GFP-CENP-C 166-324 and mRFP-CENP-C 677-864 protein were examined using anti-GFP and anti-mRFP antibodies, respectively. The loading control was Alpha-tubulin (Tub). cKO-CENP-C cells without GFP- or mRFP-fused CENP-C expression (–) were also used as a control. (C) Growth of cKO-CENP-C cells expressing GFP-CENP-C 166-324 or both GFP-CENP-C 166-324 and mRFP-CENP-C 677-864. The cell numbers of the indicated cell lines were examined at the indicated times after Tet addition (+Tet) and normalized to those at 0 h for each line. Untreated cells were also examined (–Tet). (D) Localization of GFP-CENP-C 166-324 and mRFP-CENP-C 677-864 in cKO-CENP-C cells. The cells were cultured as described in (B), fixed, and their DNA was stained with DAPI. CENP-T was stained with an anti-chicken CENP-T antibody as a centromeric marker. Scale bar, 10 μm. (E) Schematic representation of the chicken CENP-C mutants expressed in cKO-CENP-C cells. GFP-fused chicken CCAN-binding domain (aa 166-324) was fused with the C-terminus (aa 677-864) containing the Cupin domain of chicken CENP-C ([166-324]-[677-864]: Mini-CENP-C) or with that (Mif2p aa307-549) of *Saccharomyces cerevisiae* Mif2p ([166-324]-[Mif2p307-549]) and expressed in cKO-CENP-C cells. (F) Protein expression of GFP-Mini-CENP-C or GFP-CENP-C 166-324 fused with Mif2p307-549 in cKO-CENP-C cells GFP-Mini-CENP-C or GFP-CENP-C 166-324 fused with Mif2p307-549 was expressed in cKO-CENP-C cells. The cells were cultured with or without Tet (+Tet or –Tet) for 48 h. CENP-C proteins were examined with an anti-chicken CENP-C antibody and an anti-GFP. The loading control was Alpha-tubulin (Tub). cKO-CENP-C cells without GFP-CENP-C expression (−) were also used as a control. (G) Growth of cKO-CENP-C cells expressing GFP-Mini-CENP-C or GFP-CENP-C [166-324]-[Mif2p307-549]. The cell numbers of the indicated cell lines were examined at the indicated times after Tet addition (+Tet) and normalized to those at 0 h for each line. The untreated cells were also examined (–Tet). (H) Localization of GFP-Mini-CENP-C or GFP-CENP-C [166-324]-[Mif2p307-549] in cKO-CENP-C cells. The cells were cultured as described in (B), fixed, and then their DNA was stained with DAPI. CENP-T was stained with an anti-chicken CENP-T antibody as a centromeric marker. Scale bar, 10 μm.

**Figure S2.**
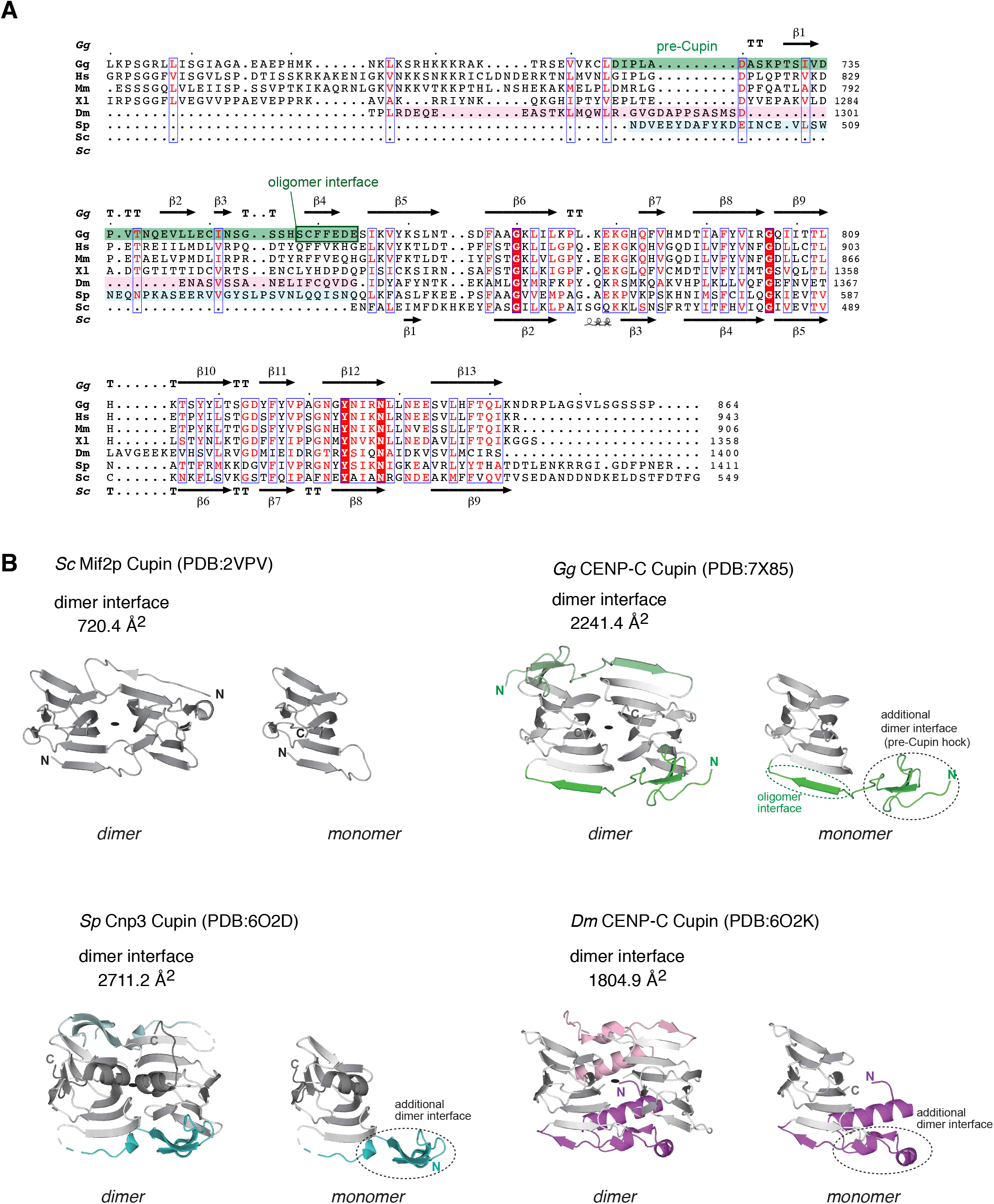
CENP-C Cupin domains. Related to Figure 3. (A) Alignment of amino acid sequence of CENP-C Cupin domains. A structure guided multiple sequence alignment was generated using the PROMALS3D server. The CENP-C Cupin structures (PDB ID 7X85, 6O2D, 6O2K, 2VPV) were used for the alignment. Gg*: Gallus gallus* CENP-C 677-864 (NP_001376225.2), Hs*: Homo sapiens* CENP-C 760-943 (NP_001803), Mm: *Mus musculus* CENP-C 725-906 (NP_031709), Xl: *Xenopus laevis* CENP-C 1228-1400 (NP_0011594), Dm: *Drosophila melanogaster* CENP-C 1270-1411 (NP_731254), Sp: *Schizosaccharomyces pombe* Cnp3 489-643, (NP_001342907), Sc: *Saccharomyces cerevisiae*, MIF2p, 437-549 (UniProtKB P35201). The pre-Cupin regions of Gg CENP-C, Sp CENP-C and Dm CNEP-C are highlighted in green, light pink and light blue, respectively (colored in each model in (B)). (B) Top view of crystal structure of CENP-C Cupin domains. The crystal structure of Cupin domain dimer and monomer of CENP-C orthologs are shown from the top view. The additional dimer interfaces in Chicken (Gg), fission yeast (Sp), and fruit fly (Dm) are colored in green, cyan, and purple, respectively.

**Figure S3.**
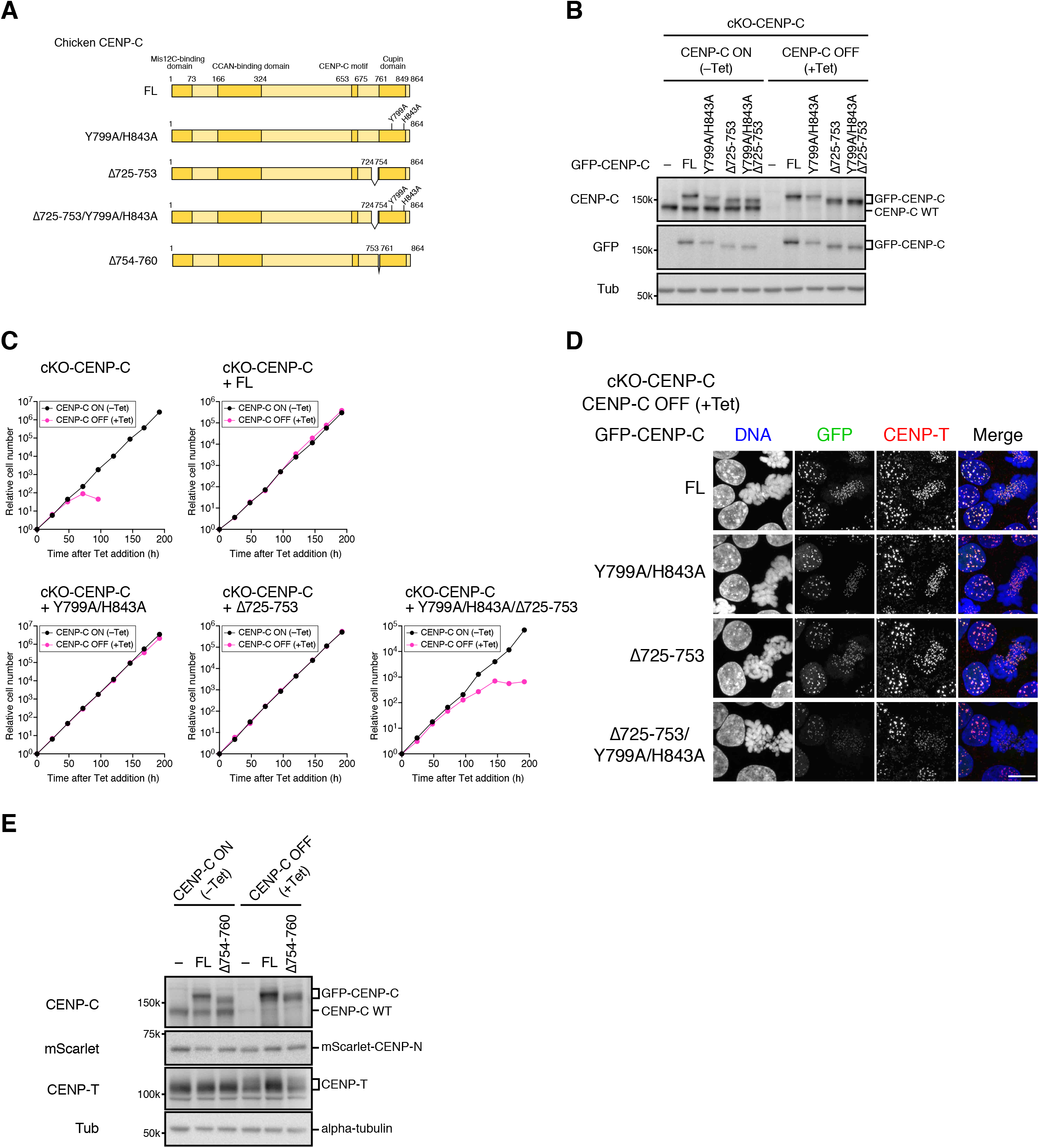
Expression of CENP-C with mutation in the major dimer interface and/or the pre-Cupin, or the oligomerization interface in cKO-CENP-C DT40 cells. Related to Figure 3 and Figure 4. (A) Schematic representation of chicken CENP-C mutants expressed in cKO-CENP-C cells. GFP-fused CENP-C of which two amino acid residues in the major dimer interface of Cupin domain were replaced with alanine (Y799A/H843A). The pre-Cupin region (aa 725-753) was deleted solely or together with Y799A/H844A mutation (Δ725-753, Δ725-753/Y799A/H844A). The oligomerization interface was deleted from GFP-fused CENP-C (Δ754-760). (B) Protein expression of GFP-CENP-C Y799A/H844A, Δ725-753, or Δ725-753/Y799A/H844A in cKO-CENP-C cells. GFP-CENP-C Y799A/H844A, Δ725-753, or Δ725-753/Y799A/H844A was expressed in cKO-CENP-C cells. The cells were cultured with or without Tet (+Tet or –Tet) for 48 h. CENP-C proteins were examined with an anti-chicken CENP-C antibody and an anti-GFP. The loading control was alpha-tubulin (Tub). cKO-CENP-C cells without GFP-CENP-C expression (–) were also examined as a control. (C) Growth of cKO-CENP-C cells expressing GFP-CENP-C Y799A/H844A, Δ725-753, or Δ725-753/Y799A/H844A. The cell numbers of indicated cell lines were examined at the indicated times after Tet addition (+Tet) and were normalized to those at 0 h for each line. Untreated cells were also examined (– Tet). (D) Localization of GFP-CENP-C Y799A/H844A, Δ725-753, or Δ725-753/Y799A/H844A in cKO-CENP-C cells. The cells were cultured as in (B), fixed, and then their DNA was stained with DAPI. CENP-T was also stained with an anti-chicken CENP-T antibody as a centromere marker. Scale bar, 10 μm. (E) GFP-fused CENP-C FL or Δ754-760 was expressed in cKO-CENP-C cells in which endogenous CENP-N was replaced with mScarlet-CENP-N. The cells were cultured with or without Tet (+Tet or –Tet) for 48 h. CENP-C, mScarlet-CENP-N and CENP-T proteins were examined with anti-chicken CENP-C, anti-mRFP and anti-chicken CENP-T antibodies, respectively. The loading control was alpha-tubulin (Tub). cKO-CENP-C cells without GFP-CENP-C expression (–) were examined as a control.

**Figure S4.**
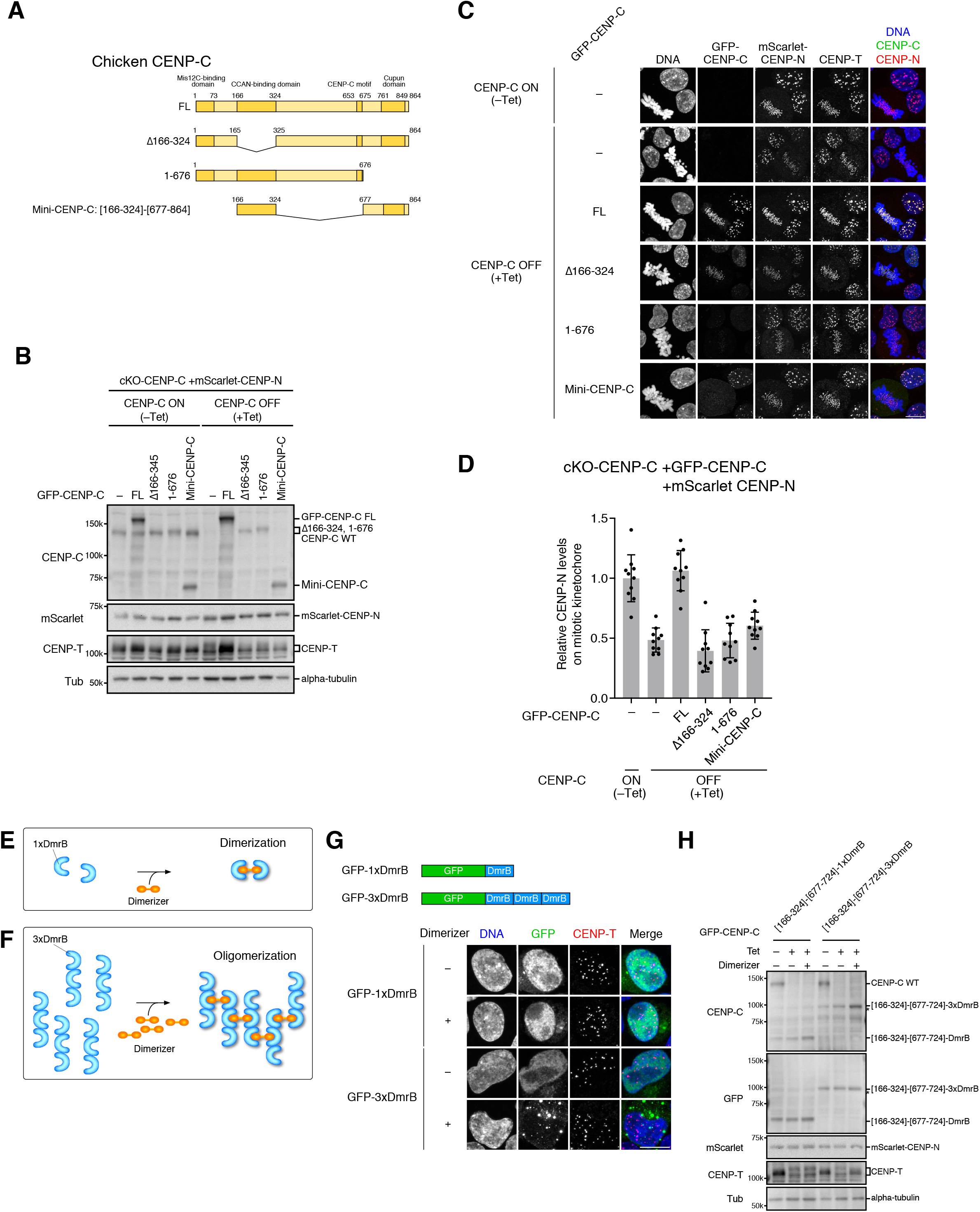
Expression of CENP-C mutants and Mini-CENP-C in cKO-CENP-C DT40 cells, and the artificial oligomerization system. Related to Figure 4 and Figure 5. (A) Schematic representation of chicken CENP-C mutants expressed in cKO-CENP-C cells. The CCAN-binding domain (aa 166-324) is deleted in CENP-C Δ166-324. The C-terminus region including the Cupin domain (aa 677-864) is deleted in CENP-C 1-676. The CCAN-binding domain is fused with the C-terminus including the Cupin domain (Mini-CENP-C: [166-324]-[677-864]). (B) GFP-CENP-C and mScarlet-CENP-N expression in cKO-CENP-C DT40 cells. GFP-fused CENP-C FL Δ166-324, 1-676, or Mini-CENP-C was expressed in cKO-CENP-C cells in which endogenous CENP-N was replaced with mScarlet-CENP-N. The cells were cultured with or without Tet (+Tet or –Tet) for 48 h. CENP-C, mScarlet-CENP-N, and CENP-T proteins were examined with anti-chicken CENP-C, anti-mRFP, and anti-chicken CENP-T antibodies, respectively. The loading control was alpha-tubulin (Tub). cKO-CENP-C cells without GFP-CENP-C expression (–) were examined as a control. (C) Localization of mScarlet-CENP-N in cKO-CENP-C cells expressing GFP-CENP-C mutants. The cells were cultured as in (B), fixed, and stained DNA with DAPI. CENP-T was also stained with an anti-chicken CENP-T antibody as a centromere marker. Scale bar, 10 μm. (D) Quantification of mScarlet-CENP-N signals on mitotic centromeres. mScarlet-CENP-N signals on centromeres in mitotic cells in (C) were quantified. (unpaired t-test, two-tailed, n = 10; – (–Tet) vs. – (+Tet): p<0.0001, – (+Tet) vs. FL (+Tet): p<0.0001, – (+Tet) vs. Δ166-324 (+Tet): ns (p = 0.1730), – (+Tet) vs. 1-676 (+Tet): ns (p = 0.9297), – (+Tet) vs. Mini-CENP-C (+Tet): p = 0.0223). (E) Dimerization of single DmrB (1xDmrB). DmrB (Takara) is an inducible homo-dimerization domain. A small chemical, B/B Homodimerizer (Dimerizer), induces 1xDmrB dimerization. (F) Oligomerization of three tandem-repeated DmrB (3xDmrB). 3xDmrB proteins are oligomerized by Dimerizer (Takamatsu et al., 2013). (G) Oligomerization of GFP-fused 3xDmrB in DT40 cells. GFP fused with 1xDmrB or 3xDmrB was expressed in cKO-CENP-C cells. The cells were treated with or without Dimerizer for 30 min, fixed, and stained DNA with DAPI. CENP-T was also stained with an anti-chicken CENP-T antibody as a centromere marker. Scale bar, 10 μm. (H) Expression of mScarlet-CENP-N and GFP-CENP-C [166-324]-[677-724] fused 1xDmrB or 3xDmrB in cKO-CENP-C DT40 cells. GFP-fused CENP-C [166-324]-[677-724]-1xDmrB C [166-324]-[677-724]-3xDmrB was expressed in cKO-CENP-C cells in which endogenous CENP-N was replaced with mScarlet-CENP-N. The cells were treated with no chemical (–Tet), Tet (+Tet), or both Tet and Dimerizer (+Tet, +Dimerizer) for 48 h. CENP-C, mScarlet-CENP-N, and CENP-T proteins were examined with anti-chicken CENP-C, anti-mRFP, and anti-chicken CENP-T antibodies, respectively. The loading control was alpha-tubulin (Tub). cKO-CENP-C cells without GFP-CENP-C expression (–) were examined as a control.

**Figure S5.**
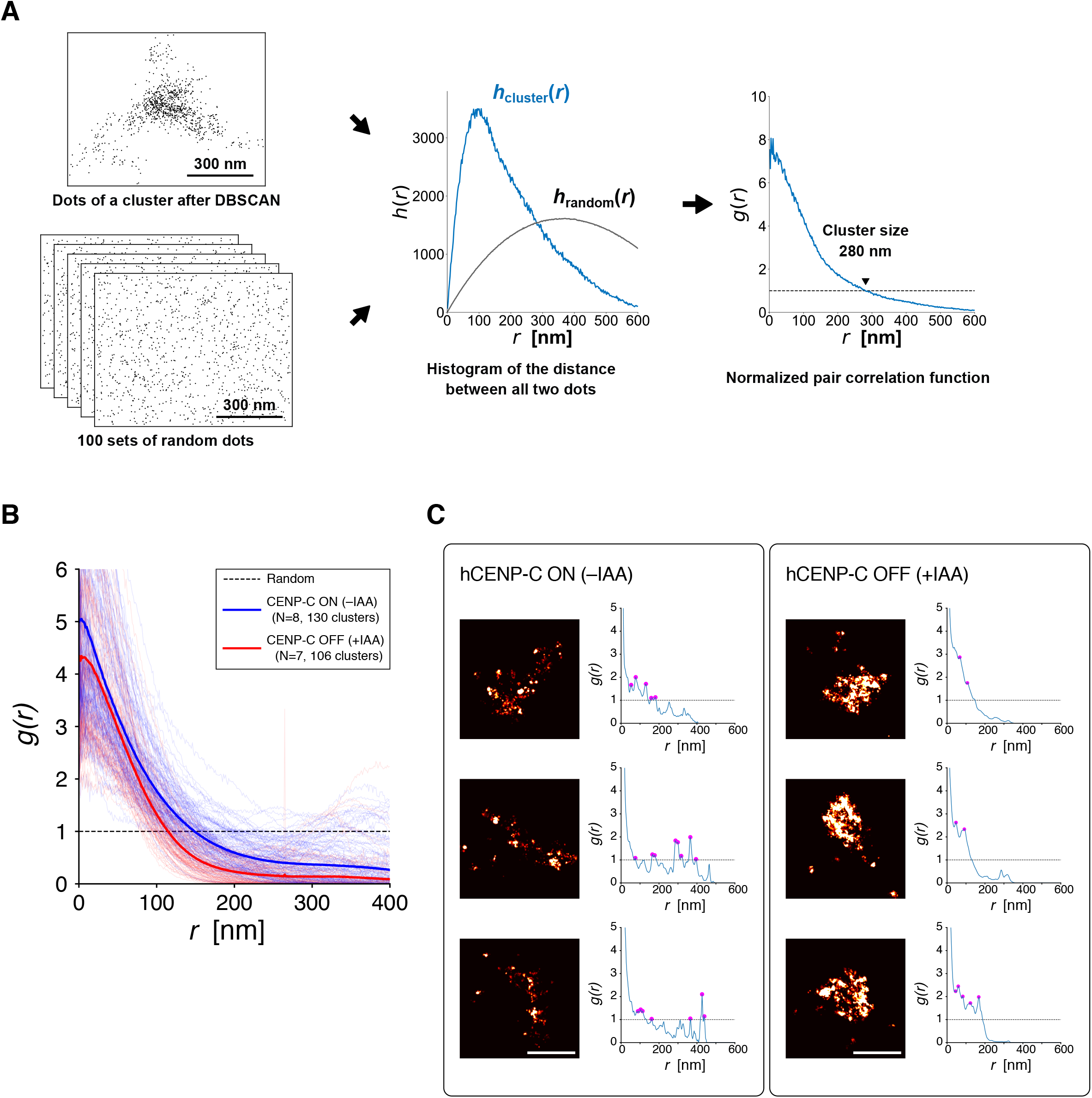
Quantification of CENP-A cluster rendered by SMLM using the pair correlation function. Related to Figure 6. (A) Flowchart of the pair correlation analysis. Based on dots data of a cluster after DBSCAN, we calculated the histograms *h*_cluster_(*r*) and *h*_random_(*r*) with the 2-nm binning width. The normalized pair correlation function of this sample crosses *g*(*r*) = 1 at *r* = 280 nm, and the characteristic length is consistent with the clustering dots of this sample. (B) The pair correlation analysis of CENP-A cluster images of STORM. The thin curves indicate *g*(*r*) of each cluster. The bold blue and red curves show the average value for CENP-C on and off states, respectively. Random distribution gives *g*(*r*) = 1 (dashed line). (C) The pair correlation analysis of CENP-A cluster images of MINFLUX nanoscopy. Three representative images for each condition are shown. Because of high resolution of emitter localization of MINFLUX, the multiple small sub-groups of CENP-A molecules were found within a cluster in RPE-1 hCENP-C-AID cells (-IAA). The distance of sub-groups are depicted as peaks on *g*(*r*) (*g*(*r)*>1) labeled with magenta circles. These peaks are less clear, and their numbers are reduced after IAA treatment (+IAA). The relative frequency of peak appearance at each *r* is shown as the probability density at the distance in Figure 6F.

**Figure S6.**
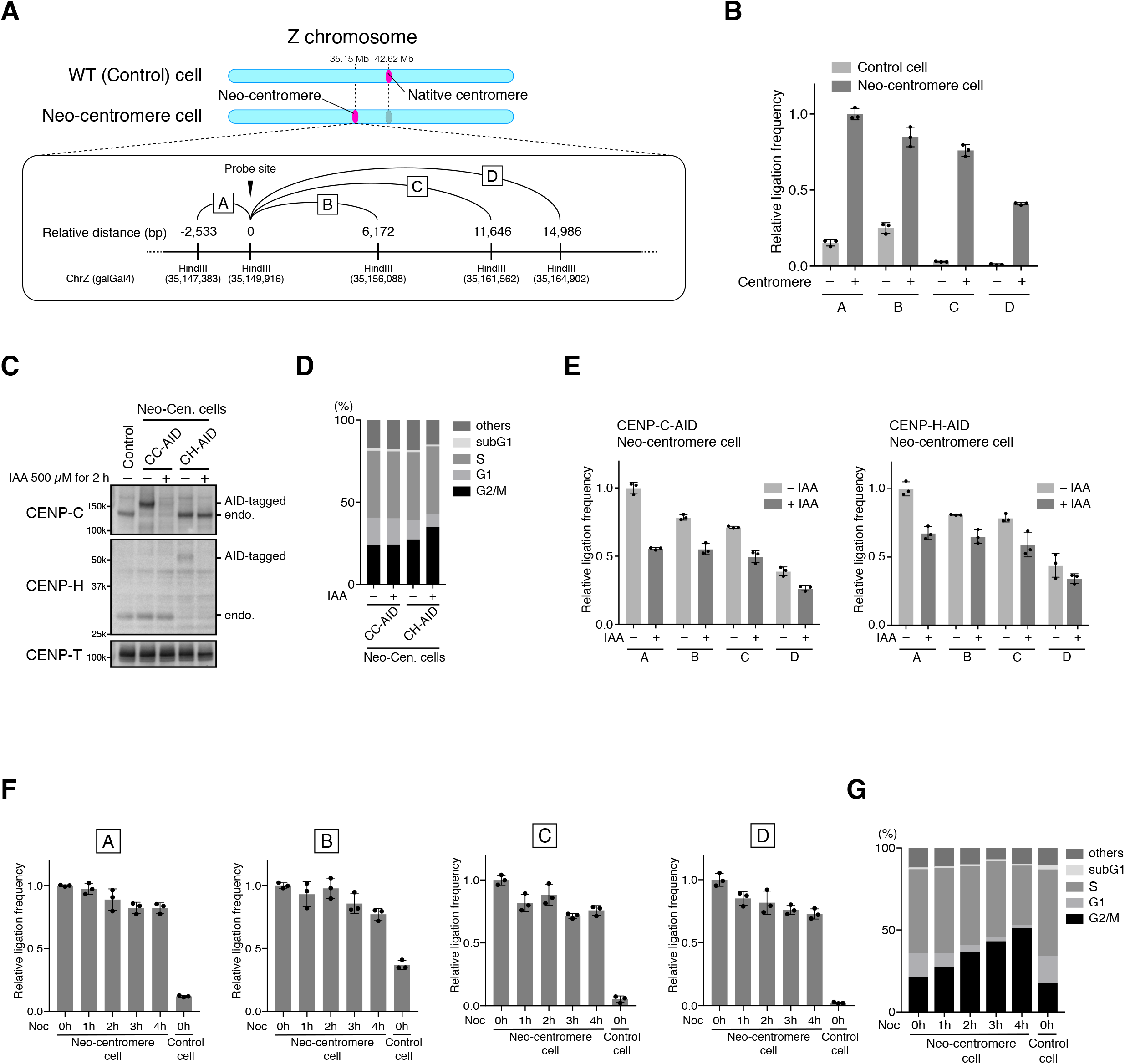
3C-qPCR detects the centromere-specific chromatin structure within neo-centromere. Related to Figure 6. (A) Schematic representation of the chicken CENP-C Z chromosome with a neo-centromere. The chicken Z chromosome in wild-type control DT40 cells (WT, control) has a native centromere spanning 40 kb in the 42.6 Mb region (Hori et al., 2017; Nishimura et al., 2019). A DT40 cell line whose native centromere of the Z chromosome is removed has a neo-centromere at the 35.15 Mb region (Neo-centromere cell) (Nishimura et al., 2019). The bottom inset shows the 3C-qPCR primer and TaqMan probe positions for the neocentromere. The TaqMan probe is designed at 5′ (left side of the diagram) of the HindIII at 0 bp (relative distance, probe site). Primers are designed at the 5′ of the indicated HindIII sites. (B) 3C-qPCR analysis of Neo-centromere cells. The 3C-libraries were prepared from control and Neo-centromere cells, and the ligation frequency between the HindIII sites was examined as indicated in (A) by TaqMan real-time PCR. (C) Conditionally protein knockdown of CENP-C or CENP-H in Neo-centromere cells. Neo-centromere cells (Neo-Cen. Cells) in which CENP-C or CENP-H proteins were conditionally depleted using the auxin-inducible degron (AID) system (CENP-C-AID: CC-AID, CENP-H-AID, CH-AID) were cultured with or without IAA for 2 h. The CENP-C, CENP-H, and CENP-T proteins were examined using anti-chicken CENP-C, anti-chicken CENP-H, and anti-chicken CENP-T antibodies, respectively. CENP-T was used as a loading control. Parental Neo-centromere cells without GFP-CENP-C expression (–) were examined as controls. (D) Cell cycle in neo-centromeric cells expressing CENP-C-AID or CENP-H-AID. Neo-centromere cells expressing CENP-C-AID (CC-AID) or CENP-H-AID (CH-AID) were incubated with BrdU for 20 min at 2 h after IAA addition (+IAA) and harvested. Following fixation, the cells were stained with an anti-BrdU antibody and propidium iodide. Cell cycle distribution was analyzed using flow cytometry. IAA-untreated cells in each line were examined as controls. (E) 3C-qPCR analysis of Neo-centromere cells after knockdown of CENP-C or CENP-H. Neo-centromere cells expressing CENP-C-AID (CC-AID) or CENP-H-AID (CH-AID) were cultured with or without IAA for 2 h. The 3C-libraries were prepared from these cells, and the ligation frequency between the HindIII sites indicated in (A) was examined by TaqMan real-time PCR. (F) 3C-qPCR analysis of Neo-centromere cells treated with nocodazole. Neo-centromere cells were treated with nocodazole (Noc) to enrich mitotic cells for the indicated time periods. The 3C-libraries were prepared from these cells, and the ligation frequency between the HindIII sites indicated in (A) was examined by TaqMan real-time PCR. Untreated control cells were also used as controls. (G) Cell cycle in Neo-centromere cells treated with nocodazole. Control cells and Neo-centromere cells treated with Noc as in (F) were analyzed by flow cytometry as in (D).

**Figure S7.**
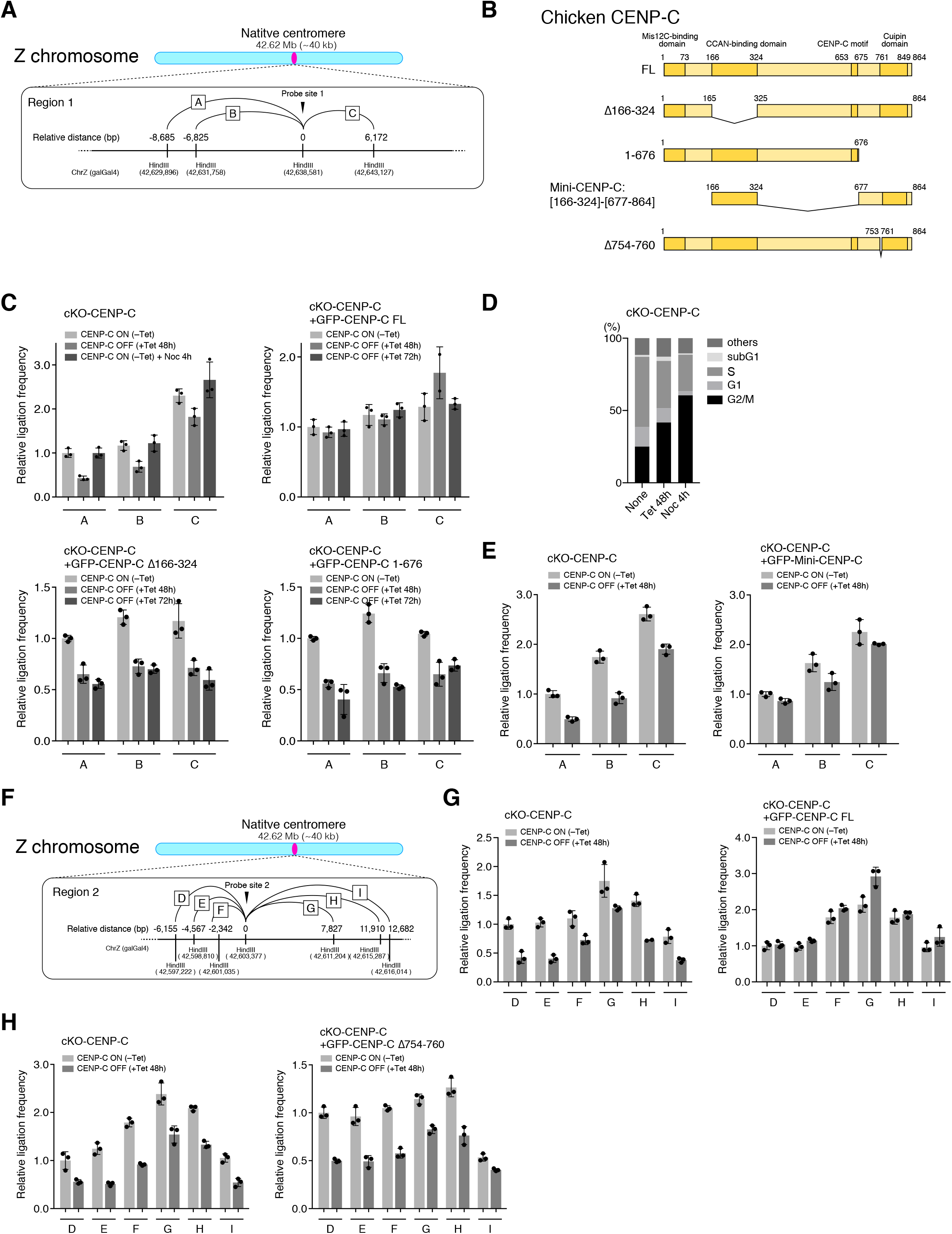
CENP-C is required for chromatin interactions within the native centromere on the Z chromosome in DT40 cells. Related to Figure 6. (A) Schematic representation of the chicken CENP-C Z chromosome. The chicken Z chromosome has the native centromere spanning 40 kb at 42.6 Mb region (Hori et al., 2017; Nishimura et al., 2019). The bottom inset shows 3C-qPCR primer and TaqMan probe positions for the region (region 1) in the native centromere of Z chromosome. The TaqMan probe is designed at 5′ (left side in the diagram) of the HindIII at 0 bp (relative distance, probe site). Primers are designed at 5′ of indicated HindIII sites. (B) Schematic representation of the chicken CENP-C mutant. The CCAN-binding domain (aa 166-324) is deleted in CENP-C Δ166-324. The C-terminus region including Cupin domain (aa 677-864) is deleted in CENP-C 1-676. The CCAN-binding domain is fused with the C-terminus region ([166-324]-[677-864]: Mini-CENP-C). The oligomer interface (aa754-760) is deleted in CENP-C Δ754-760. (C) 3C-qPCR analysis of the native centromere of the Z chromosome in cKO-CENP-C cells with or without GFP-CENP-C expression. cKO-CENP-C cells without GFP-CENP-C expression were cultured with or without Tet for 48 h (+ or – Tet). The cKO-CENP-C (–Tet) were cultured with nocodazole for 4 h (+ Noc). cKO-CENP-C cells expressing GFP-CENP-C FL, Δ166-324, or 1-676 were cultured with or without Tet for 48 h and 72 h. The 3C-libraries were prepared from these cells, and the ligation frequency between the HindIII sites indicated in (A) was examined by TaqMan Real-time PCR. (D) Cell cycle in cKO-CENP-C cells treated with Tet or Noc. cKO-CENP-C cells were incubated with BrdU for 20 min at 48 h after Tet addition (+Tet) or at 4 h after Noc addition (+Noc) and harvested. Following fixation, the cells were stained with an anti-BrdU antibody and propidium iodide. Cell cycle distribution was analyzed by flow cytometry. Untreated cells (Non) were also examined as controls. (E) 3C-qPCR analysis of the native centromere of the Z chromosome in cKO-CENP-C cells expressing GFP-Mini-CENP-C. cKO-CENP-C cells with or without GFP-Mini-CENP-C expression were cultured with or without Tet for 48 h. The 3C-libraries were prepared from these cells, and the ligation frequency between the HindIII sites indicated in (A) was examined by TaqMan Real-time PCR. (F) Schematic representation of the chicken CENP-C Z chromosome. The chicken Z chromosome has the native centromere spanning 40 kb at the 42.6 Mb region (Hori et al., 2017; Nishimura et al., 2019). The bottom inset shows 3C-qPCR primer and TaqMan probe positions for the region (region 2) in native centromere of Z chromosome. The TaqMan probe is designed at 5′ (left side in the diagram) of the HindIII at 0 bp (relative distance, probe site). Primers are designed at 5′ of indicated HindIII sites. (G) 3C-qPCR analysis of the region 2, the native centromere of the Z chromosome, in cKO-CENP-C cells with or without GFP-CENP-C FL expression. cKO-CENP-C cells with or without GFP-CENP-C FL expression were cultured in the presence or absence of Tet for 48 h (+ or – Tet). The 3C-libraries were prepared from these cells, and the ligation frequency between the HindIII sites in the region 2 indicated in (F) was examined by TaqMan Real-time PCR. (H) 3C-qPCR analysis of the native centromere of the Z chromosome in cKO-CENP-C cells expressing GFP-CENP-C Δ754-760. cKO-CENP-C cells with or without GFP-CENP-C Δ754-760 expression were cultured in the presence or absence of Tet for 48 h (+ or – Tet). The 3C-libraries were prepared from these cells, and the ligation frequency between the HindIII sites indicated in (F) was examined by TaqMan Real-time PCR.

**Table S1.**
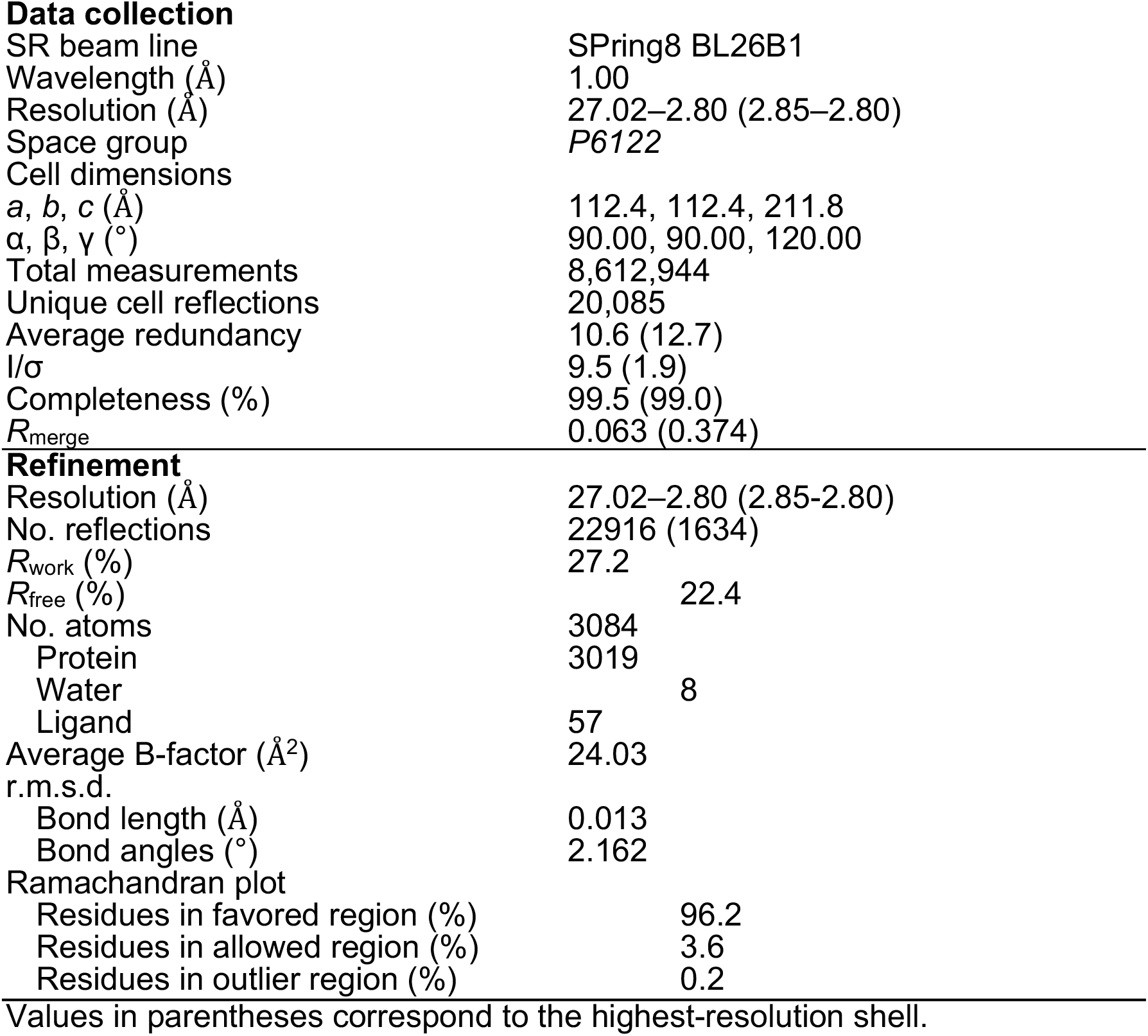
Data collection and refinement statistics.

|  |  |
| --- | --- |
| SR beam line | SPring8 BL26B1 |
| Wavelength (Å) | 1.00 |
| Resolution (Å) | 27.02–2.80 (2.85–2.80) |
| Space group | *P*6122 |
| Cell dimensions |  |
| *a*, *b*, *c* (Å) | 112.4, 112.4, 211.8 |
| $\alpha$, $\beta$, $\gamma$ (°) | 90.00, 90.00, 120.00 |
| Total measurements | 8,612,944 |
| Unique cell reflections | 20,085 |
| Average redundancy | 10.6 (12.7) |
| *I*/$\sigma$ | 9.5 (1.9) |
| Completeness (%) | 99.5 (99.0) |
| *R*<sub>merge</sub> | 0.063 (0.374) |
| **Refinement** |  |
| Resolution (Å) | 27.02–2.80 (2.85–2.80) |
| No. reflections | 22916 (1634) |
| *R*<sub>work</sub> (%) | 27.2 |
| *R*<sub>free</sub> (%) | 22.4 |
| No. atoms | 3084 |
| Protein | 3019 |
| Water | 8 |
| Ligand | 57 |
| Average B-factor (Å<sup>2</sup>) | 24.03 |
| r.m.s.d. |  |
| Bond length (Å) | 0.013 |
| Bond angles (°) | 2.162 |
| Ramachandran plot |  |
| Residues in favored region (%) | 96.2 |
| Residues in allowed region (%) | 3.6 |
| Residues in outlier region (%) | 0.2 |

Values in parentheses correspond to the highest-resolution shell.

**Table S2.**
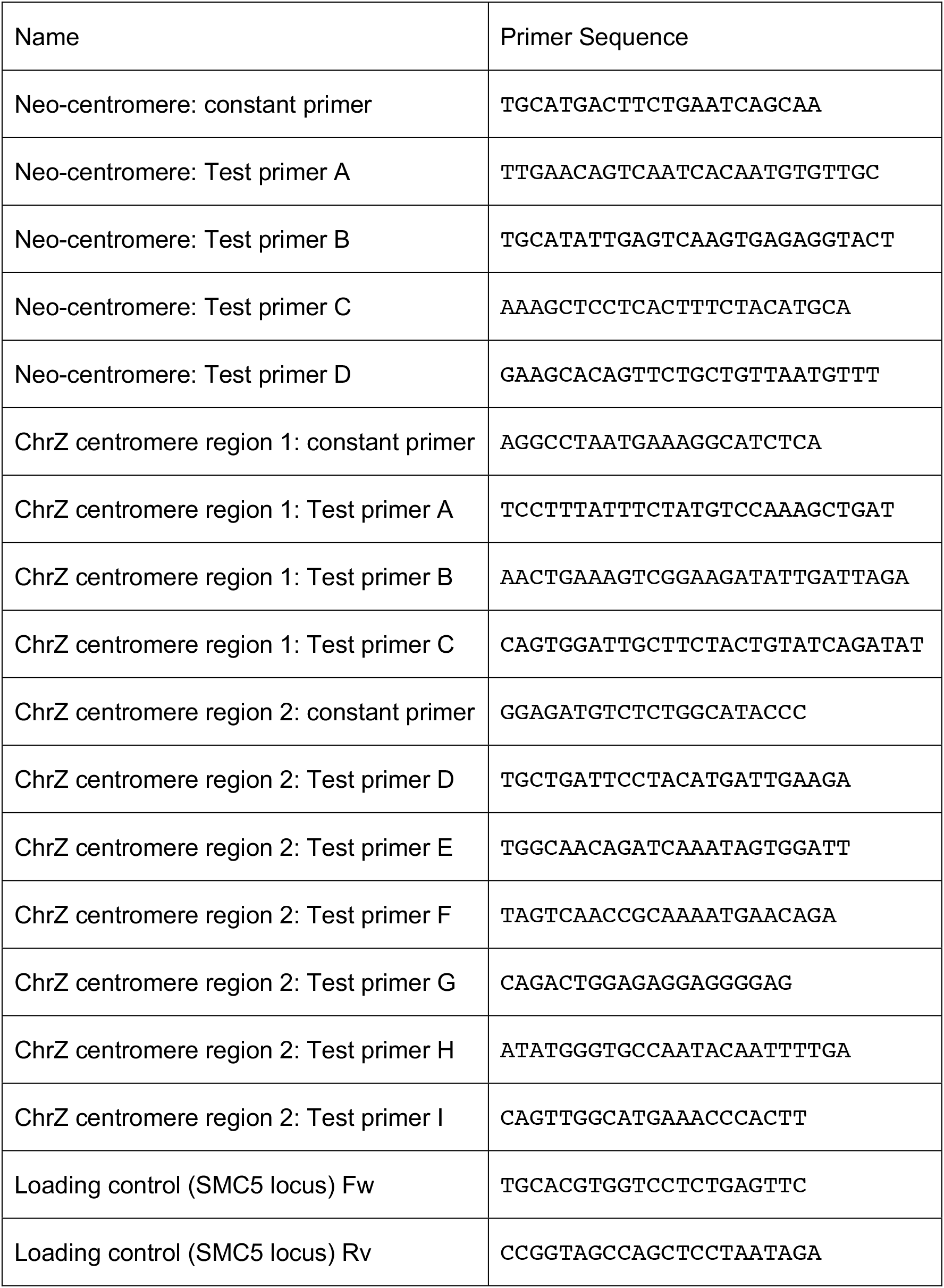
Primers for 3C-qPCR.

## References

Ali-Ahmad, A., Bilokapic, S., Schafer, I.B., Halic, M., and Sekulic, N. (2019). CENP-C unwraps the human CENP-A nucleosome through the H2A C-terminal tail. EMBO Rep 20, e48913.

Allu, P.K., Dawicki-McKenna, J.M., Van Eeuwen, T., Slavin, M., Braitbard, M., Xu, C., Kalisman, N., Murakami, K., and Black, B.E. (2019). Structure of the Human Core Centromeric Nucleosome Complex. Curr Biol 29, 2625–2639 e2625.

Alushin, G.M., Ramey, V.H., Pasqualato, S., Ball, D.A., Grigorieff, N., Musacchio, A., and Nogales, E. (2010). The Ndc80 kinetochore complex forms oligomeric arrays along microtubules. Nature 467, 805–810.

Amano, M., Suzuki, A., Hori, T., Backer, C., Okawa, K., Cheeseman, I.M., and Fukagawa, T. (2009). The CENP-S complex is essential for the stable assembly of outer kinetochore structure. J Cell Biol 186, 173–182.

Ando, S., Yang, H., Nozaki, N., Okazaki, T., and Yoda, K. (2002). CENP-A, -B, and -C chromatin complex that contains the I-type alpha-satellite array constitutes the prekinetochore in HeLa cells. Mol Cell Biol 22, 2229–2241.

Andronov, L., Ouararhni, K., Stoll, I., Klaholz, B.P., and Hamiche, A. (2019). CENP-A nucleosome clusters form rosette-like structures around HJURP during G1. Nat Commun 10, 4436.

Ariyoshi, M., Makino, F., Watanabe, R., Nakagawa, R., Kato, T., Namba, K., Arimura, Y., Fujita, R., Kurumizaka, H., Okumura, E.I., et al. (2021). Cryo-EM structure of the CENP-A nucleosome in complex with phosphorylated CENP-C. EMBO J, e105671.

Balzarotti, F., Eilers, Y., Gwosch, K.C., Gynna, A.H., Westphal, V., Stefani, F.D., Elf, J., and Hell, S.W. (2017). Nanometer resolution imaging and tracking of fluorescent molecules with minimal photon fluxes. Science 355, 606–612.

Basilico, F., Maffini, S., Weir, J.R., Prumbaum, D., Rojas, A.M., Zimniak, T., De Antoni, A., Jeganathan, S., Voss, B., van Gerwen, S., et al. (2014). The pseudo GTPase CENP-M drives human kinetochore assembly. Elife 3, e02978.

Black, B.E., and Cleveland, D.W. (2011). Epigenetic centromere propagation and the nature of CENP-a nucleosomes. Cell 144, 471–479.

Blower, M.D., Sullivan, B.A., and Karpen, G.H. (2002). Conserved organization of centromeric chromatin in flies and humans. Dev Cell 2, 319–330.

Buerstedde, J.M., Reynaud, C.A., Humphries, E.H., Olson, W., Ewert, D.L., and Weill, J.C. (1990). Light chain gene conversion continues at high rate in an ALV-induced cell line. EMBO J 9, 921–927.

Chardon, F., Japaridze, A., Witt, H., Velikovsky, L., Chakraborty, C., Wilhelm, T., Dumont, M., Yang, W., Kikuti, C., Gangnard, S., et al. (2022). CENP-B-mediated DNA loops regulate activity and stability of human centromeres. Mol Cell 82, 1751–1767 e1758.

Cheeseman, I.M., Chappie, J.S., Wilson-Kubalek, E.M., and Desai, A. (2006). The conserved KMN network constitutes the core microtubule-binding site of the kinetochore. Cell 127, 983–997.

Chik, J.K., Moiseeva, V., Goel, P.K., Meinen, B.A., Koldewey, P., An, S., Mellone, B.G., Subramanian, L., and Cho, U.S. (2019). Structures of CENP-C cupin domains at regional centromeres reveal unique patterns of dimerization and recruitment functions for the inner pocket. J Biol Chem 294, 14119–14134.

Clackson, T., Yang, W., Rozamus, L.W., Hatada, M., Amara, J.F., Rollins, C.T., Stevenson, L.F., Magari, S.R., Wood, S.A., Courage, N.L., et al. (1998). Redesigning an FKBP-ligand interface to generate chemical dimerizers with novel specificity. Proc Natl Acad Sci U S A 95, 10437–10442.

Cohen, R.L., Espelin, C.W., De Wulf, P., Sorger, P.K., Harrison, S.C., and Simons, K.T. (2008). Structural and functional dissection of Mif2p, a conserved DNA-binding kinetochore protein. Mol Biol Cell 19, 4480–4491.

Cong, L., Ran, F.A., Cox, D., Lin, S., Barretto, R., Habib, N., Hsu, P.D., Wu, X., Jiang, W., Marraffini, L.A., et al. (2013). Multiplex genome engineering using CRISPR/Cas systems. Science 339, 819–823.

DeLuca, J.G., Gall, W.E., Ciferri, C., Cimini, D., Musacchio, A., and Salmon, E.D. (2006). Kinetochore microtubule dynamics and attachment stability are regulated by Hec1. Cell 127, 969–982.

Dimitrova, Y.N., Jenni, S., Valverde, R., Khin, Y., and Harrison, S.C. (2016). Structure of the MIND Complex Defines a Regulatory Focus for Yeast Kinetochore Assembly. Cell 167, 1014–1027 e1012.

Earnshaw, W.C., and Rothfield, N. (1985). Identification of a family of human centromere proteins using autoimmune sera from patients with scleroderma. Chromosoma 91, 313–321.

Emsley, P., Lohkamp, B., Scott, W.G., and Cowtan, K. (2010). Features and development of Coot. Acta Crystallogr D Biol Crystallogr 66, 486–501.

Fachinetti, D., Folco, H.D., Nechemia-Arbely, Y., Valente, L.P., Nguyen, K., Wong, A.J., Zhu, Q., Holland, A.J., Desai, A., Jansen, L.E., et al. (2013). A two-step mechanism for epigenetic specification of centromere identity and function. Nat Cell Biol 15, 1056–1066.

Falk, S.J., Guo, L.Y., Sekulic, N., Smoak, E.M., Mani, T., Logsdon, G.A., Gupta, K., Jansen, L.E., Van Duyne, G.D., Vinogradov, S.A., et al. (2015). Chromosomes. CENP-C reshapes and stabilizes CENP-A nucleosomes at the centromere. Science 348, 699–703.

Foltz, D.R., Jansen, L.E., Black, B.E., Bailey, A.O., Yates, J.R., 3rd, and Cleveland, D.W. (2006). The human CENP-A centromeric nucleosome-associated complex. Nat Cell Biol 8, 458–469.

Fukagawa, T., and Brown, W.R. (1997). Efficient conditional mutation of the vertebrate CENP-C gene. Hum Mol Genet 6, 2301–2308.

Fukagawa, T., and Earnshaw, W.C. (2014). The Centromere: Chromatin Foundation for the Kinetochore Machinery. Dev Cell 30, 496–508.

Fukagawa, T., Mikami, Y., Nishihashi, A., Regnier, V., Haraguchi, T., Hiraoka, Y., Sugata, N., Todokoro, K., Brown, W., and Ikemura, T. (2001). CENP-H, a constitutive centromere component, is required for centromere targeting of CENP-C in vertebrate cells. EMBO J 20, 4603–4617.

Fukagawa, T., Pendon, C., Morris, J., and Brown, W. (1999). CENP-C is necessary but not sufficient to induce formation of a functional centromere. EMBO J 18, 4196–4209.

Gallego, J., Chou, S.H., and Reid, B.R. (1997). Centromeric pyrimidine strands fold into an intercalated motif by forming a double hairpin with a novel T:G:G:T tetrad: solution structure of the d(TCCCGTTTCCA) dimer. J Mol Biol 273, 840–856.

Garavis, M., Mendez-Lago, M., Gabelica, V., Whitehead, S.L., Gonzalez, C., and Villasante, A. (2015). The structure of an endogenous Drosophila centromere reveals the prevalence of tandemly repeated sequences able to form i-motifs. Sci Rep 5, 13307.

Giunta, S., and Funabiki, H. (2017). Integrity of the human centromere DNA repeats is protected by CENP-A, CENP-C, and CENP-T. Proc Natl Acad Sci U S A 114, 1928–1933.

Guo, L.Y., Allu, P.K., Zandarashvili, L., McKinley, K.L., Sekulic, N., Dawicki-McKenna, J.M., Fachinetti, D., Logsdon, G.A., Jamiolkowski, R.M., Cleveland, D.W., et al. (2017). Centromeres are maintained by fastening CENP-A to DNA and directing an arginine anchor-dependent nucleosome transition. Nat Commun 8, 15775.

Hagege, H., Klous, P., Braem, C., Splinter, E., Dekker, J., Cathala, G., de Laat, W., and Forne, T. (2007). Quantitative analysis of chromosome conformation capture assays (3C-qPCR). Nat Protoc 2, 1722–1733.

Hara, M., Ariyoshi, M., Okumura, E.I., Hori, T., and Fukagawa, T. (2018). Multiple phosphorylations control recruitment of the KMN network onto kinetochores. Nat Cell Biol 20, 1378–1388.

Hara, M., and Fukagawa, T. (2017). Critical Foundation of the Kinetochore: The Constitutive Centromere-Associated Network (CCAN). Prog Mol Subcell Biol 56, 29–57.

Hara, M., and Fukagawa, T. (2018). Kinetochore assembly and disassembly during mitotic entry and exit. Curr Opin Cell Biol 52, 73–81.

Harris, C.R., Millman, K.J., van der Walt, S.J., Gommers, R., Virtanen, P., Cournapeau, D., Wieser, E., Taylor, J., Berg, S., Smith, N.J., et al. (2020). Array programming with NumPy. Nature 585, 357–362.

Hinshaw, S.M., and Harrison, S.C. (2019). The structure of the Ctf19c/CCAN from budding yeast. Elife 8.

Hori, T., Amano, M., Suzuki, A., Backer, C.B., Welburn, J.P., Dong, Y., McEwen, B.F., Shang, W.H., Suzuki, E., Okawa, K., et al. (2008). CCAN makes multiple contacts with centromeric DNA to provide distinct pathways to the outer kinetochore. Cell 135, 1039–1052.

Hori, T., Kagawa, N., Toyoda, A., Fujiyama, A., Misu, S., Monma, N., Makino, F., Ikeo, K., and Fukagawa, T. (2017). Constitutive centromere-associated network controls centromere drift in vertebrate cells. J Cell Biol 216, 101–113.

Hunter, J.D. (2007). Matplotlib: A 2D graphics environment. Comput Sci Eng 9, 90–95.

Izuta, H., Ikeno, M., Suzuki, N., Tomonaga, T., Nozaki, N., Obuse, C., Kisu, Y., Goshima, N., Nomura, F., Nomura, N., et al. (2006). Comprehensive analysis of the ICEN (Interphase Centromere Complex) components enriched in the CENP-A chromatin of human cells. Genes Cells 11, 673–684.

Kabsch, W. (2010). Xds. Acta Crystallogr D Biol Crystallogr 66, 125–132.

Kalitsis, P., Fowler, K.J., Earle, E., Hill, J., and Choo, K.H. (1998). Targeted disruption of mouse centromere protein C gene leads to mitotic disarray and early embryo death. Proc Natl Acad Sci U S A 95, 1136–1141.

Kato, H., Jiang, J., Zhou, B.R., Rozendaal, M., Feng, H., Ghirlando, R., Xiao, T.S., Straight, A.F., and Bai, Y. (2013). A conserved mechanism for centromeric nucleosome recognition by centromere protein CENP-C. Science 340, 1110–1113.

Kim, J., Ishiguro, K., Nambu, A., Akiyoshi, B., Yokobayashi, S., Kagami, A., Ishiguro, T., Pendas, A.M., Takeda, N., Sakakibara, Y., et al. (2015). Meikin is a conserved regulator of meiosis-I-specific kinetochore function. Nature 517, 466–471.

Klare, K., Weir, J.R., Basilico, F., Zimniak, T., Massimiliano, L., Ludwigs, N., Herzog, F., and Musacchio, A. (2015). CENP-C is a blueprint for constitutive centromere-associated network assembly within human kinetochores. J Cell Biol 210, 11–22.

Kwon, M.S., Hori, T., Okada, M., and Fukagawa, T. (2007). CENP-C is involved in chromosome segregation, mitotic checkpoint function, and kinetochore assembly. Mol Biol Cell 18, 2155–2168.

Larson, A.G., Elnatan, D., Keenen, M.M., Trnka, M.J., Johnston, J.B., Burlingame, A.L., Agard, D.A., Redding, S., and Narlikar, G.J. (2017). Liquid droplet formation by HP1alpha suggests a role for phase separation in heterochromatin. Nature 547, 236–240.

Lin, Y., Mori, E., Kato, M., Xiang, S., Wu, L., Kwon, I., and McKnight, S.L. (2016). Toxic PR Poly-Dipeptides Encoded by the C9orf72 Repeat Expansion Target LC Domain Polymers. Cell 167, 789–802 e712.

Maier, N.K., Ma, J., Lampson, M.A., and Cheeseman, I.M. (2021). Separase cleaves the kinetochore protein Meikin at the meiosis I/II transition. Dev Cell 56, 2192–2206 e2198.

Marshall, O.J., Marshall, A.T., and Choo, K.H. (2008). Three-dimensional localization of CENP-A suggests a complex higher order structure of centromeric chromatin. J Cell Biol 183, 1193–1202.

Mates, L., Chuah, M.K., Belay, E., Jerchow, B., Manoj, N., Acosta-Sanchez, A., Grzela, D.P., Schmitt, A., Becker, K., Matrai, J., et al. (2009). Molecular evolution of a novel hyperactive Sleeping Beauty transposase enables robust stable gene transfer in vertebrates. Nat Genet 41, 753–761.

McKinley, K.L., and Cheeseman, I.M. (2016). The molecular basis for centromere identity and function. Nat Rev Mol Cell Biol 17, 16–29.

McKinley, K.L., Sekulic, N., Guo, L.Y., Tsinman, T., Black, B.E., and Cheeseman, I.M. (2015). The CENP-L-N Complex Forms a Critical Node in an Integrated Meshwork of Interactions at the Centromere-Kinetochore Interface. Mol Cell 60, 886–898.

Medina-Pritchard, B., Lazou, V., Zou, J., Byron, O., Abad, M.A., Rappsilber, J., Heun, P., and Jeyaprakash, A.A. (2020). Structural basis for centromere maintenance by Drosophila CENP-A chaperone CAL1. EMBO J 39, e103234.

Mellone, B.G., and Fachinetti, D. (2021). Diverse mechanisms of centromere specification. Curr Biol 31, R1491–R1504.

Milks, K.J., Moree, B., and Straight, A.F. (2009). Dissection of CENP-C-directed centromere and kinetochore assembly. Mol Biol Cell 20, 4246–4255.

Murshudov, G.N., Vagin, A.A., Lebedev, A., Wilson, K.S., and Dodson, E.J. (1999). Efficient anisotropic refinement of macromolecular structures using FFT. Acta Crystallogr D Biol Crystallogr 55, 247–255.

Nagpal, H., and Fukagawa, T. (2016). Kinetochore assembly and function through the cell cycle. Chromosoma 125, 645–659.

Nagpal, H., Hori, T., Furukawa, A., Sugase, K., Osakabe, A., Kurumizaka, H., and Fukagawa, T. (2015). Dynamic changes in CCAN organization through CENP-C during cell-cycle progression. Mol Biol Cell 26, 3768–3776.

Nechemia-Arbely, Y., Miga, K.H., Shoshani, O., Aslanian, A., McMahon, M.A., Lee, A.Y., Fachinetti, D., Yates, J.R., 3rd, Ren, B., and Cleveland, D.W. (2019). DNA replication acts as an error correction mechanism to maintain centromere identity by restricting CENP-A to centromeres. Nat Cell Biol 21, 743–754.

Nishimura, K., Komiya, M., Hori, T., Itoh, T., and Fukagawa, T. (2019). 3D genomic architecture reveals that neocentromeres associate with heterochromatin regions. J Cell Biol 218, 134–149.

Nishino, T., Takeuchi, K., Gascoigne, K.E., Suzuki, A., Hori, T., Oyama, T., Morikawa, K., Cheeseman, I.M., and Fukagawa, T. (2012). CENP-T-W-S-X forms a unique centromeric chromatin structure with a histone-like fold. Cell 148, 487–501.

Okada, M., Cheeseman, I.M., Hori, T., Okawa, K., McLeod, I.X., Yates, J.R., 3rd, Desai, A., and Fukagawa, T. (2006). The CENP-H-I complex is required for the efficient incorporation of newly synthesized CENP-A into centromeres. Nat Cell Biol 8, 446–457.

Palmer, D.K., O’Day, K., Wener, M.H., Andrews, B.S., and Margolis, R.L. (1987). A 17-kD centromere protein (CENP-A) copurifies with nucleosome core particles and with histones. J Cell Biol 104, 805–815.

Pedregosa, F., Varoquaux, G., Gramfort, A., Michel, V., Thirion, B., Grisel, O., Blondel, M., Prettenhofer, P., Weiss, R., Dubourg, V., et al. (2011). Scikit-learn: Machine Learning in Python. J Mach Learn Res 12, 2825–2830.

Pei, J., Tang, M., and Grishin, N.V. (2008). PROMALS3D web server for accurate multiple protein sequence and structure alignments. Nucleic Acids Res 36, W30–34.

Pesenti, M.E., Prumbaum, D., Auckland, P., Smith, C.M., Faesen, A.C., Petrovic, A., Erent, M., Maffini, S., Pentakota, S., Weir, J.R., et al. (2018). Reconstitution of a 26-Subunit Human Kinetochore Reveals Cooperative Microtubule Binding by CENP-OPQUR and NDC80. Mol Cell 71, 923–939 e910.

Pesenti, M.E., Raisch, T., Conti, D., Walstein, K., Hoffmann, I., Vogt, D., Prumbaum, D., Vetter, I.R., Raunser, S., and Musacchio, A. (2022). Structure of the human inner kinetochore CCAN complex and its significance for human centromere organization. Mol Cell.

Pesenti, M.E., Weir, J.R., and Musacchio, A. (2016). Progress in the structural and functional characterization of kinetochores. Curr Opin Struct Biol 37, 152–163.

Petrovic, A., Keller, J., Liu, Y., Overlack, K., John, J., Dimitrova, Y.N., Jenni, S., van Gerwen, S., Stege, P., Wohlgemuth, S., et al. (2016). Structure of the MIS12 Complex and Molecular Basis of Its Interaction with CENP-C at Human Kinetochores. Cell 167, 1028–1040 e1015.

Petrovic, A., Pasqualato, S., Dube, P., Krenn, V., Santaguida, S., Cittaro, D., Monzani, S., Massimiliano, L., Keller, J., Tarricone, A., et al. (2010). The MIS12 complex is a protein interaction hub for outer kinetochore assembly. J Cell Biol 190, 835–852.

Przewloka, M.R., Venkei, Z., Bolanos-Garcia, V.M., Debski, J., Dadlez, M., and Glover, D.M. (2011). CENP-C is a structural platform for kinetochore assembly. Curr Biol 21, 399–405.

Ribeiro, S.A., Vagnarelli, P., Dong, Y., Hori, T., McEwen, B.F., Fukagawa, T., Flors, C., and Earnshaw, W.C. (2010). A super-resolution map of the vertebrate kinetochore. Proc Natl Acad Sci U S A 107, 10484–10489.

Schindelin, J., Arganda-Carreras, I., Frise, E., Kaynig, V., Longair, M., Pietzsch, T., Preibisch, S., Rueden, C., Saalfeld, S., Schmid, B., et al. (2012). Fiji: an open-source platform for biological-image analysis. Nat Methods 9, 676–682.

Schmidt, R., Weihs, T., Wurm, C.A., Jansen, I., Rehman, J., Sahl, S.J., and Hell, S.W. (2021). MINFLUX nanometer-scale 3D imaging and microsecond-range tracking on a common fluorescence microscope. Nat Commun 12, 1478.

Shang, W.H., Hori, T., Martins, N.M., Toyoda, A., Misu, S., Monma, N., Hiratani, I., Maeshima, K., Ikeo, K., Fujiyama, A., et al. (2013). Chromosome engineering allows the efficient isolation of vertebrate neocentromeres. Dev Cell 24, 635–648.

Shang, W.H., Hori, T., Toyoda, A., Kato, J., Popendorf, K., Sakakibara, Y., Fujiyama, A., and Fukagawa, T. (2010). Chickens possess centromeres with both extended tandem repeats and short non-tandem-repetitive sequences. Genome Res 20, 1219–1228.

Sugimoto, K., Kuriyama, K., Shibata, A., and Himeno, M. (1997). Characterization of internal DNA-binding and C-terminal dimerization domains of human centromere/kinetochore autoantigen CENP-C in vitro: role of DNA-binding and self-associating activities in kinetochore organization. Chromosome Res 5, 132–141.

Sullivan, B.A., Blower, M.D., and Karpen, G.H. (2001). Determining centromere identity: cyclical stories and forking paths. Nat Rev Genet 2, 584–596.

Sullivan, B.A., and Karpen, G.H. (2004). Centromeric chromatin exhibits a histone modification pattern that is distinct from both euchromatin and heterochromatin. Nat Struct Mol Biol 11, 1076–1083.

Suzuki, A., Badger, B.L., Wan, X., DeLuca, J.G., and Salmon, E.D. (2014). The architecture of CCAN proteins creates a structural integrity to resist spindle forces and achieve proper Intrakinetochore stretch. Dev Cell 30, 717–730.

Suzuki, A., Hori, T., Nishino, T., Usukura, J., Miyagi, A., Morikawa, K., and Fukagawa, T. (2011). Spindle microtubules generate tension-dependent changes in the distribution of inner kinetochore proteins. J Cell Biol 193, 125–140.

Takamatsu, S., Onoguchi, K., Onomoto, K., Narita, R., Takahasi, K., Ishidate, F., Fujiwara, T.K., Yoneyama, M., Kato, H., and Fujita, T. (2013). Functional characterization of domains of IPS-1 using an inducible oligomerization system. PLoS One 8, e53578.

Tomkiel, J., Cooke, C.A., Saitoh, H., Bernat, R.L., and Earnshaw, W.C. (1994). CENP-C is required for maintaining proper kinetochore size and for a timely transition to anaphase. J Cell Biol 125, 531–545.

Touw, W.G., Baakman, C., Black, J., te Beek, T.A., Krieger, E., Joosten, R.P., and Vriend, G. (2015). A series of PDB-related databanks for everyday needs. Nucleic Acids Res 43, D364–368.

Trazzi, S., Perini, G., Bernardoni, R., Zoli, M., Reese, J.C., Musacchio, A., and Della Valle, G. (2009). The C-terminal domain of CENP-C displays multiple and critical functions for mammalian centromere formation. PLoS One 4, e5832.

Vagin, A., and Teplyakov, A. (2010). Molecular replacement with MOLREP. Acta Crystallogr D Biol Crystallogr 66, 22–25.

Virtanen, P., Gommers, R., Oliphant, T.E., Haberland, M., Reddy, T., Cournapeau, D., Burovski, E., Peterson, P., Weckesser, W., Bright, J., et al. (2020). SciPy 1.0: fundamental algorithms for scientific computing in Python. Nat Methods 17, 261–272.

Walstein, K., Petrovic, A., Pan, D., Hagemeier, B., Vogt, D., Vetter, I.R., and Musacchio, A. (2021). Assembly principles and stoichiometry of a complete human kinetochore module. Sci Adv 7.

Watanabe, R., Hara, M., Okumura, E.I., Herve, S., Fachinetti, D., Ariyoshi, M., and Fukagawa, T. (2019). CDK1-mediated CENP-C phosphorylation modulates CENP-A binding and mitotic kinetochore localization. J Cell Biol 218, 4042–4062.

Watanabe, R., Hirano, Y., Hara, M., Hiraoka, Y., and Fukagawa, T. (2022). Mobility of kinetochore proteins measured by FRAP analysis in living cells. Chromosome Res 30, 43–57.

Weir, J.R., Faesen, A.C., Klare, K., Petrovic, A., Basilico, F., Fischbock, J., Pentakota, S., Keller, J., Pesenti, M.E., Pan, D., et al. (2016). Insights from biochemical reconstitution into the architecture of human kinetochores. Nature 537, 249–253.

Westhorpe, F.G., and Straight, A.F. (2013). Functions of the centromere and kinetochore in chromosome segregation. Curr Opin Cell Biol 25, 334–340.

Winn, M.D., Ballard, C.C., Cowtan, K.D., Dodson, E.J., Emsley, P., Evans, P.R., Keegan, R.M., Krissinel, E.B., Leslie, A.G., McCoy, A., et al. (2011). Overview of the CCP4 suite and current developments. Acta Crystallogr D Biol Crystallogr 67, 235–242.

Yan, K., Yang, J., Zhang, Z., McLaughlin, S.H., Chang, L., Fasci, D., Ehrenhofer-Murray, A.E., Heck, A.J.R., and Barford, D. (2019). Structure of the inner kinetochore CCAN complex assembled onto a centromeric nucleosome. Nature 574, 278–282.

Yatskevich, S., Muir, K.W., Bellini, D., Zhang, Z., Yang, J., Tischer, T., Predin, M., Dendooven, T., McLaughlin, S.H., and Barford, D. (2022). Structure of the human inner kinetochore bound to a centromeric CENP-A nucleosome. Science, eabn3810.

Zasadzinska, E., Huang, J., Bailey, A.O., Guo, L.Y., Lee, N.S., Srivastava, S., Wong, K.A., French, B.T., Black, B.E., and Foltz, D.R. (2018). Inheritance of CENP-A Nucleosomes during DNA Replication Requires HJURP. Dev Cell 47, 348–362 e347.

Zhou, K., Gebala, M., Woods, D., Sundararajan, K., Edwards, G., Krzizike, D., Wereszczynski, J., Straight, A.F., and Luger, K. (2022). CENP-N promotes the compaction of centromeric chromatin. Nat Struct Mol Biol 29, 403–413.

